# Pathobiont-triggered induction of epithelial IDO1 drives regional susceptibility to Inflammatory Bowel Disease

**DOI:** 10.1101/2025.01.04.630951

**Authors:** Paige N. Spencer, Jiawei Wang, Erin P. Smith, Luisella Spiga, Alan J. Simmons, Taewoo Kim, William Kim, Monica E. Brown, Yilin Yang, Harsimran Kaur, Yanwen Xu, Seung Woo Kang, Matthew D. Helou, Mason A. Lee, Lin Zheng, Deronisha Arceneaux, Naila Tasneem, Katherine D. Mueller, Ozge S. Kuddar, Mariah H. Harned, James Ro, Jing Li, Amrita Banerjee, Nicholas O. Markham, Keith T. Wilson, Lori A. Coburn, Jeremy A. Goettel, Qi Liu, M. Kay Washington, Raphael H. Valdivia, Wenhan Zhu, Ken S. Lau

## Abstract

The structure and function of the mammalian gut vary by region, yet why inflammatory diseases manifest in specific regions and not others remains unclear. We use a TNF-overexpressing Crohn’s disease (CD) model (Tnf^ΔARE/+^), which typically presents in the terminal ileum (TI), to investigate how environmental factors interact with the host’s immune susceptibility to drive region-specific disease. We identified *Chlamydia muridarum,* an intracellular bacterium and murine counterpart to the human sexually transmitted *C. trachomatis*, as necessary and sufficient to trigger disease manifestation in the ascending colon (AC), another common site of human CD. Disease manifestation in the AC depends on indoleamine 2,3-dioxygenase (IDO1) activity induced by hypersensitive surface secretory cells in genetically susceptible hosts. Single-cell and microbial analyses of human specimens also implicates this pathobiont–epithelial IDO1 pathway in patients with a history of CD in the AC. Our findings demonstrate that genetic and microbial factors can independently drive region-specific disease and provide a unique model to study CD specific to the AC.

## INTRODUCTION

Organ regionalization plays important roles in organ function and in shaping distinct disease patterns. With advanced, high-resolution tools, recent research has shown that each section of the intestine has distinct characteristics with specialized homeostatic functions and responses to microenvironmental cues.^1,2^ The ileum, with a moderate microbial load, contains specialized Paneth cells that produce antimicrobial peptides to mitigate microbial insults while performing essential functions of micronutrient and macronutrient absorption.^3^ In the downstream colon, microbial load increases significantly, accompanied by shifts in absorption that are focused primarily on water, electrolytes, and short-chain fatty acids produced by microbial fermentation. Moreover, the ileum, proximal colon, and distal colon differ in their epithelial structure and mucous composition.^4–8^ These regional differences in the intestine have important implications for disease development.

Inflammatory bowel disease (IBD), including Crohn’s disease (CD), encompasses a wide range of pathological and symptomological characteristics and has an incompletely understood, multifactorial etiology.^9–15^ While CD can affect any site along the gastrointestinal (GI) tract, the majority of patients present with inflammation in their distal small intestine (terminal ileum, TI) and/or proximal large intestine (ascending colon, AC).^16–20^ The factors contributing to the development of disease in these distinct anatomical sites remain elusive, and current therapeutic management of CD is not fully tailored to the site of involvement, especially for different affected regions of the colon. For instance, biologics and modern small molecule therapies, currently approved to target various cytokines, integrins, and immune cell signaling pathways, are not tailored to ileal nor colonic regional involvement.

The manifestation of CD in different regions of the intestine is highly complex, driven by a multifactorial interplay between genetics, immune responses, and environmental factors. Genome-wide association studies have identified over 240 susceptibility loci for IBD, highlighting genes involved in immune function^10,12,21^ and those that regulate chemical and physical barriers lining the gut lumen.^22,23^ With the exception of very early onset IBD which can be monogenic, host susceptibility alone is insufficient to cause disease, as mouse models emphasize the critical role of the microbiome.^24–26^ Conversely, certain microbes, known as pathobionts, do not trigger disease unless present in predisposed individuals.^27^ Distinct microbiomes across gut regions, coupled with cell-type-specific effects of host susceptibility, underscore the complexity of inflammatory disease manifestation throughout the intestine.

*Chlamydia trachomatis* infection in humans is the most prevalent bacterial sexually transmitted infections worldwide.^28^ In addition to urogenital tissues, *Chlamydia* species can infect and cause pathology of other mucosal tissues such as ocular, respiratory, and intestinal tissues. Specific human-associated *C. trachomatis* serovars induce proctocolitis, known as lymphogranuloma venereum, that is often misdiagnosed as IBD.^29,30^ *Chlamydia* species, as obligate intracellular bacteria, primarily reside and replicate within epithelial cells, employing various immune evasion mechanisms.^31–33^ In turn, host cells use various defense strategies to mitigate infection. One such strategy is via the induction of indoleamine 2,3-dioxygenase 1 (IDO1), an enzyme that catalyzes the degradation of tryptophan, depriving *Chlamydia* of this essential amino acid as a tryptophan auxotroph.^34–37^ While potentially beneficial for the clearance of intracellular pathogens, IDO1 induction also promotes *Chlamydia* persistence^38^ and has broad impacts on the microenvironment, such as the specification of intestinal secretory cells through Notch signaling and the activation of regulatory T cells.^39–43^ In inflamed intestinal mucosa, including in human IBD, upregulation of IDO1 is prevalent; however, IDO1 has not been mechanistically connected to intracellular pathogens.^44–47^ How these host and bacterial components interact to drive or restrain chronic disease in a region-specific manner is incompletely understood.

In this study, we reveal the capacity for a specific pathobiont, *Chlamydia muridarum*, to drive an IDO1-mediated inflammatory response originating in secretory epithelial cells in the AC in the context of immune system dysregulation mimicking CD. We demonstrate similar pathways and pathobionts to be associated with the development of human CD in the AC but not TI or distal colon. Our study highlights how the interplay between genetic susceptibility and region-specific responses of epithelial cells contributes to the emergence of certain pathobionts and the development of disease.

## RESULTS

### The microbiome drives inflammation in the ascending colon in a genetically susceptible host with dysregulated TNF expression

While TNF is a critical cytokine for both ileal and colonic CD pathogenesis, how some patients develop inflammatory disease only in the TI, AC, or both is unclear.^48–52^ We hypothesize that exogenous factors, such as external stressors and specific microbes, influence the regional specificity of disease manifestation in a TNF-dependent manner. To investigate this, we used the Tnf^ΔARE/+^ mouse, an established model of Crohn’s disease driven by overexpression of TNF.^53^ These mice develop inflammation in the TI with only rare and mild inflammation reported in the colon.^53,54^ To determine if exogenous factors drive TNF-dependent inflammation in a region-specific manner, we reared mice in specific-pathogen-free barrier (SPF-B) and conventional (CONV) facilities, which differ in their animal management policies (Methods). Amongst age-matched mice, we found Tnf^ΔARE/+^ mice developed the expected terminal ileitis phenotype in SPF-B and CONV facilities (Figure 1A, Figure S1A). Surprisingly, Tnf^ΔARE/+^ mice reared in the CONV housing facility, but not those in SPF-B, developed severe colitis pronounced in the AC (Figure 1A-B, Figure S1B-C). Colonic inflammation was 100% penetrant in Tnf^ΔARE/+^ mice in the CONV facility and was established as early as 6 weeks of age (Figure S1D). Colonic inflammation initially developed in the AC of young mice with corresponding increases in TNF protein levels in this region, and it spread to the distal colon at later stages of disease in aged mice (Figure 1C-D, Figure S1C,E). The most prevalent features of colonic inflammation were an expansion of the lamina propria compartment and depth of inflammation extending into the submucosa and muscularis propria (Figure S1F). Of note, no cases of colonic inflammation were observed in Tnf^ΔARE/+^ mice from the SPF-B facility, nor any of the wildtype mice in either facility, even in mice aged up to one year (Figure S1G). Our findings indicate that exogenous factors can influence the site of inflammation within the gut of a genetically susceptible host, reminiscent of human CD where the TI and AC are the two most affected sites.

**Figure 1.**
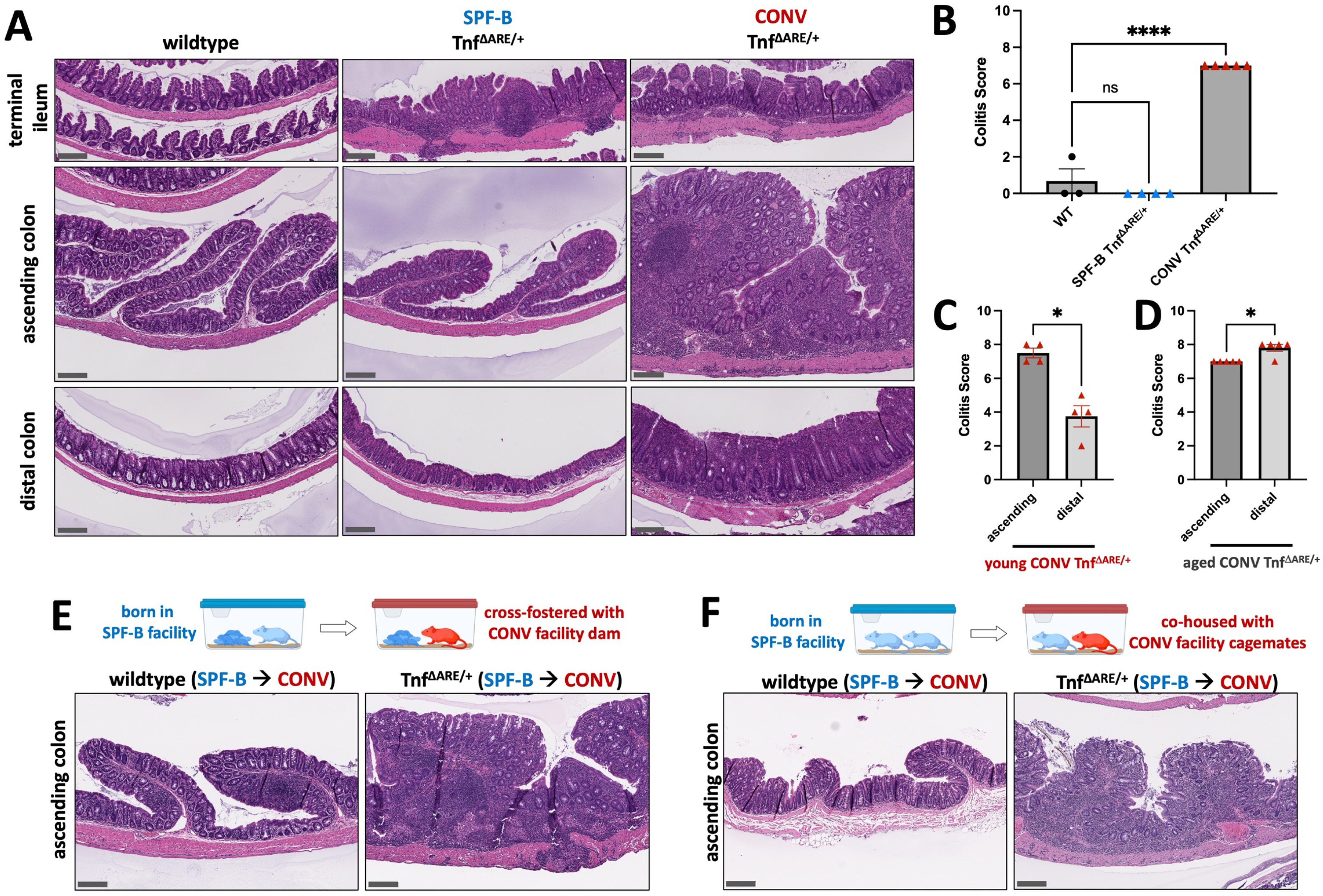
Crohn’s-like disease presents in the ascending colon as a function of murine housing facilities. **(A)** Representative H&E-stained intestinal sections (terminal ileum, ascending colon, distal colon) from wildtype (N = 3) and Tnf^ΔARE/+^ mice from SPF-B (N = 4) and CONV facilities (N = 5). Wildtype samples are from the CONV facility and all mice are age-matched (34-42 w of age). **(B)** Colitis scores from histopathological analysis of colons from CONV wildtype, SPF-B Tnf^ΔARE/+^, and CONV Tnf^ΔARE/+^ mice from the cohort in (A). Mean plus standard error of the mean (SEM) are shown and statistical significance was determined using an ordinary one-way ANOVA with CONV wildtype as the control group**. (C - D)** Colitis scores from histopathological scoring of colons from age-matched CONV Tnf^ΔARE/+^ mice (N = 4 for 12w of age in **(C)**, N = 5 for 34-42w of age in **(D)**), separated by ascending and distal colon regions. Mean plus SEM are shown and statistical significance was determined using paired t-tests. **(E)** Representative H&E-stained ascending colon sections from SPF-B wildtype Tnf^ΔARE/+^ mice transferred and co-housed/fostered as pups in the CONV facility with a wildtype or Tnf^ΔARE/+^ foster dam along with the dam’s age-matched biological pups. Pups were transferred and fostered within the first 3 days of birth and co-housed until experimental collection (37w of age, N = 3 wildtype, N = 4 Tnf^ΔARE/+^). **(F)** Representative H&E-stained ascending colon sections from SPF-B adult mice (N = 2 wildtype, N = 3 Tnf^ΔARE/+^) transferred and co-housed in the CONV facility in mixed-sex conditions until experimental collection (32-54w of age). CONV donors are of either wildtype or Tnf^ΔARE/+^ genotype. All scale bars = 200 µm. p-value * < 0.05, ** < 0.01, *** < 0.001. **** < 0.0001. **See also Figure S1.**

Human CD is considered a multifactorial disease with contributions from genetics, immune system dysregulation, environmental triggers, and the microbiome.^15,55^ We ruled out the influence of background genetics on colonic inflammation by meticulously backcrossing CONV facility mice to pure C57BL/6J breeders obtained from Jackson labs to avoid genetic drift (Methods). We performed single nucleotide polymorphism (SNP)-based background testing on mice from the CONV and SPF-B colonies to validate their inbred C57BL/6J or C57BL/6 backgrounds (Figure S1H), further demonstrating that background genetic differences do not drive colonic inflammation in the Tnf^ΔARE/+^ model. Next, we tested whether environmental triggers, such as caging conditions, food and water supply, or other unknown variables in the CONV facility were sufficient to induce colonic inflammation. We transferred colitis-free, SPF-B facility wildtype and Tnf^ΔARE/+^ mice to the CONV facility and subjected them to the same environment, but without co-housing with CONV facility mice. In this condition, neither wildtype nor Tnf^ΔARE/+^ mice developed colitis, suggesting that environmental triggers alone are not sufficient to drive colonic inflammation (Figure S1I). Altogether, our results showed that genetics and environmental stressors are not critical factors in driving colonic inflammation in Tnf^ΔARE/+^ mice.

We then examined the role of the microbiota by co-housing colitis-free, SPF-B wildtype and Tnf^ΔARE/+^ mice with wildtype or Tnf^ΔARE/+^ mice from the CONV facility. SPF-B Tnf^ΔARE/+^ mice that were transferred and co-housed developed ascending colitis, while transferred and co-housed SPF-B wildtype mice were free of colonic inflammation (Figure 1E-F, Figure S1J-K). Of note, Tnf^ΔARE/+^ mice developed ascending colonic inflammation under multiple experimental conditions, including transfer as pups to a foster dam (Figure 1E, Figure S1J) or transfer as adults (Figure 1F, Figure S1K), suggesting that the development of colonic inflammation is independent of the developmental stage of both the host’s immune system and microbiota. We demonstrated that the CONV microbiota, and not a specifically TNF-driven microbiota, is sufficient to confer inflammation of the AC, as wildtype foster dams were able to transfer colitis to SPF-B Tnf^ΔARE/+^ pups, as well as wildtype adults to SPF-B Tnf^ΔARE/+^ adults. Together, these results indicate that TNF^ΔARE/+^ mice, independent of background genetics, are susceptible to inflammation in the AC upon exposure to pathobiont microbes.

### Identification of *Chlamydia muridarum* as a pathobiont in TNF-associated inflammation in the AC

To identify pathobiont species that drive colonic inflammation, we performed shotgun metagenomics of the luminal contents of the AC to identify microbes associated with the most inflamed region (Table S1). Overall, the alpha diversity, at the species level, was not significantly different between genotypes and across facilities and ages, suggesting within-sample microbial diversity is similar across conditions (Figure S2A). In young mice examined at the phylum level, there were no major composition differences between wildtype and Tnf^ΔARE/+^ microbiota within each facility, suggesting TNF overexpression does not shift phylum level composition (Figure 2A-B). However, in the CONV facility, differences in phylum level composition emerged between wildtype and Tnf^ΔARE/+^ microbiota in aged conditions, suggesting TNF overexpression influences the composition of the AC microbiota at later stages of disease (Figure S2B-C). Moreover, differences between wildtype and Tnf^ΔARE/+^ beta diversity became more pronounced in aged conditions (Figure S2D).

**Figure 2.**
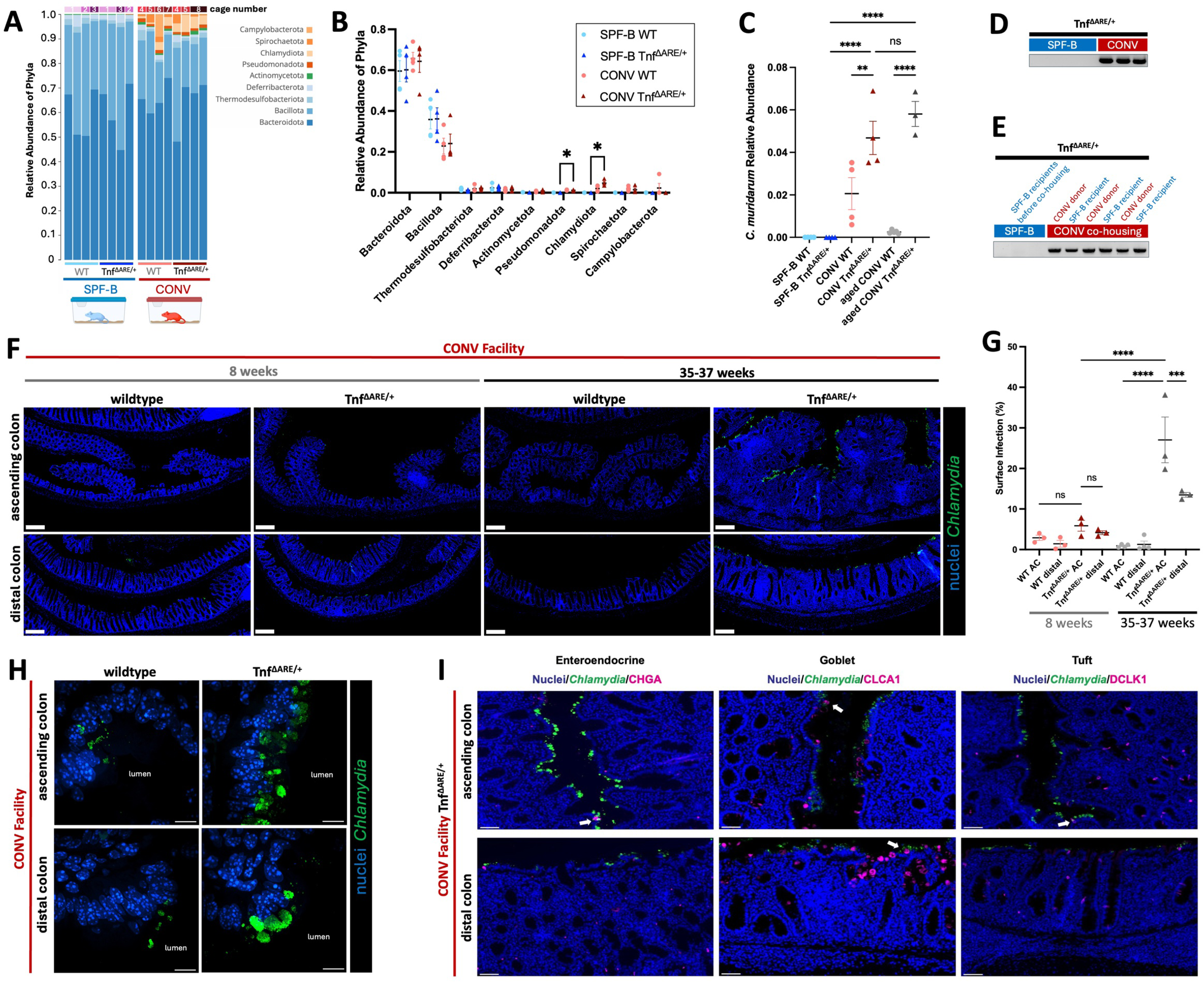
*Chlamydia muridarum* is associated with colonic inflammation in the context of TNF overexpression. **(A-B)** Shotgun metagenomic data, filtered for only the eubacteria kingdom, of proximal (ascending) colon luminal contents. Data is represented as relative abundance of mapped phyla for individual wildtype and Tnf^ΔARE/+^ mice in SPF-B and CONV facilities. N = 4 per condition from 8 cages. Mean plus standard error of the mean (SEM) are shown and statistical significance was determined using multiple unpaired t tests with false discovery rate (FDR) of 1%. **(C)** Shotgun metagenomic data with reads mapped to *C. muridarum* species. Mean plus standard error of the mean (SEM) are shown and statistical significance was determined using an ordinary one-way ANOVA with Sidak’s multiple comparison test. **(D)** Fecal DNA-based PCR testing for *C. muridarum* in Tnf^ΔARE/+^ mice from SPF-B (N = 4) and CONV (N = 3) facilities. Original gels shown in Data S1. **(E)** Fecal DNA-based PCR testing for *C. muridarum* in SPF-B Tnf^ΔARE/+^ mice before and after co-housing with Tnf^ΔARE/+^ mice from the CONV facility. N = 3 SPF-B Tnf^ΔARE/+^ recipients shown before and after co-housing with N = 3 donors. **(F)** Representative immunofluorescence (IF) images of *Chlamydia* major outer membrane protein (MOMP - green) and nuclei (Hoechst - blue) co-staining on colonic sections from wildtype (N = 3 at 8w of age, N = 4 at 35-37w of age) and Tnf^ΔARE/+^ (N = 3 at 8w of age, N = 3 at 35-37w of age) mice from the CONV facility. Scale bars = 200 µm. **(G)** Quantification of *Chlamydia* from IF images, separated by ascending colon (AC) and distal colon regions. Mean plus standard error of the mean (SEM) are shown and statistical significance was determined using an ordinary one-way ANOVA with Sidak’s multiple comparison test. **(H)** Representative confocal high-magnification IF stained images of *Chlamydia* major outer membrane protein (MOMP - green) and nuclei (Hoechst - blue) co-staining. Lumen side indicated in each image. N = 3 per condition. Scale bars = 10 µm. **(I)** Representative IF images of *Chlamydia* major outer membrane protein (MOMP - green) and nuclei (Hoechst - blue) co-staining with epithelial cell type-specific markers (CHGA – enteroendocrine cells, CLCA1 – goblet cells, DCLK1 – tuft cells) on colonic sections from age-matched CONV facility Tnf^ΔARE/+^ mice (N = 5, 35-37w of age). White arrows point to infected enteroendocrine, goblet, or tuft cells. Scale bars = 50 µm. p-value * < 0.05, ** < 0.01, *** < 0.001. **** < 0.0001. **See also Figure S2, Data S1, and Table S1.**

To identify specific pathobionts associated with colonic inflammation, we compared young Tnf^ΔARE/+^ mice across facilities and found Chlamydiota and Pseudomonadota phyla were enriched in the ACs of CONV mice compared to SPF-B mice (Figure 2A-B). Reads for the Chlamydiota phyla mapped to a single species *Chlamydia muridarum* - an obligate intracellular bacterium and the only natural *Chlamydia* pathogen of mice. *C. muridarum* is commonly used as a model to study human infections from *Chlamydia trachomatis,* a sexually transmitted disease-causing bacterium that is also associated with colonic inflammation.^29,30^ While *C. muridarum* was undetected in the AC microbiota of SPF-B mice of either genotype, the relative abundance of *C. muridarum* was higher in the AC microbiota of CONV Tnf^ΔARE/+^ mice compared to CONV wildtype, with more pronounced differences in aged conditions (Figure 2C). Moreover, the relative abundance of *C. muridarum* in the AC of Tnf^ΔARE/+^ mice was not significantly different in young versus aged mice. These results suggest that wildtype mice can mitigate *C. muridarum* colonization over time, but Tnf^ΔARE/+^ mice cannot control its growth. PCR-based fecal testing for *Chlamydia* confirmed all tested SPF-B Tnf^ΔARE/+^ mice were *Chlamydia*-negative while all CONV Tnf^ΔARE/+^ mice were *Chlamydia*-positive (Figure 2D, Data S1). We then analyzed fecal samples of mice in transfer and co-housing experiments. We found SPF-B Tnf^ΔARE/+^ mice were *Chlamydia-*positive upon co-housing, indicating that *Chlamydia* is a transmissible component of the microbiota that is associated with ascending colitis in Tnf^ΔARE/+^ mice (Figure 2E, Data S1). Given the role of *Chlamydia* species in human sexually transmitted disease, we asked whether sexual transmission is required to establish gastrointestinal colonization. We co-housed Tnf^ΔARE/+^ mice in same-sex conditions and found Tnf^ΔARE/+^ mice developed colonic inflammation, suggesting against a strictly sexually transmitted route and supporting an oral-fecal route (Figure S2E).^56,57^ Overall, these findings identify *C. muridarum* as a potential pathobiont associated with inflammation in the AC of Tnf^ΔARE/+^ mice.

Next, we used immunofluorescence (IF) microscopy and RNA fluorescence *in situ* hybridization (RNA-FISH) to assess the site of *C. muridarum* colonization along the length of the gastrointestinal tract of wildtype and Tnf^ΔARE*/+*^ mice. While *C. muridarum* was rarely detected in the small intestine, *C. muridarum* was detected at high levels in the AC of Tnf^ΔARE*/+*^ mice and progressively decreased in a gradient-like manner toward the distal colon consistent with the degree of inflammation (Figure 2F-G, Figure S2F-G). Consistent with shotgun metagenomics data, *C. muridarum* was not detected by IF microscopy in intestinal sections of SPF-B mice (Figure S2F, Figure S2H). In addition to IF-based detection of *Chlamydia*, we used RNA-FISH to detect *C. muridarum* 23S ribosomal RNA and found a similar pattern of infection in CONV wildtype and Tnf^ΔARE*/+*^ mice (Figure S2I). *Chlamydia*-infected cells were restricted to surface epithelial cells that line the colonic lumen and high-resolution imaging revealed that intracellular *C. muridarum* inclusions localized to the apical side of the cell, closely associated with the nucleus, and were larger in Tnf^ΔARE*/+*^ mice compared to wildtype mice (Figure 2H, Figure S2J). We then determined the cellular tropism of *C. muridarum* by examining its co-localization with colonic epithelial cell markers. We identified rare *Chlamydia*-positive enteroendocrine, goblet, and tuft cells at the crypt top (i.e. surface epithelium), while colocalization with colonocytes was abundant (Figure 2I). Given these different rates of colocalization amongst cell types, our results suggest that *Chlamydia* infection is primarily associated with absorptive colonocytes at the surface of the AC epithelium.

### *Chlamydia muridarum* is necessary and sufficient to drive TNF-dependent AC inflammation

Given the association of *C. muridarum* with inflammation of the AC, the role of specific *C. trachomatis* serovars in human colon pathology, and the role of TNF signaling in *Chlamydia*-induced pathology^58,59^, we assessed the necessity and sufficiency of *C. muridarum* in inducing colonic inflammation in Tnf^ΔARE*/+*^ mice. To determine necessity, we treated CONV wildtype and Tnf^ΔARE*/+*^ mice with doxycycline, a clinically effective antibiotic for clearing *Chlamydia* infections in humans (Figure 3A).^60^ *Chlamydia* was not detected by PCR in fecal samples from weanling-aged wildtype and Tnf^ΔARE/+^ mice treated with doxycycline for 1-2 weeks of treatment and remained undetectable at the time of harvest (Figure 3B-C, Figure S3A, Data S1). In mice and humans, the native gut microbiota gradually recovers after cessation of antibiotics.^61,62^ Thus, we evaluated the extent of inflammation in the AC 3-6 weeks after antibiotic withdrawal, a period during which the microbiome becomes re-established. Strikingly, doxycycline-treated Tnf^ΔARE/+^ mice had significantly lower inflammation scores in the AC compared to vehicle control treated aged-matched Tnf^ΔARE/+^ mice (Figure 3D-F, Figure S3B). Unlike inflammation in the AC, doxycycline did not reduce ileal inflammation in Tnf^ΔARE/+^ mice compared to vehicle control conditions, indicating that *C. muridarum* infection does not promote nor protect against ileal inflammation (Figure S3C). At the time of harvest, *C. muridarum* remained undetectable in shotgun metagenomic data of doxycycline-treated Tnf^ΔARE/+^ mice but was abundant in the vehicle control group (Figure 3G-H, Table S2). Given that doxycycline may deplete or permit a bloom of other microbes, we examined whether the recovered microbiome was broadly altered in the doxycycline-treated group and found that at the phylum level, only Chlamydiota and Thermodesulfobacteriodota were significantly reduced (Figure 3I). We ruled out the role of Thermodesulfobacteriodota in AC inflammation, as these microbes were also present in SPF-B mice without colonic inflammation (Figure 2A-B, Table S1). In aged mice with established inflammation prior to doxycycline administration, overall colitis scores and subscores trended lower upon treatment with doxycycline as compared to vehicle-treated controls, but statistically significant reduction was only in the lamina propria chronic inflammation subscore (Figure S3D-F). These results indicate that *C. muridarum* is required for inflammation in the AC without affecting ileitis, in a manner largely independent of other bacteria.

**Figure 3.**
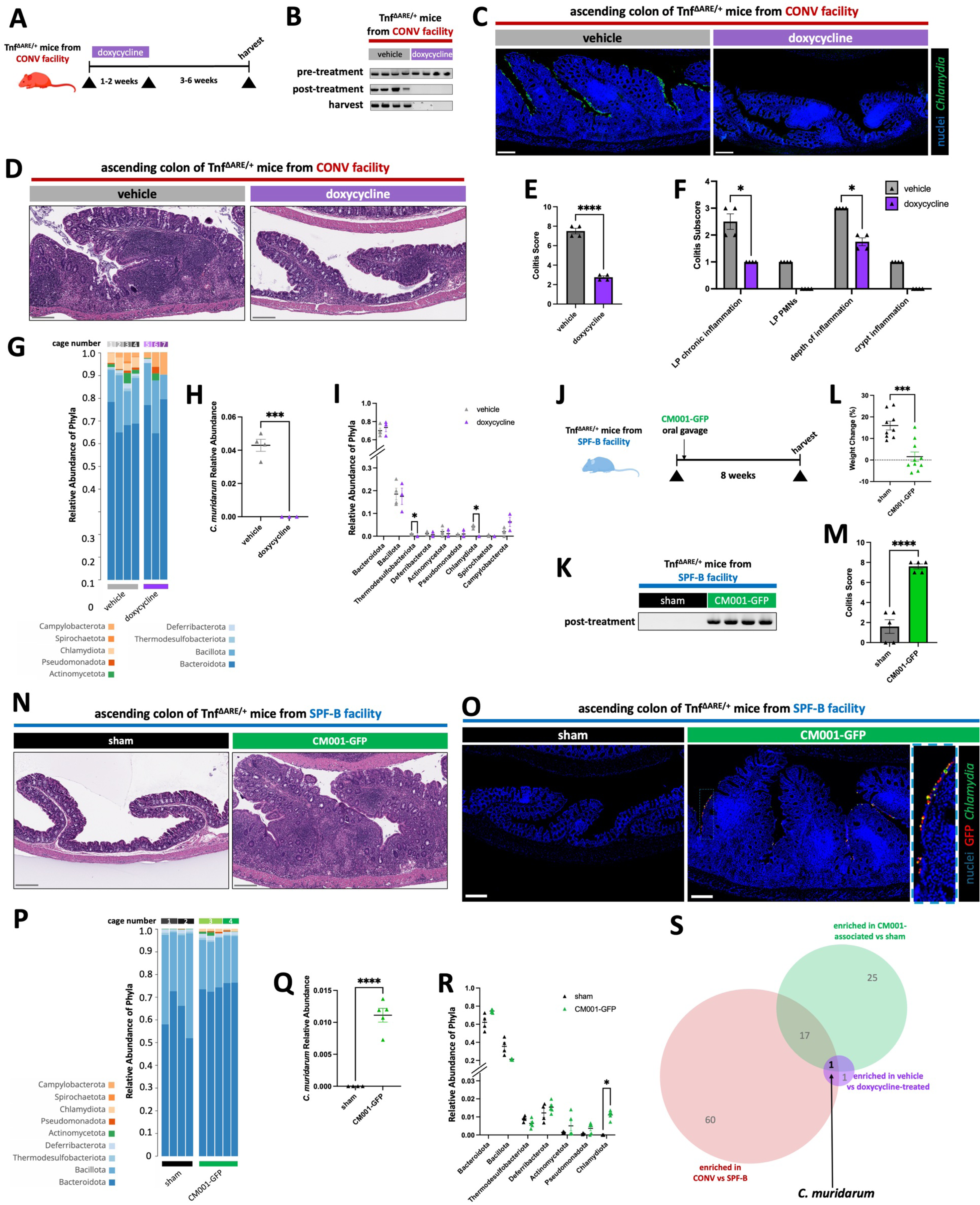
*Chlamydia muridarum* is necessary and sufficient to drive ascending colonic inflammation in the Tnf^ΔARE/+^ model of Crohn’s-like disease. **(A)** Experimental paradigm for doxycycline administration to Tnf^ΔARE/+^ mice in the CONV facility. **(B)** Fecal DNA-based PCR testing for *C. muridarum* in Tnf^ΔARE/+^ mice from doxycycline administration, where fecal DNA was tested pre-treatment, within 1 week of cessation of doxycycline, and prior to harvesting tissue. N = 4 mice per condition, age-matched at 11-12w of age at harvest. Original gels shown in Data S1. **(C)** Representative IF images of *Chlamydia* major outer membrane protein (MOMP - green) and nuclei (Hoechst - blue) co-staining on ascending colon sections from Tnf^ΔARE/+^ mice treated with doxycycline or vehicle. N = 4 mice per condition, age-matched at 11-12w of age at harvest. **(D)** Representative H&E-stained ascending colon sections from Tnf^ΔARE/+^ mice treated with doxycycline or vehicle. N = 4 mice per condition, age-matched at 11-12w of age at harvest. **(E)** Colitis scores from histopathological scoring of colons from Tnf^ΔARE/+^ mice in D. Mean plus SEM are shown, and statistical significance was determined using an unpaired t test. **(F)** Colitis subscores that contribute to overall colitis score in E. Mean plus SEM are shown, and statistical significance was determined using multiple unpaired t tests with FDR of 1%. LP = lamina propria. PMNs = polymorphonuclear leukocytes. **(G)** Shotgun metagenomic data, filtered for only the eubacteria kingdom, of proximal (ascending) colon luminal contents from Tnf^ΔARE/+^ mice treated with doxycycline or vehicle. Data is represented as relative abundance of mapped phyla for individual mice and grouped by doxycycline (N = 3) or vehicle treated (N = 4) conditions from 7 cages. Samples are from Tnf^ΔARE/+^ mice of mixed ages (11-23w of age). **(H)** Shotgun metagenomic data from G with reads specifically mapped to *C. muridarum* species. Mean plus standard error of the mean (SEM) are shown and statistical significance was determined using an unpaired t test. **(I)** Shotgun metagenomic data from G represented as relative abundance of mapped phyla for individual samples. Mean plus standard error of the mean (SEM) are shown, and statistical significance was determined using multiple unpaired t tests with false discovery rate (FDR) of 1%. **(J)** Experimental paradigm for CM001-GFP inoculation (3×10^6^ IFUs) of Tnf^ΔARE/+^ mice from the SPF-B facility. **(K)** Fecal DNA-based PCR testing for *C. muridarum* in Tnf^ΔARE/+^ mice from CM001-GFP inoculation, where fecal DNA was tested prior to harvesting tissue. N = 4 mice per condition, age-matched at 16-20w of age at harvest. **(L)** Body weights of mice from the CM001-GFP or sham-inoculated Tnf^ΔARE/+^ mice, where weight change was calculated at harvest as a percent of the individual mouse’s weight prior to treatment. N = 10 mice per condition, age-matched at 16-20w of age at harvest. **(M)** Colitis scores from histopathological scoring of colons from Tnf^ΔARE/+^ mice inoculated with CM001-GFP or sham. Mean plus SEM are shown, and statistical significance was determined using an unpaired t test. **(N)** Representative H&E-stained ascending colon sections from Tnf^ΔARE/+^ mice inoculated with CM001-GFP or sham. N = 10 mice per condition, age-matched at 16-20w of age at harvest. **(O)** Representative IF images of *Chlamydia* major outer membrane protein (MOMP - green), endogenous GFP from CM001-GFP (GFP – red), and nuclei (Hoechst - blue) co-staining on ascending colon sections from Tnf^ΔARE/+^ mice that are sham or CM001-GFP-inoculated. N = 5 mice per condition, age-matched at 16-20w of age at harvest. Inset to show colocalization of GFP and MOMP signal in CM001-GFP-inoculated conditions. **(P)** Shotgun metagenomic data, filtered for only the eubacteria kingdom, of proximal (ascending) colon luminal contents from Tnf^ΔARE/+^ mice in sham or CM001-GFP-inoculated conditions. Data is represented as relative abundance of mapped phyla for individual Tnf^ΔARE/+^ mice and grouped by CM001-GFP-inoculated (N = 5) or sham (N = 4) conditions from 4 cages. Samples are from Tnf^ΔARE/+^ mice aged-matched at 16-20w of age at collection. **(Q)** Shotgun metagenomic data from O with reads specifically mapped to *C. muridarum* species. Mean plus standard error of the mean (SEM) are shown and statistical significance was determined using an unpaired t test. **(R)** Shotgun metagenomic data from O represented as relative abundance of mapped phyla for individual samples. Mean plus standard error of the mean (SEM) are shown, and statistical significance was determined using multiple unpaired t tests with false discovery rate (FDR) of 1%. **(S)** Venn diagram depicting enriched species from shotgun metagenomic samples that are shared across all experimental paradigms. All scale bars = 200 µm. p-value * < 0.05, ** < 0.01, *** < 0.001. **** < 0.0001. **See also Figure S3, Figure S4, Table S2, Table S3, and Data S1.**

To determine whether *C. muridarum* is sufficient to induce inflammation in the AC, we inoculated *Chlamydia-*free Tnf^ΔARE/+^ mice from the SPF-B facility with a typed *C. muridarum* Nigg strain expressing GFP, CM001-GFP^63^, which shares ∼99% sequence similarity (ANIb = 0.988, TETRA = 0.999) to the *C. muridarum* strain we isolated from our CONV facility (*C. muridarum* strain VU, or “Cm-VU”) (Figure 3J, Figure S4A-B, Methods). CM001-GFP successfully engrafted into the microbiome of SPF-B Tnf^ΔARE/+^ mice after a single gavage of 3×10^6^ inclusion forming units (IFUs) without the need for antibiotic pretreatment (Figure 3K, Data S1). CM001-GFP-inoculated Tnf^ΔARE/+^ mice were not overtly ill compared to the sham-inoculated Tnf^ΔARE/+^ mice; however, they exhibited significantly lower weight gain compared to sham-inoculated Tnf^ΔARE/+^ controls, consistent with GI inflammation (Figure 3L). CM001-GFP-inoculated mice remain *Chlamydia*-positive at the time of harvest, and macroscopic and histological assessment revealed CM001-GFP-inoculated SPF-B Tnf^ΔARE/+^ mice developed severe colonic inflammation reminiscent of the disease in the CONV facility, while sham-inoculated Tnf^ΔARE/+^ mice showed no inflammation, similar to untreated SPF-B mice (Figure 3M-O, Figure S4C-E). To determine whether *C. muridarum* colonization indirectly drives additional community-level microbial changes to induce AC inflammation, we performed shotgun metagenomic sequencing of the AC luminal contents from CM001-GFP-inoculated and sham-inoculated mice (Figure 3P-R, Table S3). Compared to sham-inoculated mice, CM001-GFP-inoculated mice had no significant changes at the phylum level, except for Chlamydiota (Figure 3R), demonstrating CM001-GFP engraftment does not induce global shifts in the microbiome.

To confirm that *C. muridarum* is necessary and sufficient to induce colonic inflammation and demonstrate other species are not required, we examined shotgun metagenomic data across all experimental conditions at the species level (Figure S4F, Table S1-S3). Given the age differences across our datasets, we first determined whether age was associated with species enrichment. We found no significantly enriched species in CONV Tnf^ΔARE/+^ AC specimens between age groups (Figure S4G). To further discount age as a factor, we examined species enrichment in young SPF-B Tnf^ΔARE/+^ specimens versus aged CONV Tnf^ΔARE/+^ and young SPF-B Tnf^ΔARE/+^ specimens versus young CONV Tnf^ΔARE/+^ specimens and found 61 and 78 enriched species, respectively (Figure S4H-I). The enriched species were broadly similar in these two comparisons, further supporting that age does not significantly alter species enrichment (Figure S4J). Having discounted age as a significant contributor to species enrichment, we sought to identify pathobionts by comparing the enriched species in Tnf^ΔARE/+^ AC specimens from three experimental comparisons: SPF-B versus CONV, doxycycline-treated versus vehicle-treated, and sham-inoculated versus CM001-inoculated. Across these experiments, pathobionts necessary to drive colitis would be defined as those enriched in CONV Tnf^ΔARE/+^ compared to SPF-B Tnf^ΔARE/+^ specimens (Figure S4I), enriched in vehicle-treated control Tnf^ΔARE/+^ compared to doxycycline-treated Tnf^ΔARE/+^ specimens (Figure S4K), and enriched in CM001-GFP-inoculated Tnf^ΔARE/+^ compared to sham-inoculated Tnf^ΔARE/+^ specimens (Figure S4L). Comparative analysis showed that *C. muridarum* is the only species intersecting all experimental conditions (Figure 3S). Taken together, these results strongly support that *C. muridarum* is necessary and sufficient to drive AC inflammation in a genetically susceptible host, such as in the Tnf^ΔARE/+^ model.

### *Chlamydia* colonization induces indoleamine 2,3-dioxygenase (IDO1) expression in goblet cells

We sought to identify host responses to *C. muridarum* colonization with a focus on epithelial cells of the AC, as these cells are exclusive targets of *C. muridarum* infection. We performed single-cell RNA-sequencing (scRNA-seq) on the AC epithelia of wildtype and Tnf^ΔARE/+^ mice from SPF-B and CONV facilities, including young and aged Tnf^ΔARE/+^ mice from the CONV facility (Figure 4A, Figure S5A, Methods). Within SPF-B specimens, cells of wildtype and Tnf^ΔARE/+^ AC were intermixed in UMAP space, suggesting there are no major transcriptomic shifts in epithelial cell types induced by the Tnf^ΔARE/+^ genotype. Moreover, wildtype and Tnf^ΔARE/+^ cells from the CONV facility mostly intermixed in UMAP space, indicating that neither TNF overexpression nor *C. muridarum* abundance drastically alter colonic epithelial cells. While major transcriptomic differences amongst cell types were not observed, we aimed to determine whether epithelial cell type proportions were different amongst conditions (Figure 4B-C, Figure S5B, Table S4). Comparison of young CONV Tnf^ΔARE/+^ samples to CONV wildtype, aged CONV Tnf^ΔARE/+^, and young SPF-B Tnf^ΔARE/+^ conditions revealed only two statistically significant differences: a decrease in surface goblet cells and an increase in colonocyte progenitors in aged CONV Tnf^ΔARE/+^ AC as compared to young CONV Tnf^ΔARE/+^ AC (Figure 4D). Strikingly, there were no differences amongst epithelial cells between young CONV Tnf^ΔARE/+^ and young SPF-B Tnf^ΔARE/+^. These findings suggest that *C. muridarum* colonization does not induce changes in host responses by promoting the depletion or expansion of particular epithelial cell types of the AC, nor by drastically altering cell states, with the exception of a loss of surface goblet cells and expansion of colonocyte progenitors in later stages of inflammation.

**Figure 4.**
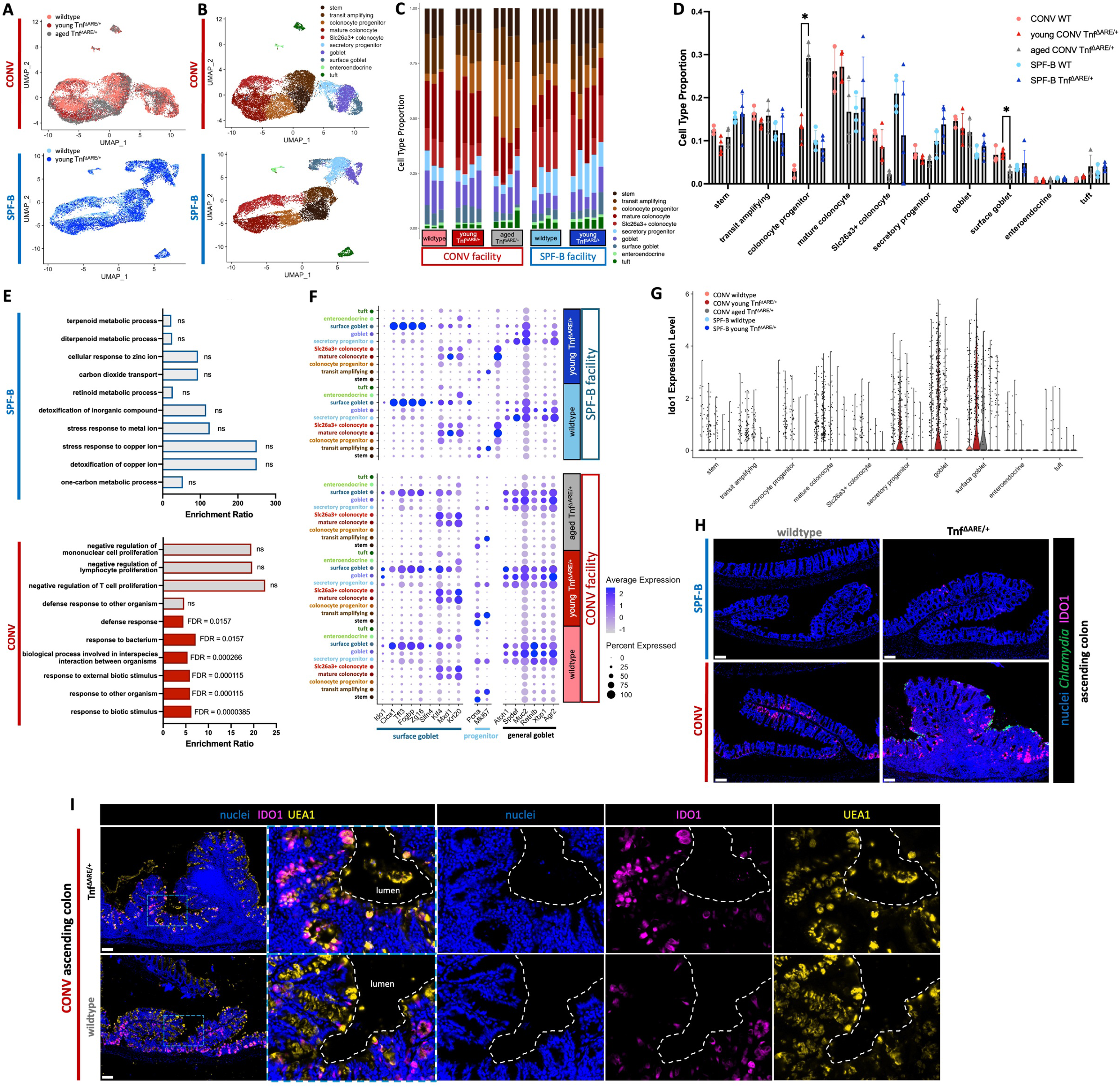
*Chlamydia muridarum* colonization upregulates ascending colonic goblet cell IDO1 expression. **(A)** UMAP co-embedding of scRNA-seq samples of AC epithelial cells from CONV wildtype (N = 3), CONV young Tnf^ΔARE/+^,(N = 4), CONV aged Tnf^ΔARE/+^ (N = 4), SPF-B wildtype (N = 4), and SPF-B young Tnf^ΔARE/+^ (N = 5) mice. Sample type overlay is indicated by color. **(B)** UMAP co-embedding of scRNA-seq data with cell type overlay indicated by color. **(C)** Bar graph of cell type proportions from scRNA-seq data, separated by sample type. **(D)** Statistical comparison of cell type proportions from scRNA-seq data separated by sample type. Mean plus standard error of the mean (SEM) are shown, and statistical significance was determined using multiple unpaired t tests with false discovery rate (FDR) of 1% between the following samples: CONV wildtype vs CONV young Tnf^ΔARE/+^, CONV young Tnf^ΔARE/+^ vs SPF-B young Tnf^ΔARE/+^, and CONV young Tnf^ΔARE/+^ vs CONV aged Tnf^ΔARE/+^. Only statistically significant results are shown. **(E)** Gene Ontology of Biological Process terms derived from over-representation analysis (ORA) of upregulated genes in cells from the SPF-B facility (top) or CONV facility (bottom). Input to ORA included all upregulated genes identified by differential expression performed for each cell type between CONV and SPF-B facilities, pooling all genotypes (wildtype and Tnf^ΔARE/+^) and age (young and aged) conditions in each facility. Enrichment ratio is shown and statistical significance was determined using false discovery rate (FDR) < 0.05. **(F)** Dot plot of gene expression from scRNA-seq data separated by sample type. Selected marker genes for surface goblet cells, progenitors, and goblet cells. Color indicates expression level while circle size represents percent of cells expressing the gene. **(G)** Violin plot of *Ido1* expression in each cell type from scRNA-seq data separated by sample type. **(H)** Representative IF images of IDO1 (magenta), *Chlamydia* major outer membrane protein (green), and nuclei (Hoechst - blue) co-staining on ascending colon sections from wildtype and Tnf^ΔARE/+^ mice from the SPF-B and CONV facilities. N = 3 mice, age-matched at 34-42w of age at harvest. Scale bars = 100 µm. **(I)** Representative IF images of IDO1 (magenta), UEA1 lectin (yellow), and nuclei (Hoechst - blue) co-staining on ascending colon sections from wildtype and Tnf^ΔARE/+^ mice from the CONV facility. Inset image to show colocalization of UEA1 lectin, a goblet and secretory granule marker, with IDO1. N = 3 mice, age-matched at 16-17w of age at harvest. Scale bars = 100 µm. p-value * < 0.05, ** < 0.01, *** < 0.001. **** < 0.0001. **See also Figure S5, Figure S6, Table S4, Table S5, Table S6, and Table S7.**

Next, we aimed to identify cell type specific changes in gene expression in *Chlamydia-*colonized (CONV facility) vs uncolonized (SPF-B facility) conditions. Surprisingly, our analysis revealed only a limited number of differentially expressed genes when comparing individual AC epithelial cell types between CONV and SPF-B housed mice (Figure S5C, Table S5). We performed a more direct comparison between young Tnf^ΔARE/+^ samples in the CONV and young Tnf^ΔARE/+^ samples in SPF-B conditions, which yielded a similar set of differentially expressed genes to those in the broader comparison, most of which were shared between comparisons (Figure S5C, Table S6). These results support that *C. muridarum* colonization induces modest transcriptomic changes in wildtype and Tnf^ΔARE/+^ AC epithelial cell types.

To reveal specific gene program alterations, we performed gene over-representation analysis (ORA) using upregulated genes from all cell types in the CONV or SPF-B facilities, separately, as input (Figure 4E, Table S7). Terms related to colonic cell function were enriched in SPF-B specimens while terms related to host responses to microbes were enriched in CONV specimens. These results indicate a loss of normal function and induction of defense responses in the AC epithelial cells from *Chlamydia-*positive mice in the CONV facility. We examined genes contributing to the ORA terms and found that most were expressed in goblet cell subtypes and secretory progenitors, with the most prominent and statistically significant differences in surface goblet cells. Among these was *Ido1,* a gene encoding indoleamine 2,3-dioxygenase (IDO1), an enzyme that converts tryptophan to kynurenine.^35,36^ This enzyme has established roles in innate immune protection from infection, including from *Chlamydia* and other pathogens that require host-derived tryptophan.^34,37^ Moreover, IDO1 has immunosuppressive functions through kynurenine-mediated modulation of regulatory T cells and, additionally, has a non-enzymatic role that promotes goblet cell differentiation.^39–43^ Intriguingly, *Ido1* was expressed at high levels in a subset of goblet cells and surface goblet cells in the AC of mice from the CONV facility and at low levels in a small subset of goblet cells from the AC of mice from the SPF-B facility (Figure 4F-G, Figure S6A). Within the CONV facility, *Ido1* was expressed at higher levels in goblet and surface goblet cells of Tnf^ΔARE/+^ mice compared to wildtype mice. We used IF microscopy to confirm the expression of IDO1 protein. Consistent with transcriptomic findings, IDO1 was expressed at significantly higher levels in samples from the CONV facility compared to the SPF-B facility in both wildtype and Tnf^ΔARE/+^ AC, with elevated signal in Tnf^ΔARE/+^ AC, and was expressed almost exclusively by goblet cells (Figure 4H-I, Figure S6B-C). These results suggest a potential role for goblet cells in *Chlamydia*-induced AC inflammation via upregulation of IDO1 in goblet cells.

### Chronic inflammation in the ascending colon is potentiated through a goblet cell indoleamine 2,3-dioxygenase (IDO1)-dependent mechanism

We next investigated whether IDO1 expression was dependent on the presence of *Chlamydia* through perturbation experiments. In CONV Tnf^ΔARE/+^ mice treated with doxycycline to clear *Chlamydia,* we found IDO1 expression was substantially lower in the AC as compared to vehicle-treated Tnf^ΔARE/+^ mice (Fig 5A, Figure S7A). Consistently, SPF-B Tnf^ΔARE/+^ mice inoculated with *C. muridarum* (CM001-GFP) showed increased IDO1 expression compared to sham-inoculated Tnf^ΔARE/+^ mice, on a similar level to CONV facility Tnf^ΔARE/+^ mice colonized by Cm-VU (Fig 5B, Figure S7B). These results demonstrate that upregulation of IDO1 is induced by *C. muridarum* and is potentially related to inflammation of the AC.

**Figure 5.**
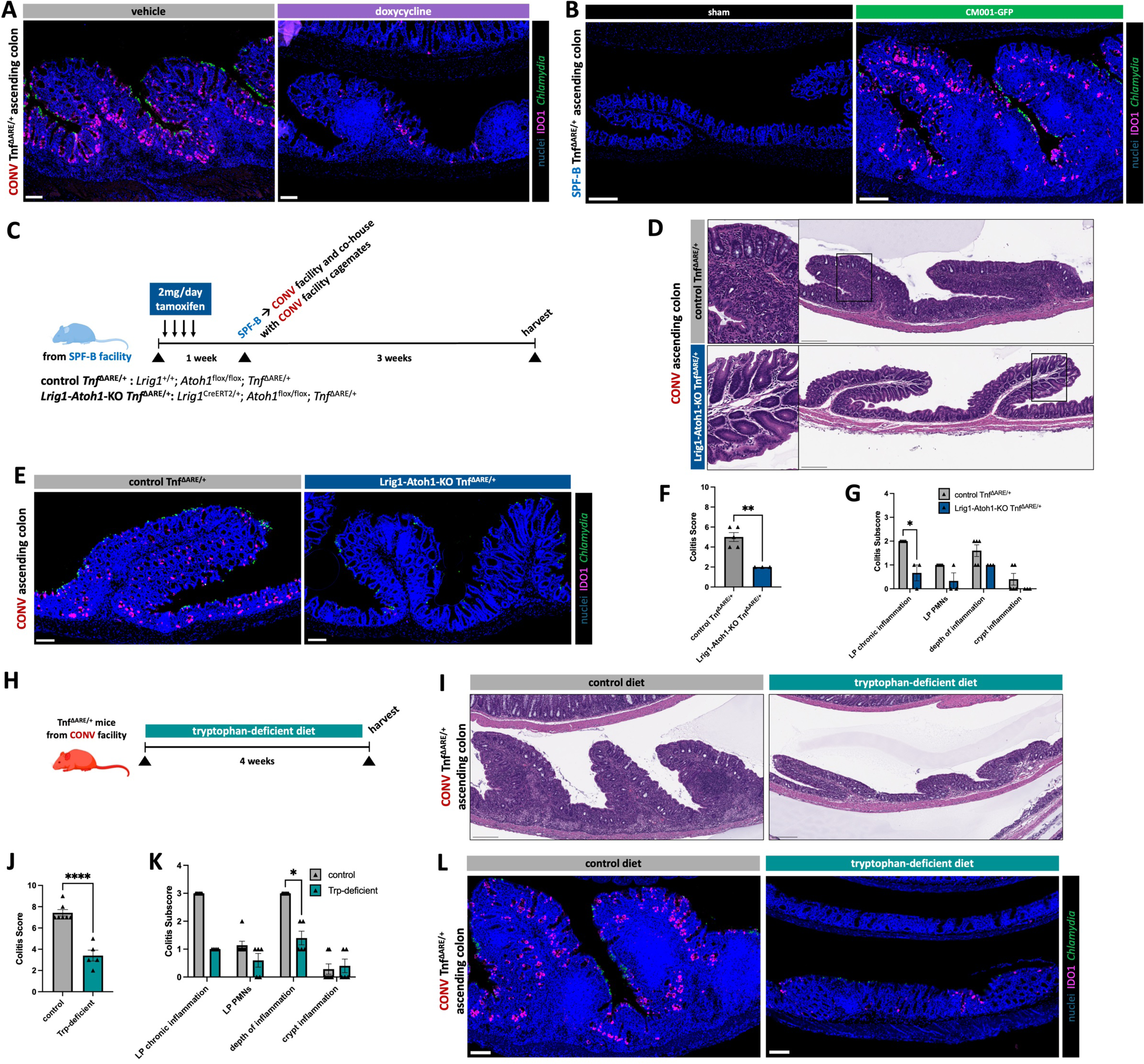
Perturbation of the IDO1 pathway reduces Chlamydia-driven ascending colon inflammation in the Tnf^ΔARE/+^ model. **(A)** Representative IF images of IDO1 (magenta), *Chlamydia* major outer membrane protein (MOMP - green), and nuclei (Hoechst - blue) co-staining on ascending colon sections from CONV Tnf^ΔARE/+^ mice treated with doxycycline or vehicle. N = 4 mice per condition, age-matched at 11-12w of age at harvest. Scale bars = 100 µm. **(B)** Representative IF images of IDO1 (magenta), *Chlamydia* major outer membrane protein (MOMP - green), and nuclei (Hoechst - blue) co-staining on ascending colon sections from SPF-B Tnf^ΔARE/+^ mice that are sham or CM001-GFP-inoculated. N = 5 mice per condition, age-matched at 16-20w of age at harvest. Scale bars = 200 µm. **(C)** Experimental paradigm for secretory cell ablation in Tnf^ΔARE/+^ mice from the SPF-B facility, subsequently transferred to the CONV facility to be co-housed for 3 weeks with *Chlamydia-*positive cagemates. **(D)** Representative H&E-stained ascending colon sections from Tnf^ΔARE/+^ mice with or without secretory cell ablation. N = 5 control Tnf^ΔARE/+^ mice, N = 3 Lrig1-Atoh1-KO Tnf^ΔARE/+^ mice, age-matched at 20-27w of age at harvest. Scale bars = 200 µm. Insets to show lack of goblet cell granules in the Lrig1-Atoh1-KO condition. **(E)** Representative IF images of IDO1 (magenta), *Chlamydia* major outer membrane protein (MOMP - green), and nuclei (Hoechst - blue) co-staining on ascending colon sections from Tnf^ΔARE/+^ mice with or without secretory cell ablation. N = 5 control Tnf^ΔARE/+^ mice, N = 3 Lrig1-Atoh1-KO Tnf^ΔARE/+^ mice, age-matched at 20-27w of age at harvest. Scale bars = 100 µm. **(F)** Colitis scores from histopathological scoring of colons from Tnf^ΔARE/+^ mice from the secretory cell ablation experiment. Mean plus SEM are shown, and statistical significance was determined using an unpaired t test. **(G)** Colitis subscores that contribute to overall colitis score. Mean plus SEM are shown, and statistical significance was determined using multiple unpaired t tests with FDR of 1%. **(H)** Experimental paradigm for administration of tryptophan-deficient diet to Tnf^ΔARE/+^ mice from the CONV facility. **(I)** Representative H&E-stained ascending colon sections from CONV Tnf^ΔARE/+^ mice fed a control diet or a tryptophan-deficient diet. N = 7 Tnf^ΔARE/+^ mice on control diet, N = 5 Tnf^ΔARE/+^ mice on tryptophan-deficient diet, age-matched at 11-12w of age at harvest. Scale bars = 200 µm. **(J)** Colitis scores from histopathological scoring of colons from CONV Tnf^ΔARE/+^ fed a control diet or a tryptophan-deficient diet. Mean plus SEM are shown, and statistical significance was determined using an unpaired t test. **(K)** Colitis subscores that contribute to overall colitis score. Mean plus SEM are shown, and statistical significance was determined using multiple unpaired t tests with FDR of 1%. **(L)** Representative IF images of IDO1 (magenta), *Chlamydia* major outer membrane protein (MOMP - green), and nuclei (Hoechst - blue) co-staining on ascending colon sections from CONV Tnf^ΔARE/+^ mice fed a control diet or a tryptophan-deficient diet. N = 7 Tnf^ΔARE/+^ mice on control diet, N = 5 Tnf^ΔARE/+^ mice on tryptophan-deficient diet, age-matched at 11-12w of age at harvest. Scale bars = 100 µm. p-value * < 0.05, ** < 0.01, *** < 0.001. **** < 0.0001. **See also Figure S7 and Figure S8.**

We next investigated the relationship between IDO1 expression in goblet cells and *Chlamydia*-induced AC inflammation. Since IDO1 expression is restricted to goblet cells, we induced secretory cell ablation in *C. muridarum-*free SPF-B Tnf^ΔARE/+^ mice by knocking out the master secretory cell transcription factor *Atoh1* in intestinal and colonic stem cells (*Lrig1^CreERT2/+^;Atoh1^fl/fl^* - “Lrig1-Atoh1-KO”).^64,65^ After Cre-mediated knockout of *Atoh1*, goblet cells within the AC of Lrig1-Atoh1-KO Tnf^ΔARE/+^ mice were lost (Figure 5C-D, Figure S7C). To colonize with *C. muridarum*, SPF-B Lrig1-Atoh1-KO Tnf^ΔARE/+^ and Tnf^ΔARE/+^ control mice were co-housed with CONV facility cagemates naturally colonized with Cm-VU (Figure 5C). While *C. muridarum* was successfully transferred in all conditions, IDO1 expression was no longer detected in the AC epithelium of Lrig1-Atoh1-KO Tnf^ΔARE/+^ mice (Figure 5E, Figure S7D). Remarkably, AC inflammation in Lrig1-Atoh1-KO Tnf^ΔARE/+^ mice was suppressed, while secretory cell-replete Tnf^ΔARE/+^ mice presented with severe AC inflammation (Figure 5D, Figure 5F-G, Figure S7C). Given the localization of *C. muridarum* to the colonic surface and high *Ido1* expression in surface goblet cells, we repeated the same experiment where only the surface fraction of *Krt20+* goblet cells were depleted by Cre-mediated knockout of *Atoh1* (*Krt20^CreERT2/+^;Atoh1^fl/fl^* - “Krt20-Atoh1-KO”, Methods). Indeed, only goblet cells at the crypt surface in the AC and villi of the TI were depleted after tamoxifen induction in this model (Figure S8A). Like the Tnf^ΔARE/+^; Lrig1-Atoh1-KO model where all goblet cells were depleted, surface goblet cell depleted Krt20-Atoh1-KO Tnf^ΔARE/+^ mice exhibited lower AC inflammation (Figure S8B). These results demonstrate the role of host responses in goblet cells, which is the main source of IDO1 expression, in inducing inflammation in the AC following *C. muridarum* colonization.

To test if IDO1 enzymatic activity of converting tryptophan to kynurenine was responsible for driving AC inflammation, we removed tryptophan from the diet of Tnf^ΔARE/+^ mice in the CONV facility (Figure 5H). AC inflammation of Tnf^ΔARE/+^ mice on a tryptophan-deficient diet was suppressed compared to Tnf^ΔARE/+^ mice on a control diet (Fig 5I-K, Figure S8C). The ACs of Tnf^ΔARE/+^ mice on tryptophan-deficient diet remained colonized by *C. muridarum*. However, *C. muridarum* colonization and IDO1 induction were diminished with the tryptophan-deficient diet (Figure 5L, Figure S8D). Taken together, these results demonstrate that *C. muridarum*-induced upregulation of IDO1 in goblet cells promotes chronic AC inflammation in a genetically susceptible host.

### *Chlamydia-*induced colonic inflammation is independent of upstream ileal inflammation or antimicrobial function

Given that AC and TI inflammation in the Tnf^ΔARE/+^ model occur concomitantly, we asked whether upstream inflammation in the small intestine is required for *C. muridarum-*induced inflammation in the AC. We leveraged a murine model where a genetic insert disrupts the *Tnf* gene and leads to TNF protein expression at an intermediate level between that of wildtype and Tnf^ΔARE/+^ mice (“Tnf^Δreg/+^”, or Tnf^ΔAREneo/+^ in Roulis *et al.,* 2011 Figure S9A, Methods).^66^ Like Tnf^ΔARE/+^ mice, Tnf^Δreg/+^ mice develop inflammation in the AC when reared in the CONV facility (Figure 6A-B, Supp 9B). Importantly, Tnf^Δreg/+^ mice did not develop concomitant ileitis, demonstrating independence between TI and AC inflammation (Figure 6C-D, Figure S9C). Similar to CONV Tnf^ΔARE/+^ mice treated with doxycycline, AC inflammation was significantly reduced when CONV facility Tnf^Δreg/+^ mice were treated with doxycycline to clear *C. muridarum,* and the TI remained uninflamed (Figure 6A-E, Figure S9B-D). Moreover, Tnf^Δreg/+^ mice in *Chlamydia-*free conditions, achieved by rederivation from sperm into *Chlamydia*-free dams (rederived Tnf^Δreg/+^), were expectedly free of *Chlamydia* and their TI and AC were not inflamed, further supporting the role of *C. muridarum* in driving AC inflammation (Fig 6A-E, Figure S9B-D, Methods). The epithelial mechanisms driving inflammation are likely conserved as evidenced by high expression of IDO1 in AC goblet cells from CONV Tnf^Δreg/+^ mice, compared to low IDO1 expression in the AC of doxycycline-treated Tnf^Δreg/+^ mice and rederived Tnf^Δreg/+^ mice into *Chlamydia-*free conditions (Figure 6E-F, Figure S9D-E). These findings demonstrate that inflammation in the AC occurs independently of upstream TI inflammation and is driven by IDO1 expression in goblet cells in response to *C. muridarum* colonization in the context of TNF overexpression.

**Figure 6.**
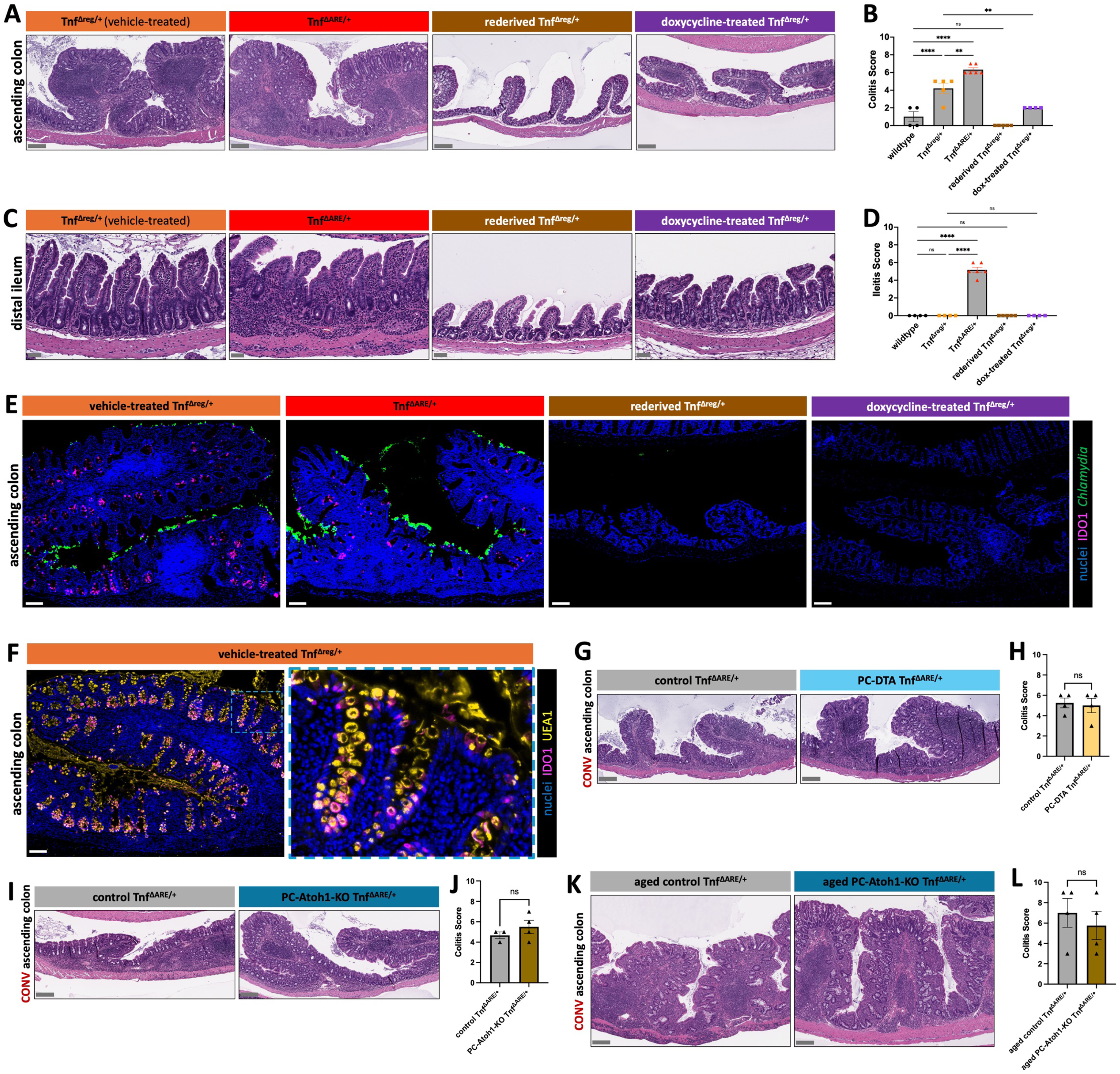
Inflammation and secretory function of the small intestine are independent of ascending colon inflammation. **(A)** Representative H&E-stained ascending colon sections from CONV Tnf^Δreg/+^ (N = 5), CONV Tnf^ΔARE/+^ (N = 6), rederived Chlamydia-negative CONV Tnf^Δreg/+^ (N = 5), and doxycycline-treated CONV Tnf^ΔARE/+^ (N = 4) mice. Mice are age-matched at 16-17w of age at harvest. Scale bars = 200 µm. **(B)** Colitis scores of ascending colons of TNF mutant mice and wildtype mice (N = 4). Mean plus SEM are shown, and statistical significance was determined using an ordinary one-way ANOVA with multiple comparisons. **(C)** Representative H&E-stained terminal ileum sections from CONV Tnf^Δreg/+^ (N = 5), CONV Tnf^ΔARE/+^ (N = 6), rederived Chlamydia-negative CONV Tnf^Δreg/+^ (N = 5), and doxycycline-treated CONV Tnf^ΔARE/+^ (N = 4) mice. Mice are age-matched at 16-17w of age at harvest. Scale bars = 50 µm. **(D)** Ileitis scores of terminal ilea of TNF mutant mice and wildtype mice (N = 4). Mean plus SEM are shown, and statistical significance was determined using an ordinary one-way ANOVA with multiple comparisons. **(E)** Representative IF images of IDO1 (magenta), *Chlamydia* major outer membrane protein (MOMP - green), and nuclei (Hoechst - blue) co-staining of ascending colon sections from CONV Tnf^Δreg/+^, CONV Tnf^ΔARE/+^, rederived Chlamydia-negative CONV Tnf^Δreg/+^, and doxycycline-treated CONV Tnf^ΔARE/+^ mice. N = 3 mice per condition, age-matched at 16-17w of age at harvest. Scale bars = 100 µm. **(F)** Representative IF images of IDO1 (magenta), UEA1 lectin (yellow), and nuclei (Hoechst - blue) co-staining of ascending colon sections from CONV Tnf^Δreg/+^ mice. Inset image to show colocalization of UEA1 lectin, a goblet and secretory granule marker, with IDO1. N = 3 mice, age-matched at 16-17w of age at harvest. Scale bars = 100 µm. **(G)** Representative H&E-stained ascending colon sections from control Tnf^ΔARE/+^ (N = 4) and PC-DTA Tnf^ΔARE/+^ (N = 4) mice. Mice are from the CONV facility and are age-matched at 8w of age at harvest. Scale bars = 200 µm. **(H)** Colitis scores of ascending colons from control Tnf^ΔARE/+^ and PC-DTA Tnf^ΔARE/+^ mice. Mean plus SEM are shown, and statistical significance was determined using an unpaired t test. **(I)** Representative H&E-stained ascending colon sections from control Tnf^ΔARE/+^ (N = 3) and PC-Atoh1-KO Tnf^ΔARE/+^ (N = 4) mice. Mice are from the CONV facility and are age-matched at 6-10w of age at harvest. Scale bars = 200 µm. **(J)** Colitis scores of ascending colons from control Tnf^ΔARE/+^ and PC-Atoh1-KO Tnf^ΔARE/+^ mice. Mean plus SEM are shown, and statistical significance was determined using an unpaired t test. **(K)** Representative H&E-stained ascending colon sections from aged control Tnf^ΔARE/+^ (N = 4) and aged PC-Atoh1-KO Tnf^ΔARE/+^ (N = 4) mice. Mice are from the CONV facility and are age-matched at 23-64w of age at harvest. Scale bars = 200 µm. **(L)** Colitis scores of ascending colons from aged control Tnf^ΔARE/+^ and aged PC-Atoh1-KO Tnf^ΔARE/+^ mice. Mean plus SEM are shown, and statistical significance was determined using an unpaired t test. p-value * < 0.05, ** < 0.01, *** < 0.001. **** < 0.0001. **See also Figure S9, Figure S10, and Figure S11.**

While our results show that upstream TI inflammation is not required for the development of *C. muridarum*-induced colitis, we asked whether antimicrobials produced by small intestinal Paneth cells, which are deposited into mucus that flows into the ascending colon^67–69^, would offer protection by modulating *C. muridarum* colonization. Loss of Paneth cell function is observed in inflamed CD lesions and has been implicated in pathogenesis, and progressive loss of these cells in inflamed regions is characteristic of the Tnf^ΔARE/+^ model of intestinal inflammation.^70–73^ To determine whether loss of Paneth cell function promotes a bloom of *C. muridarum* to drive colonic inflammation, we developed two models (*Defa4^Cre/+^;Atoh1^f/fl^* - “PC-Atoh1-KO” and *Defa4^Cre/+^;Rosa^LSL-DTA/+^* - “PC-DTA”) that achieve long-term ablation of Paneth cells in the Tnf^ΔARE/+^ model (Figure S10A-D, Methods). In these two models, there was no significant difference in the colonic inflammation score compared to age-matched Tnf^ΔARE/+^ controls, indicating Paneth cells do not promote nor protect the AC from inflammation (Figure 6G-J, Figure S11A-B). Moreover, Paneth cell-ablated Tnf^ΔARE/+^ mice displayed a similar number of *C. muridarum* inclusions when compared to control Tnf^ΔARE/+^ mice and IDO1 upregulation was sustained, suggesting Paneth cells do not modulate *C. muridarum* colonization (Figure S11C-D). PC-Atoh1-KO mice exhibit a loss of all secretory cells in the small intestine as a function of age, but have no alterations in secretory cell specification in the colon (Figure S11E-F). This model enables us to specifically examine the role of small intestinal secretory cells in AC inflammation. Aged PC-Atoh1-KO Tnf^ΔARE/+^ mice exhibited colonic inflammation scores similar to those of age-matched control Tnf^ΔARE/+^ mice, suggesting that colonic secretory cells, and not small intestinal secretory cells, promote *C. muridarum-*induced AC inflammation (Fig 6K-L, Figure S11G). Taken together, our results suggest that AC inflammation develops independently of inflammation or secretory cell function in the small intestine, and instead, is driven by specific microbial triggers and IDO1 induction in colonic goblet cells that together drive aberrant inflammatory responses in the colon.

### IDO1 activation associated with intracellular microbes is a hallmark of Crohn’s Disease patients with ascending colon involvement

We showed that an intracellular microbe, *C. muridarum*, can induce AC inflammation in a genetically susceptible host (TNF overexpression). We next investigated the generalizability of this mechanism. IL-10 is an anti-inflammatory cytokine that is implicated in the pathogenesis of CD, and IL-10 null mice are widely recognized as a colitis model which requires a microbial trigger to induce pathology.^26,74–82^ Consistent with our Tnf^ΔARE/+^ model, IL-10rb^-/-^ mice co-housed with *C. muridarum*-positive CONV facility cagemates developed AC inflammation (Figure S12A). In contrast, IL-10rb^-/-^ mice in isolated caging displayed no AC inflammation. Co-housed IL-10rb^-/-^ mice were *C. muridarum-*positive and had elevated IDO1 expression in epithelial cells as compared to isolated caging controls that were *Chlamydia-*negative with low expression of IDO1 (Figure S12B). These results demonstrate the generalizability of the intracellular microbe, *Chlamydia,* in triggering inflammation specifically in the AC of a host with genetic susceptibility to immune disruptions, mirroring mechanisms implicated in human IBD.

We next investigated whether IDO1 upregulation and epithelia-associated microbes are associated with human CD with active AC involvement. We analyzed a human scRNA-seq dataset of CD AC and TI specimens, where a novel population of epithelial cells, termed LND cells, were identified and associated with CD activity (Figure 7A-B, Figure S12C).^83^ CD AC specimens with active inflammation had significantly higher proportions of LND cells compared to inactive CD of the AC or normal AC (Figure 7C). We examined *IDO1* expression in AC specimens and found expression was restricted to CD specimens with a history (active or inactive) of AC inflammation (Figure 7D). Intriguingly*, IDO1* was almost exclusively expressed by the LND cell subpopulation and was significantly higher in patients with active AC inflammation compared to those with inactive AC inflammation (Figure 7E-F, Figure S12D). Using IF microscopy, we confirmed IDO1 protein expression was upregulated in epithelial cells from CD AC samples with active AC inflammation, and was not expressed by epithelial cells from CD AC samples without AC inflammation (Figure 7G, Figure S12E). Notably, *IDO1* expression was largely undetected in TI epithelial cells, further supporting the notion that epithelial IDO1 expression contributes to region-specific susceptibility of inflammation in the AC (Figure S12F-H).

**Figure 7.**
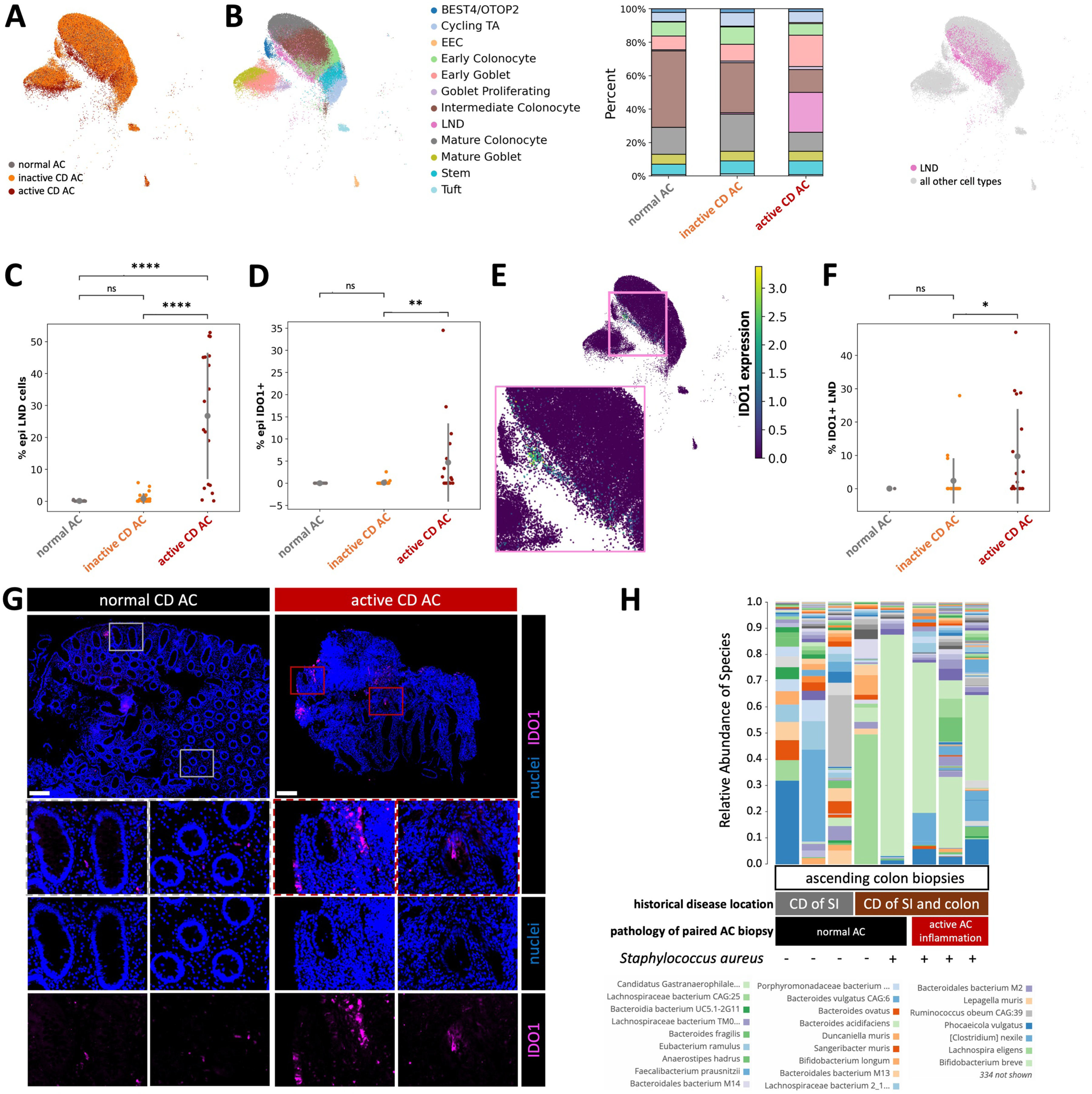
Analysis of human Crohn’s disease in the AC reveals epithelial IDO1 upregulation coupled to alternative microbial colonization. **(A)** UMAP co-embedding of human scRNA-seq data of AC epithelial cells from normal (N = 15), inactive CD (N = 34), and active CD (N = 19) specimens. Sample type overlay is indicated by color. Normal AC is composed of healthy control specimens, inactive CD AC is composed of CD specimens histopathologically scored as normal or quiescent, and active CD AC is composed of CD specimens histopathologically scored as mild, moderate, or severe. **(B)** Cell type breakdown of AC scRNA-seq data: UMAP with cell type overlay indicated by color (left), barplot with cell type proportion indicated by color, separated by sample type (middle), UMAP with LND cell overlay indicated by color (right). **(C)** Statistical comparison of LND cell type proportion amongst all epithelial cells from AC scRNA-seq data separated by sample type. Mean plus standard deviation are shown, and statistical significance was determined using independent t tests. **(D)** Statistical comparison of IDO1-expressing cell proportions amongst all epithelial cells from AC scRNA-seq data separated by sample type. Mean plus standard deviation are shown, and statistical significance was determined using independent t tests. **(E)** UMAP of AC scRNA-seq data with overlay of *IDO1* gene expression indicated by the color gradient. Inset to show *IDO1* expression in the LND cluster. **(F)** Statistical comparison of the proportion of IDO1-expressing LND cells amongst all epithelial cells from AC scRNA-seq data separated by sample type. Mean plus standard deviation are shown, and statistical significance was determined using independent t tests. **(G)** Representative IF images of IDO1 (magenta) and nuclei (Hoechst - blue) co-staining on AC biopsy sections from CD specimens with normal (N = 3) or active (N = 3) histopathological scoring. Inset and individual channels to show IDO1 expression in epithelial cells. Scale bars = 200 µm. **(H)** Shotgun metagenomic data, filtered for only the eubacteria kingdom, of AC biopsies from CD specimens with normal (N = 5) or active (N = 3) histopathological scoring. Data is represented as relative abundance of mapped species for individual specimens. “CD of SI” indicates AC specimens from CD patients with history of small intestinal inflammation and no history of AC inflammation, while “CD of SI and colon” indicates AC specimens from CD patients with a history of ileocolonic inflammation. p-value * < 0.05, ** < 0.01, *** < 0.001. **** < 0.0001. **See also Figure S12 and Table S8.**

To close the pathobiont-IDO1 mechanistic connection in human specimens, we performed shotgun metagenomics of AC biopsies from CD patients with normal AC and active AC inflammation by histology. While most DNA sequences mapped to the host, dozens of microbes were detected, with variable species abundances across specimens (Figure 7H, Figure S12I, Table S8). Rather than comparing the relative abundance of individual microbes, we focused on the presence or absence of microbes that are capable of replicating intracellularly and thus triggering IDO1 expression. Amongst specimens from patients with active AC inflammation, all samples tested were positive for *Staphylococcus aureus.* In contrast, *S. aureus* was detected in only one of five specimens from CD patients with normal AC histology. *S. aureus,* frequently detected in human stool samples, is a highly prevalent bacterium acting as a pathobiont that can cause life-threatening disease in specific tissues and in immunocompromised individuals, making it an interesting pathobiont to consider in CD pathogenesis.^84–87^ Our results demonstrate that like *Chlamydia*-driven upregulation of epithelial IDO1 in mouse models of CD, an analogous epithelial IDO1 upregulation in response to pathobionts is a hallmark of active CD in the AC. In addition to supporting the finding by Mishkin *et al.* that *C. muridarum* is a reemergent pathogen in murine research facilities, our work demonstrates how this bacterium induces chronic colonic inflammation in genetically susceptible hosts, supporting their notion that *C. muridarum* infection may confound experimental studies.^57^ Additionally, our work provides a unique preclinical animal model with inflammation targeted to the AC, a common site of CD involvement, that can be leveraged to develop new therapies.

## DISCUSSION

Over the last two decades, numerous biologic and small molecular therapies have been approved for the induction of remission of CD, none of which differentiate based on the specific regions of the GI tract affected.^88,89^ Though CD can affect any region along the gastrointestinal tract, it is most often localized to the TI or AC, and while these anatomical locations are contiguous tissues, they are distinct organs with unique functions and microenvironments. Here, we demonstrate that increased expression of IDO1 in secretory epithelial cells of the AC, but not in those of the TI, promotes chronic, pathobiont-triggered inflammatory disease in a region-specific manner. Given that some CD patients exhibit ileocolonic involvement, a compelling hypothesis is that inflammation-associated microbiota from the TI migrate downstream into the AC, triggering disease within the same susceptible individual through a similar mechanism. However, our results indicate pathobiont-driven AC inflammation is independent of upstream ileal disease. Moreover, depletion of Paneth cells, which produce antimicrobials that regulate the local and potentially downstream colonic microbiota, did not impact pathobiont-driven inflammation in the AC. Thus, CD is not simply a general inflammatory disease of the intestine, and disease etiology varies based on the unique cellular and molecular features specific to each affected region.

The anatomical site of disease development may depend on how each region responds to pathobionts and commensal microbes. Generally, microbial diversity and abundance increase along the length of the small intestine and colon, and the mucous layer becomes thicker and increasingly impenetrable to microbiota.^4–6,8^ Protection conferred by secretory cells, such as antimicrobial-secreting Paneth cells in the TI, help maintain host-microbe balance and prevent inflammatory disease. In the AC, however, there is an abrupt loss of Paneth cells and a sharp increase in bacterial load. Although goblet cells respond by increasing mucus secretion, the AC mucus layer is more penetrable compared to that of the distal colon.^4–8^ In our CD model, *C. muridarum* colonized surface epithelial cells along the entire length of the colon, but inclusions were markedly more prevalent in the ascending colon. The regional transition from TI to AC may create a vulnerable point in the mucosal barrier, potentially explaining why the AC is more susceptible to pathobiont-triggered inflammation.

Pathobionts are members of the stable microbiota that usually coexist with the host and become harmful under some conditions, potentially triggering CD.^27,55^ We identified *C. muridarum* as a pathobiont that is necessary and sufficient to drive chronic inflammation in the AC, inducing IDO1 in susceptible secretory epithelial cells. *Chlamydia* species are known to infect a wide range of host mucosal tissues and cause multiple diseases.^30^ Notably, specific *C. trachomatis* serovars are causative agents of proctocolitis that are commonly misdiagnosed as IBD.^29^ *Chlamydia* infection symptoms can vary from mild to severe, with some individuals remaining as long-term, asymptomatic carriers.^28^ In our CONV mouse colony, *C. muridarum* is part of the endogenous microbiota and is transmitted to offspring and cagemates. Wildtype mice are asymptomatic carriers, and our data in Tnf^ΔARE/+^ and IL10rb^-/-^ susceptible models indicates that *C. muridarum* is a pathobiont. While *Chlamydia* species were not detected in our relatively small cohort of patient AC biopsies, there’s emerging evidence of asymptomatic GI carriage in both humans and animals.^90^ Recent work demonstrated that *C. trachomatis* is frequently detected in the gastrointestinal tract and can infect colonic epithelial cells in patient-derived organoids.^91,92^ In our cohort of human specimens, *S. aureus* is prevalent in the AC mucosa of CD patients with a history of AC involvement and absent from the AC mucosa of CD patients with only TI or distal colon involvement. Both *Chlamydia* and *S. aureus* replicate inside host cells and their dormancy within epithelial cells of the gastrointestinal tract is speculated as a long-term reservoir for persistent colonization.^56,92–97^ Our results support that pathobionts persisting in colonic epithelial cells can induce chronic disease when encountering both anatomical and genetic susceptibilities.

Our results in the mouse and human demonstrate a mechanism by which epithelial cells upregulate IDO1 in response to the pathobionts to induce chronic inflammation of the AC. While we did not investigate the immune cascade downstream of IDO1 upregulation, we hypothesize increased conversion of tryptophan to kynurenine, coupled with increased TNF levels, promotes dysregulation of the regulatory T cell (Treg) compartment. While Tregs are known for suppressing an active immune response, elevated Tregs are frequently observed in inflamed IBD mucosa despite uncontrolled inflammation.^98,99^ Recent work demonstrated the emergence of proinflammatory Tregs in CD-associated inflammation and may arise from sustained TNF signaling.^100–102^ Adoptive transfer of Tregs, as an immunosuppressive strategy to combat an overactive immune response, is currently being investigated for therapeutic treatment of active CD.^103^ However, immune suppression may not be beneficial when considering pathobiont colonization, as this may thwart efforts by other immune cells to clear the microbe. Indeed, meta-analyses of patient adverse events indicate the most frequent consequences of immunosuppression with modern IBD therapies are opportunistic infections driven by pathobionts.^104^ Our work, together with recent findings of proinflammatory Treg function, suggests Treg therapy should be approached with caution for the treatment of CD. In contrast, our findings may warrant investigation and drug repurposing of IDO1 inhibitors, which have been in development for the treatment of various cancers, with none securing approval by the U.S. Food and Drug Administration (FDA).^105–107^ We envision that future CD therapies will balance the value of immunosuppression with the necessity of proper immune function to clear microbes, and additionally, will be tailored to regionalized features within the small intestine and colon.

## AUTHOR CONTRIBUTIONS

Conceptualization, P.N.S. and K.S.L.; data curation, P.N.S., E.P.S., A.J.S., K.T.W., L.A.C., Q.L., R.H.V., and K.S.L.; formal analysis, P.N.S., E.P.S., M.E.B., Y.Y., H.K., K.D.M., and K.S.L.; funding acquisition, N.O.M., K.T.W., L.A.C., J.A.G., Q.L., M.K.W., R.H.V., W.Z., and K.S.L.; investigation, P.N.S., J.W., E.P.S., L.S., A.J.S., T.K., W.K., M.E.B., Y.Y., H.K., Y.X., S.K., M.D.H., M.A.L., L.Z., D.A., N.T., K.D.M., O.S.K., M.H.H., J.R., J.L., A.B., M.K.W., and K.S.L.; methodology, P.N.S., E.P.S., L.S., A.J.S., M.E.B., Y.Y., H.K., M.D.H., K.D.M., J.A.G., M.K.W., R.H.V., W.Z., and K.S.L.; project administration, P.N.S., E.P.S., L.S., A.J.S., K.T.W., L.A.C., J.A.G., Q.L., M.K.W., R.H.V., W.Z., and K.S.L.; software, P.N.S., E.P.S., Y.Y., H.K., M.D.H., K.D.M., Q.L., and K.S.L.; resources; N.O.M., K.T.W., L.A.C., J.A.G., Q.L., M.K.W., R.H.V., W.Z., and K.S.L.; supervision, K.T.W., L.A.C., J.A.G., Q.L., M.K.W., R.H.V., W.Z., and K.S.L.; validation, B.C. and C.N.H.; P.N.S., J.W., E.P.S., K.D.M., R.H.V., and K.S.L.; visualization, P.N.S., J.W., E.P.S., L.S., M.E.B., Y.Y., H.K., S.K., M.A.L., K.D.M., and K.S.L.; writing – original draft, P.N.S. and K.S.L.

## ACKNOWLEDGEMENTS

The authors would like to thank Frank Revetta, Zhengyi Chen, Jacob M. Curry, Clara D. Si, and Jia A. Mei and Drs. Julia Drewes, Christopher Peritore-Galve, Borden Lacy, Cynthia Sears, James Cassat, Kara Eichelberger, Joseph Roland, Mirazul Islam, Judith Agudo, as well as the Vanderbilt Epithelial Biology Center for stimulating discussions centered around inflammation and the microbiome. We apologize to those we have failed to acknowledge due to space constraints. We would also like to thank Drs. Bob Coffey, George Kollias, Taha Jan, Peter Dempsey, Kevin Haigis, the Vanderbilt Gut Cell Atlas Program, and the European Mouse Mutant Archive for making critical mouse, human tissue, and microscopy resources available. The authors would also like to thank various cores and centers and their supporting grant at Vanderbilt: Translational Pathology Shared Resource (P30CA068485), Cell Imaging Shared Resource and Digital Histology Shared Resource (P30DK058404), VANTAGE Genomics Core, Division of Animal Care, and Vanderbilt Genome Editing Resource. This study was supported by NIH grants R01DK103831 and U54CA274367 (to K.S.L.), F31DK127687 (to P.N.S.), T32HD007502 (in support of P.N.S.), R01DK134692 and R35GM147470 (to W.Z.), R01DK128200 (to K.T.W.), R01AI173599 (to R.H.V.), HHMI Gilliam Fellowship GT16446 (to D.A. and K.S.L.), NSF GRF 2444112 (to M.E.B.), the Stanley Cohen Innovation Fund (to K.S.L.), The Leona M. and Harry B. Helmsley Charitable Trust G-1903-03793 (to the VUMC Gut Cell Atlas which K.S.L., L.A.C., K.T.W. are members), CCF SRA 1061046 (to J.A.G.), VA Merit grants I01CX002473 (to K.T.W.), I01CX002662 (to L.A.C.). Diagrams were created with images from <u>BioRender.com</u>.

## List of Tables

**Table S1.** Shotgun metagenomic data of ascending colon luminal contents from wildtype and Tnf^ΔARE/+^ mice in CONV and SPF-B facilities.

**Table S2.** Shotgun metagenomic data of ascending colon luminal contents of vehicle-treated and doxycycline-treated Tnf^ΔARE/+^ mice from the CONV facility.

**Table S3.** Shotgun metagenomic data of ascending colon luminal contents of sham-inoculated and CM001-GFP-inoculated Tnf^ΔARE/+^ mice from the SPF-B facility.

**Table S4.** Upregulated genes (logFC > 1) in each ascending colonic epithelial cell type from scRNA-seq data of wildtype and Tnf^ΔARE/+^ mice from the CONV and SPF-B facilities.

**Table S5.** Facility-specific upregulated genes (logFC > 1.5, pooled wildtype and Tnf^ΔARE/+^ samples) in each ascending colon epithelial cell type from scRNA-seq data.

**Table S6.** Facility-specific upregulated genes (logFC > 1.5, only young Tnf^ΔARE/+^ samples) in each ascending colon epithelial cell type from scRNA-seq data.

**Table S7.** Overrepresentation analysis (ORA) results from enriched gene sets specific to CONV and SPF-B facilities.

**Table S8.** Shotgun metagenomic data of ascending colon biopsies from human specimens.

**Table S9.** Patient metadata for shotgun metagenomics, immunofluorescence imaging, and scRNA-seq specimens.

**Table S10.** NCBI accession numbers for species and strains used in pangenomic analysis.

## STARIZMETHODS

### RESOURCE AVAILABILITY

#### Lead contact

Further information and requests for resources and reagents should be directed to and will be fulfilled by Lead Contact: Ken S. Lau, PhD at

#### Materials availability

Contact: Ken S. Lau, PhD at

#### Data and code availability

Not yet deposited.

*Any additional information required to reanalyze the data reported in this paper is available from the lead contact upon request*.

### KEY RESOURCES TABLE

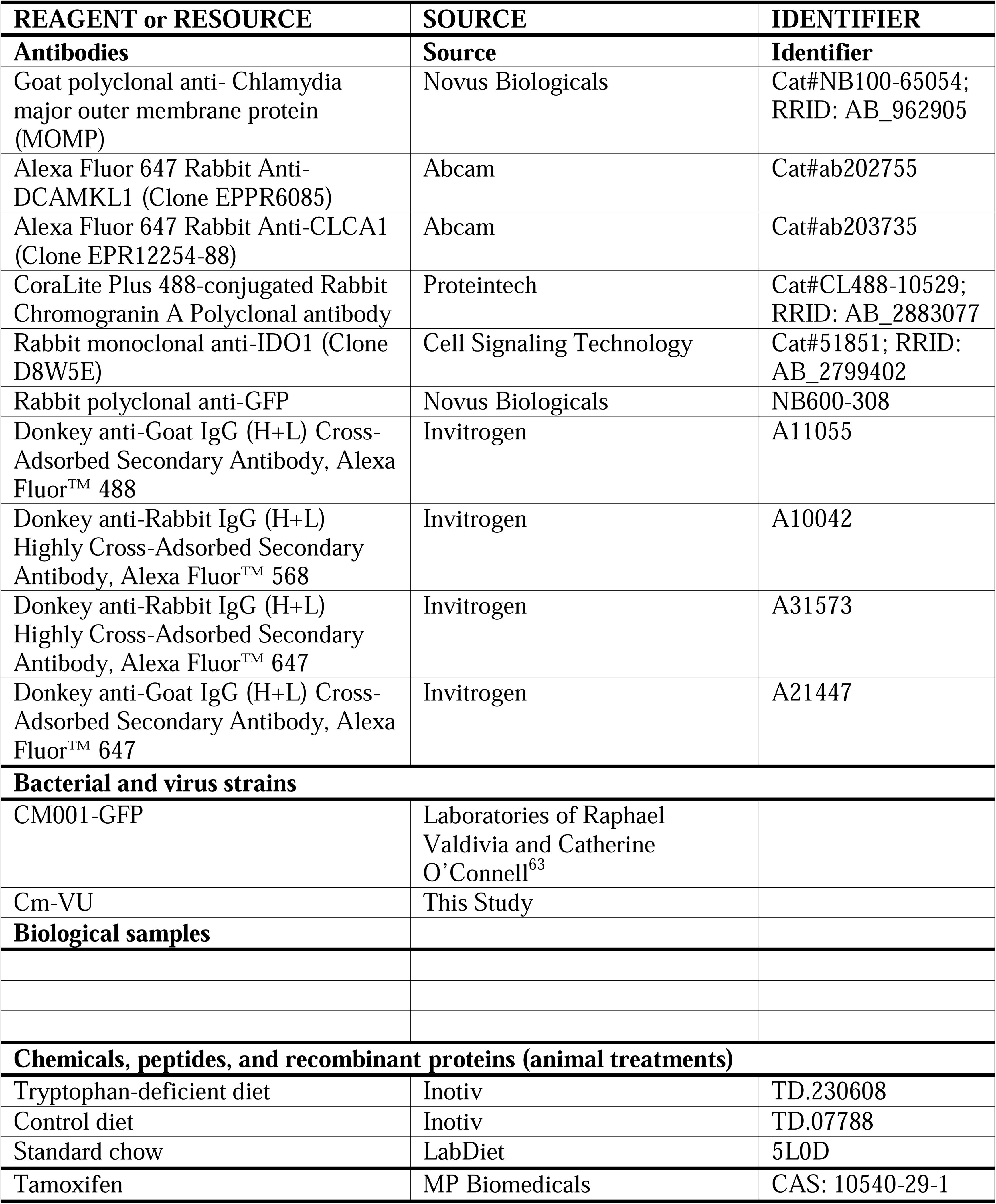
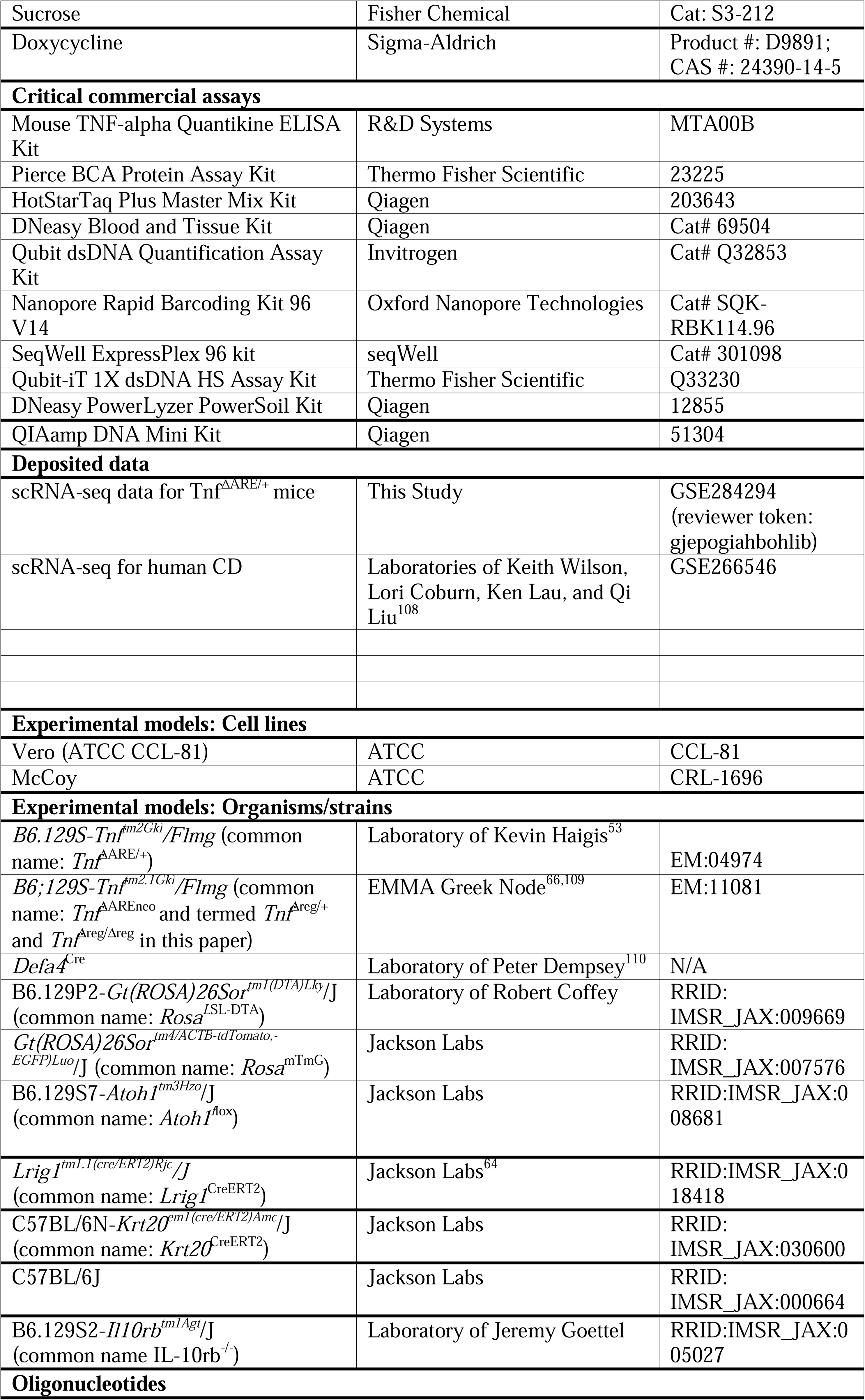
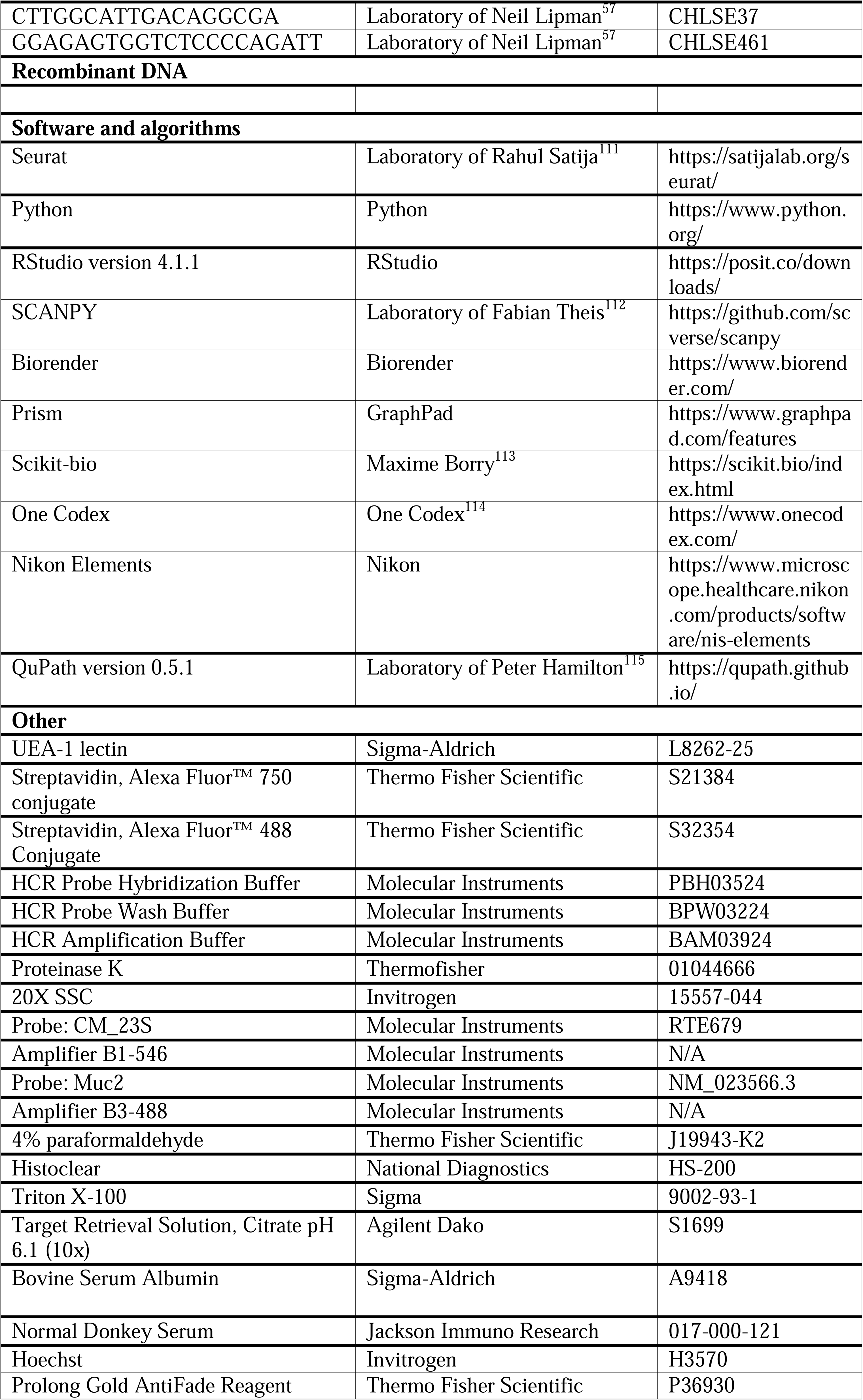
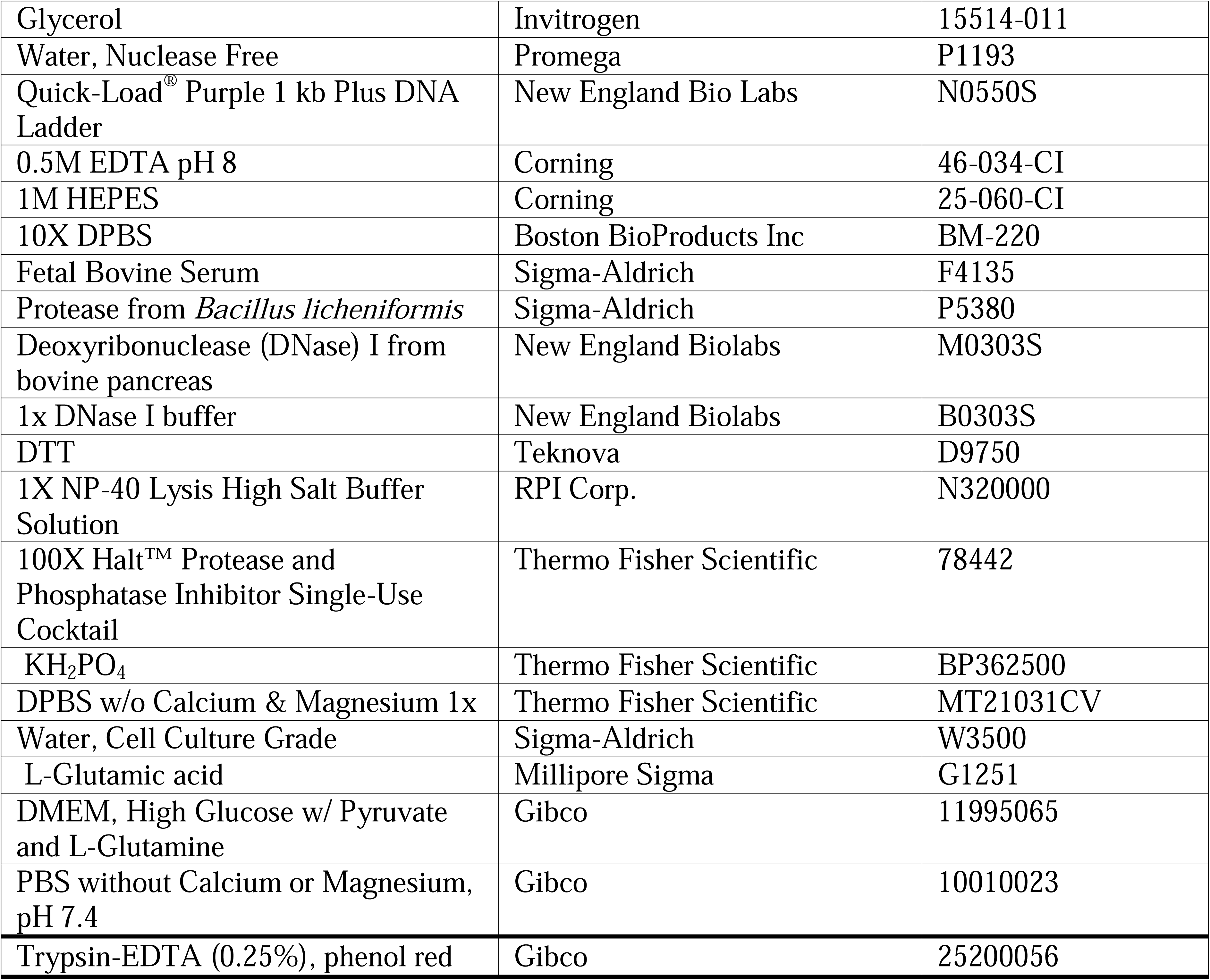

### EXPERIMENTAL MODEL AND STUDY PARTICIPANT DETAILS

#### Human Specimens from the Gut Cell Atlas

The Gut Cell Atlas (GCA) study protocol was approved by the Institutional Review Board at Vanderbilt University Medical Center (IRB #191738). Written informed consent was obtained from non-IBD control and CD subjects to obtain TI and AC tissues at the time of scheduled endoscopic procedures. All samples were obtained as a part of the clinical trial “Combinatorial Single Cell Strategies for a Crohn’s Disease Gut Cell Atlas”, identifier NCT04113733 (<u>clinicaltrials.gov</u>).

Between December 2019 and July 2023, endoscopy subjects were prospectively recruited in the IBD clinic or GI endoscopy unit at Vanderbilt University Medical Center prior to colonoscopy for CD disease activity assessment or non-IBD indications including colorectal cancer screening or polyp surveillance. Exclusion criteria for the study were: pregnancy, known coagulopathy or bleeding disorders, known renal or hepatic impairment, history of organ transplantation, or inability to give informed consent. For all participants, demographics including age, gender, medical history, and medication use were determined from participant reporting and review of the electronic medical record. Patient metadata is available in Table S9.

Tissue biopsies in the TI and AC for immediate scRNA-seq data generation were placed in chelation buffer (4 mM EDTA, 0.5 mM DTT in PBS) as previously described.^83^ Tissue biopsies for later DNA extraction were flash frozen on dry ice and stored at -80°C. Tissue biopsies for formalin-fixed and paraffin-embedded (FFPE) tissue block generation underwent standard fixation and paraffin embedding protocols. 5 µm sections were used from each FFPE block, stained with hematoxylin and eosin (H&E), and examined in a blinded manner by a gastrointestinal pathologist (M.K.W.) and graded accordingly as: inactive (normal_CD or quiescent) or active (mild, moderate, or severe activity). All associated study data were collected and managed using Research Electronic Data Capture (REDCap) electronic data capture tools hosted at Vanderbilt^116,117^, including Clinical Data Interoperability Services, such as Clinical Data Pull^118^ and e-consent.^119^

#### Murine Husbandry and Housing Conditions

All animal experiments were performed under protocols approved by the Vanderbilt University Animal Care and Use Committee and in accordance with NIH guidelines. Mice were housed (up to 5 per cage) in a conventional (CONV), specific pathogen-free barrier (SPF-B), or ABSL-2 environment under a standard 12-hour daylight cycle, and were fed a standard irradiated chow (PicoLab® Laboratory Rodent Diet 5L0D) and provided water ad libitum, unless otherwise specified. Littermate controls of both sexes were used for experiments when possible. Housing facility conditions are detailed below.

#### CONV Murine Housing Facility

Mice were procured from approved commercial vendors or non-commercial sources that meet facility health requirements after a period of quarantine and testing. Mice were housed in individual ventilated cages. Acidified municipal water is provided via automatic watering valves. Traffic flow of animals was two-way (i.e., mice may leave the facility for procedures and return to the housing room). PPE required to work in Level 5 rooms in this facility includes gowns and gloves. Excluded pathogens: Mouse Hepatitis Virus, Mouse Parvovirus, Minute Virus of Mice, Lymphocytic Choriomeningitis Virus, Sendai Virus, Pneumonia Virus of Mice, Epizootic Diarrhea of Infant Mice, Theiler’s Mouse Encephalomyelitis Virus, Mouse Pox, Mouse Adenovirus, Mouse Reovirus, *Mycoplasma pulmonis*, *Syphacia* spp, *Aspiculuris* spp, *Myobia musculi*, *Radfordia affinis*, *Myocoptes musculinus*, *Psorergates simplex*.

#### SPF-B Murine Housing Facility

Mice were procured only from approved commercial vendors or via rederivation. Mice were housed in autoclaved individual ventilated cages. Autoclaved reverse osmosis water was provided in water bottles. The SPF-B facility may not be entered after working in any other mouse facility. Traffic flow of animals is one way from the facility (i.e., mice may not return if they leave the facility for procedures). All equipment and supplies must be dedicated to this facility. PPE required to work in the facility includes gowns, gloves, caps, masks, and shoe covers. Excluded pathogens include those in the CONV facility as well as mouse Norovirus and *Helicobacter* spp.

#### Animal Biosafety Level 2 (ABSL-2) Murine Housing Facility

Rooms were dedicated for mice experimentally infected with agents that require ABSL-2 housing conditions. Mice were purchased directly from approved vendors and delivered to ABSL-2 housing rooms or transferred from other SPF-B housing rooms. Mice were housed in individual ventilated cages and provided reverse osmosis water in bottles. All handling of animals was conducted in biosafety cabinets and animals were not transferred to other housing areas from ABSL-2 rooms. PPE required to work in ABSL-2 areas includes gowns, double gloves, and masks.

### METHOD DETAILS

#### Murine models and experiments

Tnf^ΔARE/+^ and wildtype (Tnf^+/+^) mice were housed in CONV, SPF-B, or ABSL-2 facilities as described above. Mice were maintained on a C57BL/6J background by backcrossing to breeders purchased from Jackson Labs. Genotyping of the mice was performed through Transnetyx (Cordova, TN, USA). Other alleles on C57BL/6J backgrounds were bred into the Tnf^ΔARE/+^ mouseline as described below. Weights were monitored to ensure mice were euthanized at humane endpoints. Unless otherwise specified, equal representation of male and female mice were used in all experiments wherever possible.

Genetic backgrounds of the mice were tested using the miniMUGA array performed through Transnetyx (Cordova, TN, USA).^120^

For co-housing experiments, SPF-B wildtype and Tnf^ΔARE/+^ mice were transferred to the CONV facility and placed into fresh cages with CONV facility, *Chlamydia muridarum-*positive wildtype or Tnf^ΔARE/+^ cagemates. This experiment was performed at various ages, with same-sex or mixed-sex conditions, and harvested at various aged, as indicated in figure legends. If transferred as pups, the SPF-B litter was combined with a CONV wildtype or Tnf^ΔARE/+^ dam with similar age litter and were monitored. Genotyping for transgenes or mutated alleles was used to distinguish CONV and SPF-B pups in the mixed litters. Mice were euthanized and tissue was harvested at timepoints and ages indicated in figure legends.

For isolated caging experiments, adult SPF-B wildtype and Tnf^ΔARE/+^ mice were transferred to the CONV facility and placed into fresh cages without CONV facility cagemates. Mice and cage changes were handled with fresh gloves, a clean hood, and clean materials to minimize transfer of microbiota from CONV facility mice. Mice were euthanized and tissue was harvested at timepoints and ages indicated in figure legends.

For doxycycline treatment experiments, adult *C. muridarum*-positive CONV mice were given vehicle (5% sucrose) or doxycycline (5% sucrose, 2 mg/mL doxycycline) water ad libitum for 1-2 weeks, as indicated in figure legends. To minimize re-inoculation with *C. muridarum* from fecal matter, cages were changed at the start of the experiment and upon cessation of vehicle or doxycycline treatment. Mice and cage changes were handled with fresh gloves, a clean hood, and clean materials to minimize transfer of microbiota from other cages. Mice were treated at various ages as stated in figure legends. Fecal samples were collected throughout the experiment including before treatment, after cessation of treatment, and at harvest. Mice were euthanized and tissue was harvested at timepoints and ages indicated in figure legends.

For CM001-GFP association experiments, adult SPF-B mice were transferred to the ABSL-2 facility. Mice and cage changes were handled with fresh gloves, a clean hood, and clean materials to minimize transfer of microbiota from other cages. Mice were administered a single treatment of sham or CM001-GFP inoculate via oral gavage in 100 µL volume. CM001-GFP (3×10^6^ IFUs) inoculate was prepared from CM001-GFP-infected Vero cell lysate and sham inoculate was prepared from uninfected Vero cell lysate, both prepared in 1X SPG (sucrose-phosphate-glutamate) buffer (219 mM sucrose, 3.7 mM KH_2_PO_4_, 4.9 mM L-glutamic acid in cell-grade water to pH 7.4-7.6 using NaOH). Fecal samples were collected throughout the experiment including before treatment, at timepoints post-treatment, and at harvest. Mice were euthanized and tissue was harvested at timepoints and ages indicated in figure legends.

For Lrig1-Atoh1-KO Tnf^ΔARE/+^ experiments, tamoxifen was administered to SPF-B mice in experimental (*Lrig1^CreERT2/+^;Atoh1^fl/fl^;Tnf*^ΔARE/+^) or control (*Lrig1^+/+^;Atoh1^fl/fl^;Tnf*^ΔARE/+^) conditions via intraperitoneal injection of 2 mg of tamoxifen for 4 consecutive days (8 mg total dose). Seven days after the first injection, mice were transferred to the CONV facility and co-housed with *C. muridarum-*positive CONV Tnf^ΔARE/+^ cagemates. Mice were euthanized and tissue was harvested at timepoints and ages indicated in figure legends.

For Krt20-Atoh1-KO Tnf^ΔARE/+^ experiments, tamoxifen was administered to CONV mice in experimental (*Krt20^CreERT2/+^;Atoh1^fl/fl^;Tnf*^ΔARE/+^) or control (*Krt20^+/+^;Atoh1^fl/fl^;Tnf*^ΔARE/+^) conditions via two intraperitoneal injections per week of 6.25 mg of tamoxifen for 3 weeks (37.5 mg total dose). Mice were euthanized and tissue was harvested three weeks after the first injection at ages indicated in figure legends.

For tryptophan-deficient diet experiments, CONV Tnf^ΔARE/+^ mice were given synthetic diets that were tryptophan-deficient or 0.18% tryptophan (control) ad libitum. Mice were euthanized and tissue was harvested after four weeks of treatment at ages indicated in figure legends.

For rederivation of the Tnf^Δreg^ line in *C. muridarum-*free CONV facility conditions, *in vitro* fertilization was performed using Tnf^Δreg/+^ sperm and C57BL/6J eggs. Embryos were transferred into superovulating dams. Weanlings were transferred to the CONV facility and kept in isolated caging conditions, as described above.

For PC-DTA Tnf^ΔARE/+^ experiments, CONV mice of experimental (*Defa4^Cre/+^;Rosa^LSL-DTA/+^;Tnf*^ΔARE/+^) or control (*Defa4^+/+^;Rosa^LSL-DTA/+^;Tnf*^ΔARE/+^ or *Defa4^+/+^;Rosa^+/+^;Tnf*^ΔARE/+^) genotypes were euthanized and tissue was harvested at ages indicated in figure legends.

For PC-Atoh1-KO Tnf^ΔARE/+^ experiments, CONV mice of experimental (*Defa4^Cre/+^;Atoh1^fl/fl^;Tnf*^ΔARE/+^) or control (*Defa4^+/+^;Atoh1^fl/+^;Tnf*^ΔARE/+^, *Defa4^Cre/+^;Atoh1^fl/+^;Tnf*^ΔARE/+^, or *Defa4^+/+^;Atoh1^fl/fl^;Tnf*^ΔARE/+^) genotypes were euthanized and tissue was harvested at ages indicated in figure legends.

For IL-10rb^-/-^ experiments, *C. muridarum-*free IL-10rb^-/-^ CONV mice were either kept in isolated caging (control condition) or co-housed with *C. muridarum-*positive Tnf^ΔARE/+^ cagemates (experimental condition) for 4-8 weeks. Mice were euthanized and tissue was harvested at ages indicated in figure legends.

#### DNA extraction and shotgun metagenomic sequencing of murine intestinal luminal contents and human AC biopsies

Mouse colons were harvested and the proximal colon, defined by the first third, was isolated. The contents of the proximal colon were flushed using sterile PBS pH 7.4 without Calcium or Magnesium into individual barcoded tubes containing DNA stabilization buffer to ensure reproducibility, stability, and traceability, and shipped for DNA extraction, library preparation, and sequencing by Transnetyx (Cordova, TN USA). DNA extraction was optimized and fully automated using a robust process for reproducible extraction of inhibitor-free, high molecular weight genomic DNA that captures the true microbial diversity of samples.

Human AC biopsies were thawed at room temperature and DNA was extracted using the QIAamp DNA Mini Kit (Qiagen). Extracted DNA concentrations were measured using the Qubit-iT 1X dsDNA HS Assay Kit (Thermo Fisher Scientific). Extracted DNA was frozen and sent for library preparation and sequencing by Transnetyx (Cordova, TN USA). Transnetyx (Cordova, TN, USA).

Mouse and human DNA extractions were subjected to quality control (QC) steps. Next, genomic DNA was converted into sequencing libraries using a method optimized for minimal bias. Unique dual indexed (UDI) adapters were used to ensure that reads and/or organisms are not mis-assigned. After QC, the libraries were sequenced using the shotgun sequencing method (a depth of 2 million 2×150 bp read pairs), which enables species and strain level taxonomic resolution.

#### Shotgun metagenomic sequencing analysis

Sequencing data were uploaded automatically onto One Codex analysis software and analyzed against the One Codex database consisting of ∼148K complete microbial reference genomes.^114^ The classification results were filtered through several statistical post-processing steps designed to eliminate false positive results caused by contamination or sequencing artifacts. Host reads were identified using mouse and human genomes. The relative abundance of each microbial species is estimated based on the depth and coverage of sequencing across every available reference genome.

#### Murine scRNA-seq data generation

Colons were dissected and trisected. The proximal colon, defined by the first third of the colon, was isolated and luminal content was flushed using PBS without calcium or magnesium. Next, the tissue was flayed open, washed twice in PBS without calcium and magnesium, and placed into cold chelation buffer consisting of 3 mM EDTA (Corning), 20 mM HEPES (Corning), and 0.5 mM DTT (Teknova) in PBS without calcium or magnesium for 1 hour at 4°C. Chelation buffer was discarded and colonic epithelium (crypts) were collected by vigorously shaking the colonic tissue in PBS. The shaking process was repeated for a total of four times and the pooled crypt isolates were passed through a 100 µm filter. Crypts were dissociated into single cells using cold protease buffer consisting of 5 mg/mL Protease from *Bacillus licheniformis* (Sigma-Aldrich) and 2.5 mg/mL DNase (Sigma-Aldrich) in PBS without calcium or magnesium on a rotator for 25 minutes at 4°C. After dissociation, the suspension was mechanically dissociated into single cells by gentle pipetting with a wide bore pipette tip. The solution was passed through a 70 µm filter and resultant single cells were quenched with 2% FBS. A series of washes were performed to obtain a single-cell suspension with mostly live cells and minimal debris. Single-cell encapsulation, library preparation, and sequencing were performed as previously described.^121,122^

#### scRNA-seq data alignment and filtering

For human scRNA-seq data, alignment and filtering were described previously.^83^ For murine scRNA-seq data, raw sequencing data were aligned using dropEst to mouse GRCm38.85 resulting in count matrices.^123^ Resultant count matrices were subjected to filtering protocols as follows. First, cells were ranked according to number of transcripts (counts) detected and an inflection point cutoff was used to filter out cells with low counts. Next, cells were projected in 2D space using UMAP embedding and overlays of total counts, percent mitochondrial reads, and cell type specific markers were used to iteratively remove low-quality cells.

#### scRNA-seq data analysis

Analysis of pre-processed murine scRNA-seq data was carried out Seurat version 4.0.4 as described previously.^111,124^ Functions use default arguments unless specified. Batch effects were minimal as seen from the intermixing of cell types, and therefore, no batch corrections were performed. Briefly, Seurat’s clustering (FindClusters) function with iterative adjustment of the clustering resolution parameter was used along with differential expression analysis (FindMarkers) to label cell types based on marker gene expression. Differential expression analysis (FindMarkers) was performed to identify differences between cell types and across conditions, using log fold change of 1 or 1.5, only positive (upregulated) genes, and all other standard parameters. Over-representation analysis (ORA) was performed using WebGestalt web-based tool with the geneontology/biological process functional database.^125^

Analysis of pre-processed human scRNA-seq data was carried out in SCANPY with UMAP embeddings and cell type annotations as described previously.^83,112^

#### Murine intestinal tissue fixation, embedding, and staining

Intestinal tissue was dissected, flushed with PBS, flayed longitudinally onto Whatman filter paper, and fixed in 4% PFA (Thermo Scientific) for approximately 24 hrs at room temperature. Fixed tissues were washed with PBS, swiss-rolled, stabilized in 2% agar, and stored in 70% EtOH until processing and paraffin embedding to generate formalin-fixed, paraffin embedded (FFPE) blocks. FFPE blocks were sectioned at 5 µm thick onto charged glass slides. Slides were incubated in Histoclear (National Diagnostics) solution for deparaffinization. Next, slides were rehydrated by stepped incubation in 90%, 70%, and 50% ethanol solutions, followed by permeabilization in 0.3% Triton X-100 (Sigma). Next, slides were antigen retrieved in citrate buffer (Dako) for 20 minutes in a pressure cooker at 105°C followed by a 20-minute bench cool down. For IF staining, slides were blocked for 1+ hour in a humidified chamber in blocking buffer consisting of 2.5% Normal Donkey Serum, 1% BSA, 0.3% Triton X-100, 1:10,000 Hoechst 33342 in PBS prior to antibody staining. Primary antibodies were incubated on the slides in a humidified chamber for 1+ hour, followed by three washes in PBS. Compatible conjugated secondaries, if applicable, were incubated on the slides in a humidified chamber for 1 hour, followed by three washes in 1X PBS. Slides were mounted in Prolong Gold mounting media (Thermo Fisher Scientific) or 50% glycerol in PBS. For histological analysis, slides were processed and stained for hematoxylin and eosin using standard approaches. For RNA-FISH, Molecular Instruments HCR™ RNA-FISH protocol was adapted as described previously^83^ and used with custom probes designed for *Muc2* or *C. muridarum 23S* RNA on FFPE tissue sections.^126^

#### Brightfield and fluorescence imaging

Brightfield whole slide images were imaged on a Leica Aperio AT2 automated slide scanner (Leica Biosystems) at 20x magnification to a resolution of 0.273 µm /pixel. Immunofluorescent tissue sections for whole slide imaging were imaged on a Leica Aperio Versa automated slide scanner (Leica Biosystems) at 20x magnification to a resolution of 0.325 µm/pixel utilizing a combination of five filter cubes at six wavelengths (405, 488, 546, 594, 647, 750 nm). Spinning disk confocal microscopy was performed using a Nikon Ti2 inverted light microscope with a Yokogawa CSU-X1 spinning disk head, an Andor DU-897 EMCCD camera, and four excitation LASERs (405, 488, 561, and 647nm), a Plan Apo Lambda 20x/0.75 NA air objective, and an Apo TIRF 100x/1.49NA oil immersion objective. High-magnification images were deconvolved (Richardson-Lucy deconvolution of image volumes, 20 iterations) using Nikon Elements software.

#### Image-based quantification of *C. muridarum*-infected surface epithelium

Quantification of *C. muridarum*-infected colonic epithelia surface was done utilizing QuPath. Proximal and distal regions of the colon were drawn in as an annotation, where the proximal region was defined as the portion with proximal folds. Next, a thresholder was created utilizing the nuclei signal (Hoechst) channel to detect cells. Cells positive or negative for *C. muridarum* infection, based on signal for *Chlamydia* major outer membrane protein (MOMP), were quantified by thresholding. The surface epithelia was annotated by manual drawing and was then measured as a distance. Finally, the proportion of infected epithelium was quantified by dividing the percent of MOMP-positive cells by the surface epithelium distance.

#### Histopathological Scoring

Inflammation and damage in H&E stained FFPE slides was quantified by a trained pathologist in a blinded fashion using methods adapted from Erben *et al.,* 2014.^127^ A Colitis Score was calculated based on the sum of four subscores that each range from 0-4, with 0 as normal and 4 as most pronounced: lamina propria (LP) chronic inflammation, LP polymorphonuclear leukocytes (PMNs), depth of inflammation, and crypt inflammation.

#### *In vitro* bone marrow-derived macrophage (BMDM) stimulation

Bone marrow was flushed from tibia and femur bones of wildtype, Tnf^Δreg/+^, Tnf^ΔARE/+^, and Tnf^Δreg/Δreg^ mice in an aseptic environment. Cell clumps were disaggregated gently, and the cell suspension was centrifuged at 250xg for 5 min. The cell pellets were resuspended in DMEM medium with 20% FBS, 100 IU/ml penicillin, 100 μg/ml streptomycin and 30% L cell-conditioned medium, and cultured at 37°C in 5% CO_2_. For generation of M1 macrophages, BMDM were stimulated for 24 hr with 100 ng/mL of LPS and 20 ng/mL IFN-γ. After 6–7 days, nonadherent cells were aspirated and adherent macrophages were removed by washing plate with ice-cold PBS and scraping.

#### TNF enzyme-linked immunosorbent assay (ELISA)

To quantify TNF levels in stimulated BMDMs from wildtype, Tnf^Δreg/+^, Tnf^ΔARE/+^, and Tnf^Δreg/Δreg^ mice, TNF ELISA was performed on stimulated BMDM lysates by adding equal volumes of each sample to individual wells of the Mouse TNF-alpha Quantikine ELISA Kit (R&D Systems) and following the manufacturer’s protocol.

To quantify TNF levels in intestinal epithelial cells of CONV wildtype and Tnf^ΔARE/+^ mice, small intestine was dissected and trisected (duodedum, jejunum, ileum) and colon was dissected and bisected (proximal colon and distal colon). Luminal content was flushed using PBS without calcium or magnesium. Next, the tissue was flayed open, washed twice in PBS without calcium and magnesium, and placed into cold chelation buffer consisting of 3 mM EDTA (Corning), 20 mM HEPES (Corning), and 0.5 mM DTT (Teknova) in PBS without calcium or magnesium for 1 hour at 4°C. Chelation buffer was discarded and colonic epithelium (crypts) were collected by vigorously shaking the colonic tissue in PBS. The shaking process was repeated for a total of four times and the pooled crypt isolates were passed through a 100 µm filter. The dissociated crypts were instead lysed in NP-40 Lysis Buffer (RPI Corp.) with Protease and 1X Phosphatase Inhibitor Cocktail (Thermo Fisher Scientific) combined with brief sonication. Lysates were then quantified for total protein level using the Pierce BCA Protein Assay Kit (Thermo Fisher Scientific), diluted to a standard protein concentration, and TNF ELISA was performed by adding samples to individual wells of the Mouse TNF Quantikine ELISA Kit (R&D Systems) and following the manufacturer’s protocol.

#### Fecal DNA testing for *C. muridarum* 23S rRNA

Fecal pellets were collected and frozen on dry ice immediately. Samples were thawed and DNA was extracted using the DNeasy PowerLyzer PowerSoil Kit (Qiagen) according to manufacturer’s instructions A region of *C. muridarum* 23S rRNA gene was amplified by PCR using HotStarTaq Plus Master Mix Kit (Qiagen) with the primers and thermocycling conditions developed by Mishkin *et al.,* 2022, which generated an expected product of 425 bp^57^ as determined by gel electrophoresis.

#### Cell lines and bacterial strains

*C. muridarum* strains CM001 (a gift from Catherine O’Connell^63^), CM001-GFP, and Cm-VU were propagated in Vero cells (CCL-81; ATCC). The Vero cells were maintained in Dulbecco’s Modified Eagle Medium (DMEM; Gibco) supplemented with 10% fetal bovine serum (FBS; Sigma-Aldrich). Confluent Vero cells were infected with *C. muridarum* by centrifugation at 1500 × g for 30 minutes at 10°C. *C. muridarum* strains were harvested by water lysis and sonication. Concentrations were determined using tittering protocols described previously.^128^

#### *Chlamydia* shuttle vector cloning and transformation

The *Chlamydia* plasmid p2TK2_spec_-Nigg-GFP was constructed by replacing *mCherry* in the p2TK2_spec_-Nigg mCh(Gro_L2_) vector^129^ with *gfp*. The *gfp* gene was amplified from pGFP::Nigg^130^ using the primers GFP_gro_inf_F (*Sac* I): 5’-AGCTTAAACgagctcATGAGTAAAGGAGAAGCACT-3’ and GFP_gro_inf_R (*Kpn* I): 5’-ggtaccTTACTTGTATAGTTCATCCATGCCATGTG-3’. Additionally, the *groESL* promoter was amplified from p2TK2_spec_-Nigg mCh(Gro_L2_) using the primers groPro_inf_F (*Age* I): 5’-ACCGTATTACaccggtATTTTTAAAAATAGCAGTTG-3’ and groPro_inf_R (*Sac* I): 5’-ggtaccTTACTTGTATAGTTCATCCATGCCATGTG-3’. The *groESL* terminator was amplified from the same vector using the primers groTerm_inf_F (*Kpn* I): 5’-TACAAGTAAggtaccTTCCTCTAATGGGAACAAATAG-3’ and groTerm_inf_R (*Not* I): 5’-TCCGTCGACgcggccgcAGAAAAGGATGGTCGTAA-3’. The vector p2TK2_spec_-Nigg mCh(Gro_L2_) was digested with *Age* I and *Not* I, and the three PCR products were combined and ligated into the digested vector using In-Fusion Cloning (Takara Bio). The resulting plasmid, p2TK2_spec_-Nigg-GFP, was sequence verified by Sanger sequencing using the groPro_inf_F and the groTerm_inf_R primers at Eton Biosciences.

*C. muridarum* CM001 was transformed with p2TK2_spec_-Nigg-GFP as follows: 1×10^7^ IFUs were incubated with 15 µg of plasmid DNA in buffer containing 0.9 mM calcium chloride for 40 minutes, then added to confluent Vero cells in a 6-well plate. Sixteen hours post-infection, 500 µg /ml of spectinomycin was added. Lysates from infected cells were used sequentially to infect new monolayers every 48 hours until GFP-positive inclusions were visible. Transformants were plaque-purified to obtain a clonal isolate.

#### Cm-VU isolation

Whole colon was isolated from a *C. muridarum*-positive *Tnf*^ΔARE/+^ mouse from the CONV facility and placed in ice-cold 1X SPG (sucrose-phosphate-glutamate) buffer (219 mM sucrose, 3.7 mM KH_2_PO_4_, 4.9 mM L-glutamic acid in cell-grade water to pH 7.4-7.6 using NaOH). All steps were performed on ice or at 4°C. After flushing out fecal material with DPBS (Gibco), the colon was filleted, minced into 2 mm pieces, and washed with DPBS. Epithelial cells were isolated by incubating the tissue in 8 mM EDTA-DPBS for 75 minutes with rotation, followed by three DPBS washes. Cells were further dislodged by vigorous shaking in 0.1% BSA-DPBS, and the supernatant was filtered through a 70 µm filter. The filtrate was centrifuged at 300 ×g for 5 minutes, and the pellet was resuspended in water, sonicated, and filtered through a 0.45 µm filter. The *Chlamydi*a-containing filtrate was added to Vero cells. After 18 hours, 0.5 µg/ml cycloheximide and 50 µg/ml gentamicin were added. Bacteria were passaged as necessary.

#### Whole genome sequencing (WGS) of *C. muridarum*

*C. muridarum* isolates Cm-VU and CM001 were prepared from six 6-well plates of Vero cells infected at an MOI of 2 and grown for 48 hours. Cells were lysed in sterile water, sonicated, and centrifuged at 500 ×g for 5 minutes at 4°C to remove host cell debris. The supernatant was then centrifuged at 19,000 ×g for 5 minutes at 4°C to pellet *Chlamydia* elementary bodies (EBs), which were treated with 8 U of DNase I (New England Biolabs) in 1x DNase I buffer (New England Biolabs) for 1 hour at 37°C to degrade host DNA. EBs were washed in DPBS, and genomic DNA was extracted using the DNeasy Blood and Tissue Kit (Qiagen), following the manufacturer’s protocol, quantified with the Qubit dsDNA Quantification Assay Kit (Invitrogen), and assessed for the quality of high molecular weight DNA by agarose gel electrophoresis.

Cm*-*VU was sequenced by Plasmidsaurus (https://www.plasmidsaurus.com) using their Hybrid Bacterial Genome workflow, combining Oxford Nanopore and Illumina sequencing. An amplification-free library was prepared using the Oxford Nanopore Rapid Barcoding Kit 96 V14 (Cat# SQK-RBK114.96) and sequenced on a PromethION P24 instrument with a R10.4.1 flow cell. Basecalling was performed in super-accurate mode using ont-doradod-for promethion v7.1.4, with a minimum Qscore of 10, and adapters were trimmed using MinKnow. The Illumina library was prepared using the SeqWell ExpressPlex 96 kit (Cat# 301098) and sequenced on an Illumina NextSeq2000 instrument with paired-end 2×150bp reads. Nanopore fastq reads were first processed using filtlong (v0.2.1), retaining reads > 1000 bp, and subsampled to 100x coverage using rasusa (v2.0.0).^131^ Assembly was performed with Flye (v2.9.4).^132,133^ The assembly was polished using Polypolish (v0.6.0)^134,135^ with Illumina reads, resulting in a 1,072,053 bp chromosome and 7,501 bp plasmid.

*C. muridarum* CM001 was sequenced by the Microbial Sequencing and Analysis Center, now SeqCenter in Pittsburgh, PA (https://www.seqcenter.com). The library was prepared using the Illumina DNA Prep kit with IDT 10bp UDI indices and sequenced on an Illumina NextSeq 2000, producing 2×151bp reads. Demultiplexing and adapter trimming was preformed using bcl-convert (v3.9.3).^136^ The *C. muridarum* CM001 genome was assembled in Geneious Prime (v2022.1). Fastq reads were trimmed using BBDuk, normalized using BBNorm, paired reads were merged with BBMerge, and duplicate reads were removed using Dedupe. The processed reads were aligned to the published CM001 reference genome (NZ_CP027217) ^63^ using Bowtie2, and the resulting consensus sequence was used for pangenomic analysis.

#### Pangenomic analysis

*Chlamydia* genomes (n=13) and one *Chlamydiifrater phoenicopteri* genome were retrieved from NCBI (Table S10). A comparison of 15 *Chlamydia* genomes including Cm-VU and CM001 was performed using anvi’o version 7.1.^137^ To satisfy anvi’o formatting requirements, the fasta files containing all genomes were converted into contigs-fasta format using anvi-script-reformat-fasta with parameters --simplify_names and --seq_type NT. The reformatted genomes were then used to generate databases using the command anvi-gen-contigs-database. These databases were annotated using anvi-run-hmms and anvi-run-cogs^138^ and used to generate a genomes storage database using anvi-gen-genomes-storage. The pangenome was constructed using anvi-pan-genome with parameters --minbit 0.5, --mcl-inflation 10, and --use-ncbi-blast. Average nucleotide identity between all pairs of genomes was calculated using anvi-compute-genome-similarity using --pyANI program^139^ after including *C. phoenicopteri* as an outgroup. ANI calculations were run once each using the ANIb and TETRA alignment methods. The pangenome figure was finalized using Inkscape.^140^ The ANI dendrogram was visualized using the Interactive Tree of Life ^141^.

#### Genomic sequencing and comparison of native *C. muridarum* strain (Cm-VU)

Genome comparisons between reference *Chlamydia* genomes and the *Chlamydia* isolate Cm-VU were conducted using average nucleotide identity (ANI) and tetranucleotide frequency correlation coefficient (TETRA) analyses, with species-level cutoffs set at >95% for ANI and >0.989 for TETRA.^142–144^ The genome of the Cm-VU isolate was sequenced and compared to 14 other *Chlamydia* genomes, including 6 *C. trachomatis* strains, 3 *C. muridarum* strains (including CM001, which was used for experiments in this study), and reference genomes from 5 other *Chlamydia* species. All genomes, except for CM001 and Cm-VU, were acquired from publicly available sequences on NCBI (Table S10).

## REFERENCES

1. Burclaff, J. et al. A Proximal-to-Distal Survey of Healthy Adult Human Small Intestine and Colon Epithelium by Single-Cell Transcriptomics. Cell. Mol. Gastroenterol. Hepatol. 13, 1554–1589 (2022).

2. Zwick, R. K. et al. Epithelial zonation along the mouse and human small intestine defines five discrete metabolic domains. Nat. Cell Biol. 26, 250–262 (2024).

3. Lentle, R. G. & Janssen, P. W. M. The Physical Processes of Digestion. Phys. Process. Dig. (2011). doi:10.1007/978-1-4419-9449-3

4. Bergstrom, K. et al. Proximal colon-derived O-glycosylated mucus encapsulates and modulates the microbiota. Science (80-. ). 370, (2020).

5. Kamphuis, J. B. J., Mercier-Bonin, M., Eutamène, H. & Theodorou, V. Mucus organisation is shaped by colonic content; a new view. Sci. Reports 2017 71 7, 1–13 (2017).

6. Johansson, M. E. V., Holmén Larsson, J. M. & Hansson, G. C. The two mucus layers of colon are organized by the MUC2 mucin, whereas the outer layer is a legislator of host-microbial interactions. Proc. Natl. Acad. Sci. U. S. A. 108, 4659–4665 (2011).

7. Johansson, M. E. V. et al. The inner of the two Muc2 mucin-dependent mucus layers in colon is devoid of bacteria. Proc. Natl. Acad. Sci. U. S. A. 105, 15064–15069 (2008).

8. Ermund, A., Gustafsson, J. K., Hansson, G. C. & Keita, Å. V. Mucus properties and goblet cell quantification in mouse, rat and human ileal Peyer’s patches. PLoS One 8, (2013).

9. Roda, G. et al. Crohn’s disease. Nat. Rev. Dis. Prim. 2020 61 6, 1–19 (2020).

10. Noble, A. J., Nowak, J. K., Adams, A. T., Uhlig, H. H. & Satsangi, J. Defining Interactions Between the Genome, Epigenome, and the Environment in Inflammatory Bowel Disease: Progress and Prospects. Gastroenterology 165, 44–60.e2 (2023).

11. Kaser, A., Zeissig, S. & Blumberg, R. S. Inflammatory Bowel Disease. Annu. Rev. Immunol. 28, 573 (2010).

12. Khor, B., Gardet, A. & Xavier, R. J. Genetics and pathogenesis of inflammatory bowel disease. Nature 474, 307– 317 (2011).

13. Torres, J., Mehandru, S., Colombel, J. F. & Peyrin-Biroulet, L. Crohn’s disease. Lancet (London, England) 389, 1741–1755 (2017).

14. Jostins, L. et al. Host–microbe interactions have shaped the genetic architecture of inflammatory bowel disease. Nat. 2012 4917422 491, 119–124 (2012).

15. Xavier, R. J. & Podolsky, D. K. Unravelling the pathogenesis of inflammatory bowel disease. Nature 448, 427– 434 (2007).

16. Silverberg, M. S. et al. Toward an integrated clinical, molecular and serological classification of inflammatory bowel disease: report of a Working Party of the 2005 Montreal World Congress of Gastroenterology. Can. J. Gastroenterol. 19 Suppl A, (2005).

17. Cosnes, J., Gowerrousseau, C., Seksik, P. & Cortot, A. Epidemiology and natural history of inflammatory bowel diseases. Gastroenterology 140, 1785–1794.e4 (2011).

18. Ng, S. C. et al. Incidence and phenotype of inflammatory bowel disease based on results from the Asia-pacific Crohn’s and colitis epidemiology study. Gastroenterology 145, 158–165.e2 (2013).

19. Park, S. J., Kim, W. H. & Cheon, J. H. Clinical characteristics and treatment of inflammatory bowel disease: a comparison of Eastern and Western perspectives. World J. Gastroenterol. 20, 11525–11537 (2014).

20. Richard, N. et al. Crohn’s disease: Why the ileum? World J. Gastroenterol. 29, 3222 (2023).

21. El Hadad, J., Schreiner, P., Vavricka, S. R. & Greuter, T. The Genetics of Inflammatory Bowel Disease. Mol. Diagn. Ther. 28, 27 (2023).

22. Ntunzwenimana, J. C. et al. Functional screen of inflammatory bowel disease genes reveals key epithelial functions. Genome Med. 13, 1–21 (2021).

23. Odenwald, M. A. & Turner, J. R. The intestinal epithelial barrier: A therapeutic target? Nature Reviews Gastroenterology and Hepatology 14, 9–21 (2017).

24. Schaubeck, M. et al. Dysbiotic gut microbiota causes transmissible Crohn’s disease-like ileitis independent of failure in antimicrobial defence. Gut 65, 225–237 (2016).

25. Bamias, G. et al. Commensal Bacteria Exacerbate Intestinal Inflammation but Are Not Essential for the Development of Murine Ileitis. J. Immunol. 178, 1809–1818 (2007).

26. Sellon, R. K. et al. Resident Enteric Bacteria Are Necessary for Development of Spontaneous Colitis and Immune System Activation in Interleukin-10-Deficient Mice. Infect. Immun. 66, 5224 (1998).

27. Mazmanian, S. K., Round, J. L. & Kasper, D. L. A microbial symbiosis factor prevents intestinal inflammatory disease. Nature 453, 620–625 (2008).

28. Organization, W. H. Report on global sexually transmitted infection surveillance 2018. South. Med. J. 70, 01–58 (2018).

29. Stoner, B. P. & Cohen, S. E. Lymphogranuloma Venereum 2015: Clinical Presentation, Diagnosis, and Treatment. Clin. Infect. Dis. 61, S865–S873 (2015).

30. Elwell, C., Mirrashidi, K. & Engel, J. Chlamydia cell biology and pathogenesis. Nat. Rev. Microbiol. 2016 146 14, 385–400 (2016).

31. Stelzner, K., Vollmuth, N. & Rudel, T. Intracellular lifestyle of Chlamydia trachomatis and host–pathogen interactions. Nat. Rev. Microbiol. 2023 217 21, 448–462 (2023).

32. Bastidas, R. J., Elwell, C. A., Engel, J. N. & Valdivia, R. H. Chlamydial Intracellular Survival Strategies. Cold Spring Harb. Perspect. Med. 3, a010256 (2013).

33. Cocchiaro, J. L. & Valdivia, R. H. New insights into Chlamydia intracellular survival mechanisms. Cell. Microbiol. 11, 1571–1578 (2009).

34. Fehlner-Gardiner, C. et al. Molecular basis defining human Chlamydia trachomatis tissue tropism. A possible role for tryptophan synthase. J. Biol. Chem. 277, 26893–26903 (2002).

35. Taylor, M. W. & Feng, G. Relationship between interferon-gamma, indoleamine 2,3-dioxygenase, and tryptophan catabolism. FASEB J 5, 2516–2522 (1991).

36. Feng, G. S. & Taylor, M. W. Interferon gamma-resistant mutants are defective in the induction of indoleamine 2,3-dioxygenase. Proc. Natl. Acad. Sci. U. S. A. 86, 7144–7148 (1989).

37. Roshick, C., Wood, H., Caldwell, H. D. & McClarty, G. Comparison of Gamma Interferon-Mediated Antichlamydial Defense Mechanisms in Human and Mouse Cells. Infect. Immun. 74, 225 (2006).

38. Schoborg, R. V. Chlamydia persistence – a tool to dissect chlamydia–host interactions. Microbes Infect. 13, 649 (2011).

39. Ibana, J. A. et al. Inhibition of Indoleamine 2,3-Dioxygenase Activity by Levo-1-Methyl Tryptophan Blocks Gamma Interferon-Induced Chlamydia trachomatis Persistence in Human Epithelial Cells. Infect. Immun. 79, 4425 (2011).

40. Alvarado, D. M. et al. Epithelial IDO1 Modulates AHR and Notch Signaling to Increase Differentiation of Secretory Cells and Alter Mucus-Associated Microbiota. Gastroenterology 157, 1093 (2019).

41. Mezrich, J. D. et al. An interaction between kynurenine and the aryl hydrocarbon receptor can generate regulatory T cells. J. Immunol. 185, 3190–3198 (2010).

42. Yan, Y. et al. IDO upregulates regulatory T cells via tryptophan catabolite and suppresses encephalitogenic T cell responses in experimental autoimmune encephalomyelitis. J. Immunol. 185, 5953–5961 (2010).

43. Baban, B. et al. IDO activates regulatory T cells and blocks their conversion into TH17-like T cells. J. Immunol. 183, 2475 (2009).

44. Ciorba, M. A. Indoleamine 2,3 dioxygenase (IDO) in Intestinal Disease. Curr. Opin. Gastroenterol. 29, 146 (2013).

45. Ferdinande, L. et al. Inflamed Intestinal Mucosa Features a Specific Epithelial Expression Pattern of Indoleamine 2,3-Dioxygenase. 10.1177/039463200802100205 21, 289–295 (2008).

46. Barceló-Batllori, S. et al. Proteomic analysis of cytokine induced proteins in human intestinal epithelial cells: Implications for inflammatory bowel diseases. Proteomics 2, 551–560 (2002).

47. Dieckgraefe, B. K., Stenson, W. F., Korzenik, J. R., Swanson, P. E. & Harrington, C. A. Analysis of mucosal gene expression in inflammatory bowel disease by parallel oligonucleotide arrays. Physiol. Genomics 4, 1–11 (2000).

48. Braegger, C. P., Nicholls, S., Murch, S. H., MacDonald, T. T. & Stephens, S. Tumour necrosis factor alpha in stool as a marker of intestinal inflammation. Lancet 339, 89–91 (1992).

49. Breese, E. J. et al. Tumor necrosis factor α-producing cells in the intestinal mucosa of children with inflammatory bowel disease. Gastroenterology 106, 1455–1466 (1994).

50. MacDonald, T. T., Hutchings, P., Choy, M. Y., Murch, S. & Cooke, A. Tumour necrosis factor-alpha and interferon-gamma production measured at the single cell level in normal and inflamed human intestine. Clin. Exp. Immunol. 81, 301–305 (1990).

51. Reinecker, H. C. et al. Enhanced secretion of tumour necrosis factor-alpha, IL-6, and IL-1 beta by isolated lamina propria mononuclear cells from patients with ulcerative colitis and Crohn’s disease. Clin. Exp. Immunol. 94, 174–181 (1993).

52. Nicholls, S., Stephens, S., Braegger, C. P., Walker-Smith, J. A. & MacDonald, T. T. Cytokines in stools of children with inflammatory bowel disease or infective diarrhoea. J. Clin. Pathol. 46, 757–760 (1993).

53. Kontoyiannis, D., Pasparakis, M., Pizarro, T. T., Cominelli, F. & Kollias, G. Impaired on/off regulation of TNF biosynthesis in mice lacking TNF AU-rich elements: implications for joint and gut-associated immunopathologies. Immunity 10, 387–98 (1999).

54. Schaubeck, M. et al. Dysbiotic gut microbiota causes transmissible Crohn’s disease-like ileitis independent of failure in antimicrobial defence. Gut 65, 225–237 (2016).

55. Caruso, R., Lo, B. C., Chen, G. Y. & Núñez, G. Host-pathobiont interactions in Crohn’s disease. Nat. Rev. Gastroenterol. Hepatol. (2024). doi:10.1038/S41575-024-00997-Y

56. Yeruva, L., Spencer, N., Bowlin, A. K., Wang, Y. & Rank, R. G. Chlamydial infection of the gastrointestinal tract: a reservoir for persistent infection. Pathog. Dis. 68, 88 (2013).

57. Mishkin, N. et al. Reemergence of the Murine Bacterial Pathogen Chlamydia muridarum in Research Mouse Colonies. Comp. Med. 72, 230–242 (2022).

58. Manam, S. et al. Tumor Necrosis Factor (TNF) Receptor Superfamily Member 1b on CD8+ T Cells and TNF Receptor Superfamily Member 1a on Non-CD8+ T Cells Contribute Significantly to Upper Genital Tract Pathology Following Chlamydial Infection. J. Infect. Dis. 211, 2014–2022 (2015).

59. Zafiratos, M. T. et al. Tumor necrosis factor receptor superfamily members 1a and 1b contribute to exacerbation of atherosclerosis by Chlamydia pneumoniae in mice. Microbes Infect. 21, 104–108 (2019).

60. RECOMMENDATIONS FOR TREATMENT OF CHLAMYDIAL INFECTIONS. (2016).

61. Palleja, A. et al. Recovery of gut microbiota of healthy adults following antibiotic exposure. Nat. Microbiol. 2018 311 3, 1255–1265 (2018).

62. Anthony, W. E. et al. Acute and persistent effects of commonly used antibiotics on the gut microbiome and resistome in healthy adults. Cell Rep. 39, (2022).

63. Poston, T. B. et al. T Cell-Independent Gamma Interferon and B Cells Cooperate To Prevent Mortality Associated with Disseminated Chlamydia muridarum Genital Tract Infection. Infect. Immun. 86, (2018).

64. Powell, A. E. et al. The pan-ErbB negative regulator lrig1 is an intestinal stem cell marker that functions as a tumor suppressor. Cell 149, 146–158 (2012).

65. Herring, C. A. et al. Unsupervised Trajectory Analysis of Single-Cell RNA-Seq and Imaging Data Reveals Alternative Tuft Cell Origins in the Gut. Cell Syst. 6, 37–51.e9 (2018).

66. Roulis, M., Armaka, M., Manoloukos, M., Apostolaki, M. & Kollias, G. Intestinal epithelial cells as producers but not targets of chronic TNF suffice to cause murine Crohn-like pathology. Proc. Natl. Acad. Sci. U. S. A. 108, 5396–5401 (2011).

67. Meyer, K., Gellhorn, A., Prudden, J. F., Lehman, W. L. & Steinberg, A. Lysozyme in Chronic Ulcerative Colitis.∗. 10.3181/00379727-65-15917P 65, 221–222 (1947).

68. Meyer, K., Gellhorn, A., Prudden, J. F., Lehman, W. L. & Steinberg, A. Lysozyme activity in ulcerative alimentary disease. II. Lysozyme activity in chronic ulcerative colitis. Am. J. Med. 5, 496–502 (1948).

69. Yu, S. et al. Paneth Cell Multipotency Induced by Notch Activation following Injury. Cell Stem Cell 23, 46–59.e5 (2018).

70. Roulis, M. et al. Host and microbiota interactions are critical for development of murine Crohn’s-like ileitis. (2016). doi:10.1038/mi.2015.102

71. Perminow, G. et al. Defective Paneth Cell—Mediated Host Defense in Pediatric Ileal Crohn’s Disease. Am. J. Gastroenterol. 105, 452–459 (2010).

72. Wehkamp, J. & Stange, E. F. Paneth’s disease. J. Crohn’s Colitis 4, 523–531 (2010).

73. Liu, T.-C. et al. Paneth cell defects in Crohn’s disease patients promote dysbiosis. JCI Insight 1, e86907 (2016).

74. Kühn, R., Löhler, J., Rennick, D., Rajewsky, K. & Müller, W. Interleukin-10-deficient mice develop chronic enterocolitis. Cell 75, 263–274 (1993).

75. Spencer, S. D. et al. The orphan receptor CRF2-4 is an essential subunit of the interleukin 10 receptor. J. Exp. Med. 187, 571–578 (1998).

76. Glocker, E.-O. et al. Inflammatory Bowel Disease and Mutations Affecting the Interleukin-10 Receptor. N. Engl. J. Med. 361, 2033 (2009).

77. Kotlarz, D. et al. Loss of interleukin-10 signaling and infantile inflammatory bowel disease: implications for diagnosis and therapy. Gastroenterology 143, 347–355 (2012).

78. Noguchi, E., Homma, Y., Kang, X., Netea, M. G. & Ma, X. A Crohn’s disease-associated NOD2 mutation suppresses transcription of human IL10 by inhibiting activity of the nuclear ribonucleoprotein hnRNP-A1. Nat. Immunol. 10, 471–479 (2009).

79. Burich, A. et al. Helicobacter-induced inflammatory bowel disease in IL-10- and T cell-deficient mice. Am. J. Physiol. Gastrointest. Liver Physiol. 281, (2001).

80. Dieleman, L. A. et al. Helicobacter hepaticus does not induce or potentiate colitis in interleukin-10-deficient mice. Infect. Immun. 68, 5107–5113 (2000).

81. Moran, J. P., Walter, J., Tannock, G. W., Tonkonogy, S. L. & Sartor, R. B. Bifidobacterium animalis causes extensive duodenitis and mild colonic inflammation in monoassociated interleukin-10-deficient mice. Inflamm. Bowel Dis. 15, 1022–1031 (2009).

82. Kim, S. C., Tonkonogy, S. L., Karrasch, T., Jobin, C. & Balfour Sartor, R. Dual-association of gnotobiotic IL-10-/- mice with 2 nonpathogenic commensal bacteria induces aggressive pancolitis. Inflamm. Bowel Dis. 13, 1457–1466 (2007).

83. Li, J. et al. Identification and multimodal characterization of a specialized epithelial cell type associated with Crohn’s disease. Nat. Commun. 2024 151 15, 1–19 (2024).

84. Rimland, D. & Roberson, B. Gastrointestinal carriage of methicillin-resistant Staphylococcus aureus. J. Clin. Microbiol. 24, 137–138 (1986).

85. Ray, A. J., Pultz, N. J., Bhalla, A., Aron, D. C. & Donskey, C. J. Coexistence of vancomycin-resistant enterococci and Staphylococcus aureus in the intestinal tracts of hospitalized patients. Clin. Infect. Dis. 37, 875–881 (2003).

86. Boyce, J. M., Havill, N. L. & Maria, B. Frequency and possible infection control implications of gastrointestinal colonization with methicillin-resistant Staphylococcus aureus. J. Clin. Microbiol. 43, 5992–5995 (2005).

87. Acton, D. S., Tempelmans Plat-Sinnige, M. J., Van Wamel, W., De Groot, N. & Van Belkum, A. Intestinal carriage of Staphylococcus aureus: How does its frequency compare with that of nasal carriage and what is its clinical impact? Eur. J. Clin. Microbiol. Infect. Dis. 28, 115–127 (2009).

88. Lichtenstein, G. R. et al. ACG Clinical Guideline: Management of Crohn’s Disease in Adults. Am. J. Gastroenterol. 113, 481–517 (2018).

89. Gordon, H. et al. ECCO Guidelines on Therapeutics in Crohn’s Disease: Medical Treatment. J. Crohn’s Colitis 18, 1531–1555 (2024).

90. Rank, R. G. & Yeruva, L. Hidden in plain sight: chlamydial gastrointestinal infection and its relevance to persistence in human genital infection. Infect. Immun. 82, 1362–1371 (2014).

91. Hovhannisyan, P. et al. Infection of human organoids supports an intestinal niche for Chlamydia trachomatis. bioRxiv (2024).

92. Zhong, G. Chlamydia overcomes multiple gastrointestinal barriers to achieve long-lasting colonization. Trends Microbiol. 29, 1004 (2021).

93. Gresham, H. D. et al. Survival of Staphylococcus aureus inside neutrophils contributes to infection. J. Immunol. 164, 3713–3722 (2000).

94. Clement, S. et al. Evidence of an intracellular reservoir in the nasal mucosa of patients with recurrent Staphylococcus aureus rhinosinusitis. J. Infect. Dis. 192, 1023–1028 (2005).

95. Hayes, S. M. et al. Intracellular residency of Staphylococcus aureus within mast cells in nasal polyps: A novel observation. J. Allergy Clin. Immunol. 135, 1648–1651.e5 (2015).

96. Hanssen, A. M. et al. Localization of Staphylococcus aureus in tissue from the nasal vestibule in healthy carriers. BMC Microbiol. 17, (2017).

97. Flaxman, A. et al. Development of persistent gastrointestinal S. aureus carriage in mice. Sci. Reports 2017 71 7, 1– 13 (2017).

98. Saruta, M. et al. Characterization of FOXP3+CD4+ regulatory T cells in Crohn’s disease. Clin. Immunol. 125, 281–290 (2007).

99. Maul, J. et al. Peripheral and intestinal regulatory CD4+CD25high T cells in inflammatory bowel disease. Gastroenterology 128, 1868–1878 (2005).

100. Hovhannisyan, Z., Treatman, J., Littman, D. R. & Mayer, L. Characterization of IL-17-producing regulatory T cells in inflamed intestinal mucosa from patients with inflammatory bowel diseases. Gastroenterology 140, 957 (2010).

101. Kosinsky, R. L. et al. The FOXP3+ Pro-Inflammatory T Cell: A Potential Therapeutic Target in Crohn’s Disease. Gastroenterology 166, 631–644.e17 (2024).

102. Martin, J. C. et al. Single-Cell Analysis of Crohn’s Disease Lesions Identifies a Pathogenic Cellular Module Associated with Resistance to Anti-TNF Therapy. Cell 178, 1493 (2019).

103. Clough, J. N., Omer, O. S., Tasker, S., Lord, G. M. & Irving, P. M. Regulatory T-cell therapy in Crohn’s disease: challenges and advances. Gut 69, 942–952 (2020).

104. Wang, K., Zhu, Y., Liu, K., Zhu, H. & Ouyang, M. Adverse events of biologic or small molecule therapies in clinical trials for inflammatory bowel disease: A systematic review and meta-analysis. Heliyon 10, e25357 (2024).

105. Van Den Eynde, B. J., Van Baren, N. & Baurain, J. F. Is There a Clinical Future for IDO1 Inhibitors after the Failure of Epacadostat in Melanoma? Annu. Rev. Cancer Biol. 4, 241–256 (2020).

106. Tang, K., Wu, Y. H., Song, Y. & Yu, B. Indoleamine 2,3-dioxygenase 1 (IDO1) inhibitors in clinical trials for cancer immunotherapy. J. Hematol. Oncol. 14, 1–21 (2021).

107. Prendergast, G. C., Malachowski, W. P., DuHadaway, J. B. & Muller, A. J. Discovery of IDO1 inhibitors: from bench to bedside. Cancer Res. 77, 6795 (2017).

108. Li, J. et al. Identification and multimodal characterization of a specialized epithelial cell type associated with Crohn’s disease. Nat. Commun. 2024 151 15, 1–19 (2024).

109. Kontoyiannis, D., Pasparakis, M., Pizarro, T. T., Cominelli, F. & Kollias, G. Impaired on/off regulation of TNF biosynthesis in mice lacking TNF AU-rich elements: implications for joint and gut-associated immunopathologies. Immunity 10, 387–98 (1999).

110. Burger, E. et al. Loss of Paneth Cell Autophagy Causes Acute Susceptibility to Toxoplasma gondii-Mediated Inflammation. Cell Host Microbe 23, 177–190.e4 (2018).

111. Butler, A., Hoffman, P., Smibert, P., Papalexi, E. & Satija, R. Integrating single-cell transcriptomic data across different conditions, technologies, and species. Nat. Biotechnol. 2018 365 36, 411–420 (2018).

112. Wolf, F. A., Angerer, P. & Theis, F. J. SCANPY: Large-scale single-cell gene expression data analysis. Genome Biol. 19, 1–5 (2018).

113. Borry, M. Sourcepredict: Prediction of metagenomic sample sources using dimension reduction followed by machine learning classification. J. Open Source Softw. 4, 1540 (2019).

114. Minot, S. S., Krumm, N. & Greenfield, N. B. One Codex: A Sensitive and Accurate Data Platform for Genomic Microbial Identification. *bioRxiv* 027607 (2015). doi:10.1101/027607

115. Bankhead, P. et al. QuPath: Open source software for digital pathology image analysis. Sci. Reports 2017 71 7, 1– 7 (2017).

116. Harris, P. A. et al. Research electronic data capture (REDCap)--a metadata-driven methodology and workflow process for providing translational research informatics support. J. Biomed. Inform. 42, 377–381 (2009).

117. Harris, P. A. et al. The REDCap consortium: Building an international community of software platform partners. J. Biomed. Inform. 95, 103208 (2019).

118. Cheng, A. C. et al. REDCap on FHIR: Clinical Data Interoperability Services. J. Biomed. Inform. 121, (2021).

119. Lawrence, C. E. et al. A REDCap-based model for electronic consent (eConsent): Moving toward a more personalized consent. *J*. Clin. Transl. Sci. 4, 345–353 (2020).

120. Sigmon, J. S. et al. Content and Performance of the MiniMUGA Genotyping Array: A New Tool To Improve Rigor and Reproducibility in Mouse Research. Genetics 216, 905–930 (2020).

121. Banerjee, A. et al. Succinate Produced by Intestinal Microbes Promotes Specification of Tuft Cells to Suppress Ileal Inflammation. Gastroenterology 159, 2101–2115.e5 (2020).

122. Chen, B. et al. Differential pre-malignant programs and microenvironment chart distinct paths to malignancy in human colorectal polyps. Cell 184, 6262–6280.e26 (2021).

123. Petukhov, V. et al. dropEst: pipeline for accurate estimation of molecular counts in droplet-based single-cell RNA-seq experiments. Genome Biol. 19, 78 (2018).

124. Vega, P. N. et al. Cancer-Associated Fibroblasts and Squamous Epithelial Cells Constitute a Unique Microenvironment in a Mouse Model of Inflammation-Induced Colon Cancer. Front. Oncol. 12, (2022).

125. Elizarraras, J. M. et al. WebGestalt 2024: faster gene set analysis and new support for metabolomics and multi-omics. Nucleic Acids Res. 52, W415–W421 (2024).

126. Choi, H. M. T. et al. Third-generation in situ hybridization chain reaction: Multiplexed, quantitative, sensitive, versatile, robust. Dev. 145, (2018).

127. Erben, U. et al. A guide to histomorphological evaluation of intestinal inflammation in mouse models. Int. J. Clin. Exp. Pathol. 7, 4557 (2014).

128. Dolat, L. et al. Chlamydia repurposes the actin-binding protein EPS8 to disassemble epithelial tight junctions and promote infection. Cell Host Microbe 30, 1685 (2022).

129. Cortina, M. E., Clayton Bishop, R., DeVasure, B. A., Coppens, I. & Derre, I. The inclusion membrane protein IncS is critical for initiation of the Chlamydia intracellular developmental cycle. PLoS Pathog. 18, (2022).

130. Skilton, R. J. et al. The Chlamydia muridarum plasmid revisited[: new insights into growth kinetics. Wellcome Open Res. 3, 25 (2018).

131. Hall, M. B. Rasusa: Randomly subsample sequencing reads to a specified coverage. J. Open Source Softw. 7, 3941 (2022).

132. Kolmogorov, M., Yuan, J., Lin, Y. & Pevzner, P. A. Assembly of long, error-prone reads using repeat graphs. Nat. Biotechnol. 2019 375 37, 540–546 (2019).

133. Lin, Y. et al. Assembly of long error-prone reads using de Bruijn graphs. Proc. Natl. Acad. Sci. U. S. A. 113, E8396–E8405 (2016).

134. Wick, R. R. & Holt, K. E. Polypolish: Short-read polishing of long-read bacterial genome assemblies. PLoS Comput. Biol. 18, (2022).

135. Bouras, G. et al. How low can you go? Short-read polishing of Oxford Nanopore bacterial genome assemblies. Microb. Genomics 10, 001254 (2024).

136. BCL Convert. Available at: https://support-docs.illumina.com/SW/BCL_Convert/Content/SW/FrontPages/BCL_Convert.htm. (Accessed: 15th August 2024)

137. Eren, A. M. et al. Community-led, integrated, reproducible multi-omics with anvi’o. Nat. Microbiol. 6, 3–6 (2021).

138. Eddy, S. R. Accelerated Profile HMM Searches. PLOS Comput. Biol. 7, e1002195 (2011).

139. Pritchard, L., Glover, R. H., Humphris, S., Elphinstone, J. G. & Toth, I. K. Genomics and taxonomy in diagnostics for food security: soft-rotting enterobacterial plant pathogens. Anal. Methods 8, 12–24 (2015).

140. IW, D. Inkscape - Draw Freely. Available at: https://inkscape.org/. (Accessed: 15th August 2024)

141. Letunic, I. & Bork, P. Interactive Tree Of Life (iTOL) v5: an online tool for phylogenetic tree display and annotation. Nucleic Acids Res. 49, W293–W296 (2021).

142. Richter, M. & Rosselló-Móra, R. Shifting the genomic gold standard for the prokaryotic species definition. Proc. Natl. Acad. Sci. U. S. A. 106, 19126–19131 (2009).

143. Goris, J. et al. DNA-DNA hybridization values and their relationship to whole-genome sequence similarities. Int. J. Syst. Evol. Microbiol. 57, 81–91 (2007).

144. Pillonel, T., Bertelli, C., Salamin, N. & Greub, G. Taxogenomics of the order Chlamydiales. Int. J. Syst. Evol. Microbiol. 65, 1381–1393 (2015).

