## Supplementary material for "Pathobiont-triggered induction of epithelial IDO1 drives regional susceptibility to Inflammatory Bowel Disease": Data S1

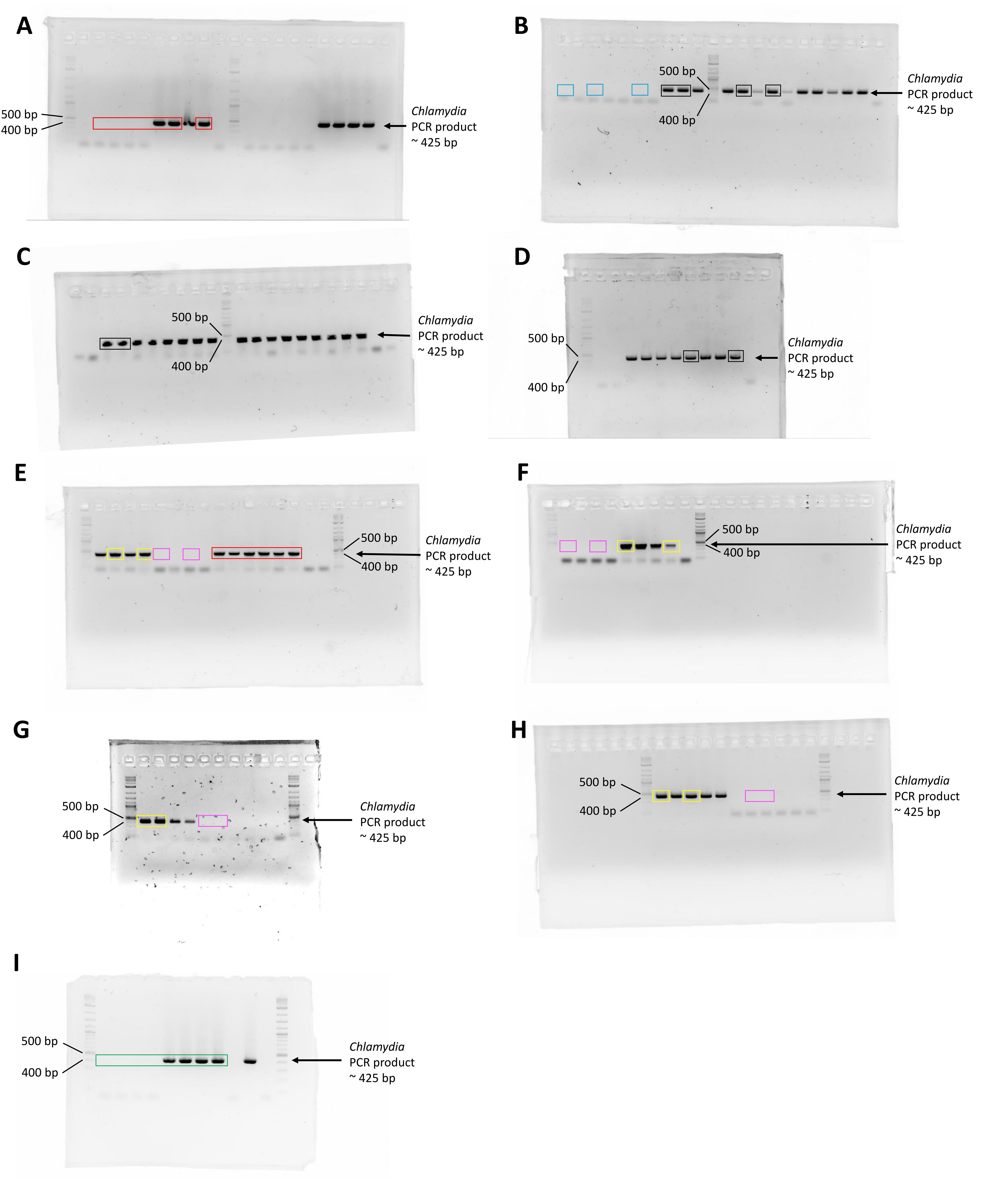


**Data S1. Full PCR gels for *Chlamydia.***

A) Red boxed areas are cropped regions in Figure 2D.

B) Blue boxed areas are cropped regions in Figure 2E before co-housing condition. Black boxed areas are cropped regions in Figure 3B vehicle pre-treatment condition.

C) Black boxed areas are cropped regions in Figure 3B doxycycline pre-treatment condition.

D) Black boxed areas are cropped regions in Figure 3B doxycycline pre-treatment condition.

E) Red boxed areas are cropped regions in Figure 2E post co-housing condition. Yellow boxed areas are cropped regions in Figure 3B vehicle post-treatment condition. Magenta boxed areas are cropped regions in Figure 3B doxycycline post-treatment condition.

F) Yellow boxed areas are cropped regions in Figure 3B vehicle post-treatment condition. Magenta boxed areas are cropped regions in Figure 3B doxycycline post-treatment condition.

G) Yellow boxed areas are cropped regions in Figure 3B vehicle harvest condition. Magenta boxed areas are cropped regions in Figure 3B doxycycline harvest condition.

H) Yellow boxed areas are cropped regions in Figure 3B vehicle harvest condition. Magenta boxed areas are cropped regions in Figure 3B doxycycline harvest condition.

I) Green boxed areas are cropped regions in Figure 3K.

**Related to Figure 2 and Figure 3.**
