## Supplemental Figures for "Pathobiont-triggered induction of epithelial IDO1 drives regional susceptibility to Inflammatory Bowel Disease"

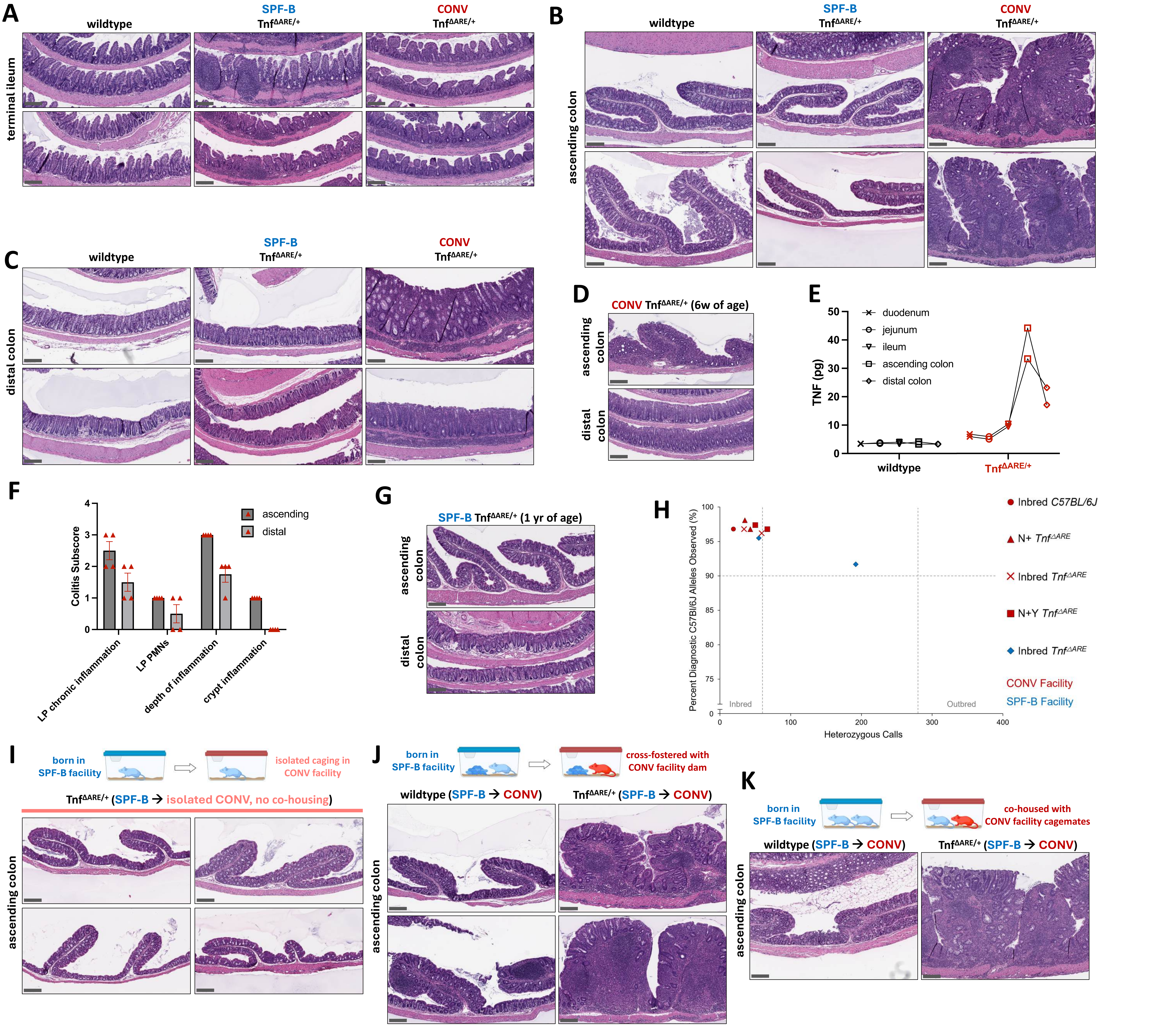


**Figure S1. Genetic background and environmental factors are insufficient to drive colonic inflammation in the Tnf$^{\Delta ARE/+}$ model of Crohn's-like disease.**
(A - C) Additional replicates of H&E-stained intestinal sections (terminal ileum, ascending colon, distal colon) from wildtype (N = 3) and Tnf$^{\Delta ARE/+}$ mice from SPF-B (N = 4) and CONV facilities (N = 5). Wildtype samples are from the CONV facility and all mice are age-matched (34-42 w of age).
(D) Representative H&E-stained colonic sections from a 6w old CONV Tnf$^{\Delta ARE/+}$ mouse.
(E) TNF protein levels measured by ELISA in each region of the small intestine and colon from wildtype (N = 2) and Tnf$^{\Delta ARE/+}$ (N = 2) mice, aged 26-38w.
(F) Colitis subscores that contribute to overall colitis score from histopathological scoring of colons from age-matched CONV Tnf$^{\Delta ARE/+}$ mice (N = 4, 12w of age), separated by ascending and distal colon regions. Mean plus SEM are shown, and statistical significance was not determined. LP = lamina propria. PMNs = polymorphonuclear leukocytes.
(G) Representative H&E-stained colonic sections from a 1 year-old SPF-B Tnf$^{\Delta ARE/+}$ mouse.
(H) Comparison of background amongst SPF-B and CONV facility mice from various indicated generations of backcrossing and inbreeding using MiniMUGA genetic analysis. "N+" indicates more than one backcross to C57BL/6J and "N+Y" indicates the more than one backcross to C57BL/6J, one of which was male to refresh the Y chromosome.
(I) H&E-stained ascending colon sections from SPF-B Tnf$^{\Delta ARE/+}$ mice (N = 4, 27w of age at collection) transferred as adults and housed in isolated caging in the CONV facility.
(J) Additional replicates of H&E-stained ascending colon sections from SPF-B wildtype Tnf$^{\Delta ARE/+}$ mice transferred and co-housed/fostered as pups in the CONV facility with a wildtype or Tnf$^{\Delta ARE/+}$ foster dam along with the dam's age-matched biological pups. Pups were transferred and fostered within the first 3 days of birth and co-housed until experimental collection (37w of age, N = 3 wildtype, N = 4 Tnf$^{\Delta ARE/+}$).
(K) Additional replicates of H&E-stained ascending colon sections from SPF-B adult mice (N = 2 wildtype, N = 3 Tnf$^{\Delta ARE/+}$) transferred and co-housed in the CONV facility in mixed-sex conditions until experimental collection (32-54w of age). CONV donors are of either wildtype or Tnf$^{\Delta ARE/+}$ genotype.
Scale bars = 200 μm.
**Related to Figure 1**.

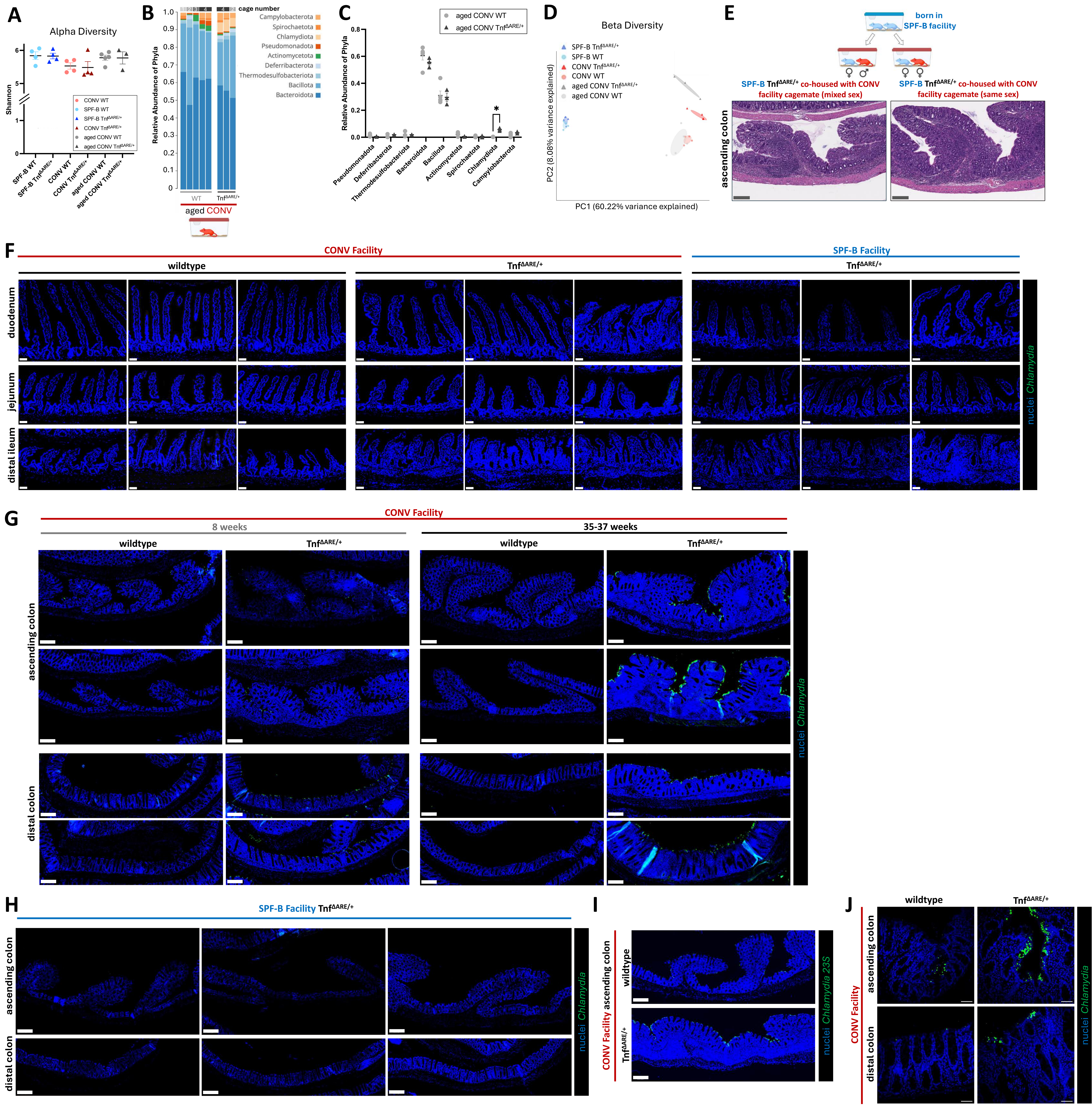


**Figure S2. Shotgun metagenomic sequencing and *in situ* imaging of *Chlamydia muridarum* in wildtype and $Tnf^{\Delta ARE/+}$ intestinal specimens.**

(A) Alpha diversity, measured as the Shannon index, of individual proximal colon luminal content shotgun metagenomic datasets at the species level (filtered for Eubacteria kingdom only). N = 4 per condition. Mean plus standard error of the mean (SEM) are shown and statistical significance was determined using ordinary one-way ANOVA with multiple comparisons.

(B-C) Shotgun metagenomic data of proximal (ascending) colon luminal contents from aged (20-24w of age) wildtype (N = 4) and $Tnf^{\Delta ARE/+}$ (N = 3) mice from the CONV facility from 4 cages. Data is represented as relative abundance of mapped phyla for individual wildtype and $Tnf^{\Delta ARE/+}$ mice in the CONV facility. Mean plus standard error of the mean (SEM) are shown and statistical significance was determined using multiple unpaired t tests with false discovery rate (FDR) of 1%.

(D) Beta diversity at the species level amongst the same samples. N = 4 per condition.

(E) Representative H&E-stained ascending colon sections from SPF-B adult $Tnf^{\Delta ARE/+}$ mice (N = 2 same-sex, N = 4 mixed-sex) transferred and co-housed in the CONV facility in same-sex or mixed-sex conditions until experimental endpoint (15-24w of age). Scale bars = 200 µm.

(F) IF images of *Chlamydia* major outer membrane protein (MOMP - green) and nuclei (Hoechst - blue) co-staining in small intestinal sections. Representative images from duodenum, jejunum, and terminal/distal ileum regions of CONV wildtype (N = 3), CONV $Tnf^{\Delta ARE/+}$ (N = 3), and SPF-B $Tnf^{\Delta ARE/+}$ (N = 3) mice are shown. Scale bars = 50 µm.

(G) Additional replicates of IF images of *Chlamydia* major outer membrane protein (MOMP - green) and nuclei (Hoechst - blue) co-staining on colonic sections from wildtype (N = 3 at 8w of age, N = 4 at 35-37w of age) and $Tnf^{\Delta ARE/+}$ (N = 3 at 8w of age, N = 3 at 35-37w of age) mice from the CONV facility. Scale bars = 200 µm.

(H) IF images of *Chlamydia* major outer membrane protein (MOMP - green) and nuclei (Hoechst - blue) co-staining on colonic sections from aged $Tnf^{\Delta ARE/+}$ (N = 3 at 35-37w of age) mice from the SPF-B facility. Scale bars = 200 µm.

(I) Representative images of fluorescence *in situ* hybridization of *C. muridarum* 23S RNA (green) with nuclei (Hoechst – blue) co-staining in age-matched (13w) wildtype and $Tnf^{\Delta ARE/+}$ mice from the CONV facility. Scale bars = 200 µm.

(J) Representative IF confocal microscopy images of *Chlamydia* major outer membrane protein (MOMP - green) and nuclei (Hoechst - blue) co-staining of the ascending and distal colon. N = 3 per condition. Scale bars = 50 µm.

p-value * < 0.05, ** < 0.01, *** < 0.001. **** < 0.0001.

**Related to Figure 2 and Table S1.**

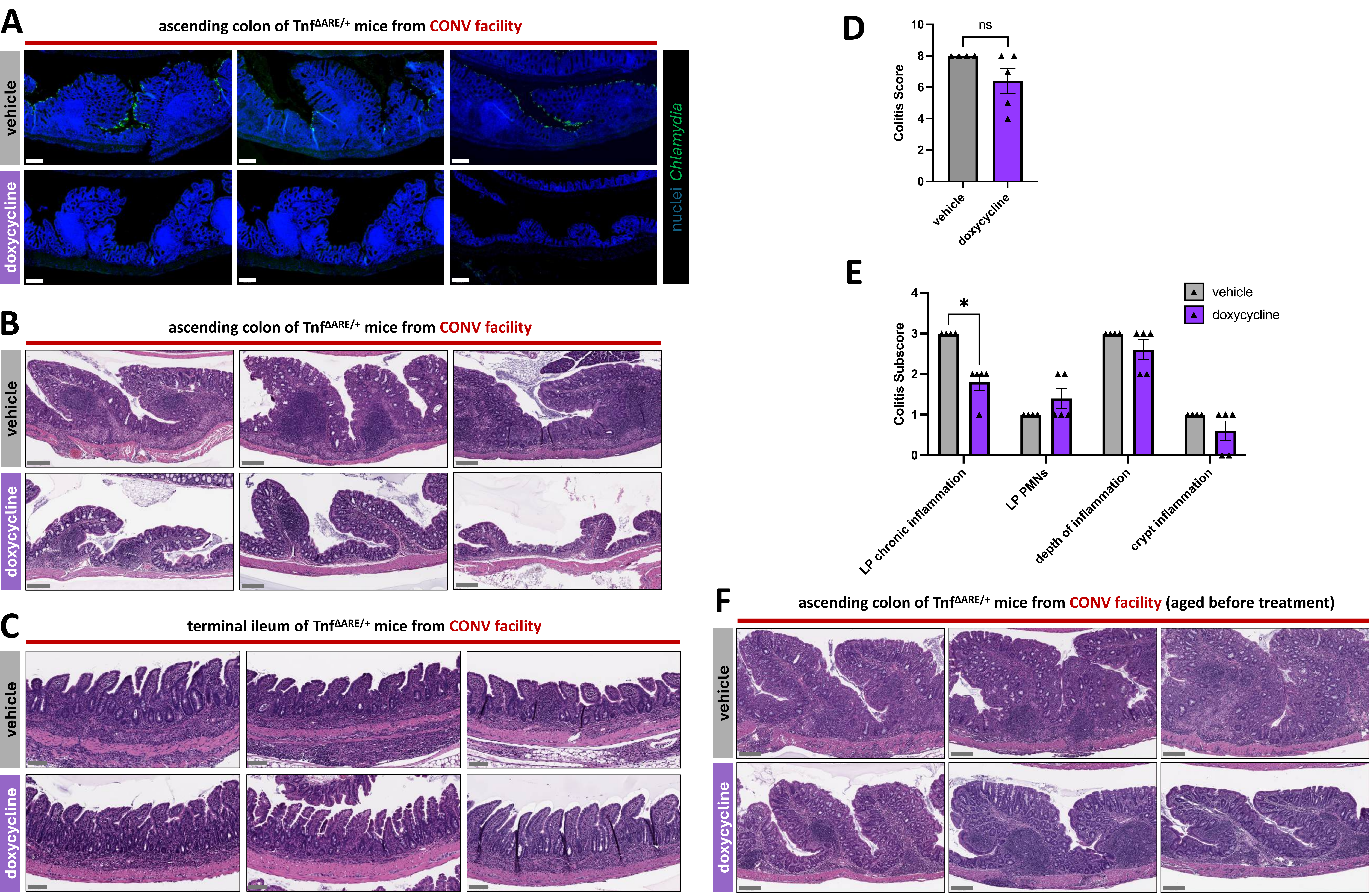


**Figure S3. Doxycycline treatment clears *Chlamydia muridarum* infection and resolves inflammation in the colon of Tnf$^{\Delta ARE/+}$ mice.**

(A) Additional replicates of IF images of *Chlamydia* major outer membrane protein (MOMP - green) and nuclei (Hoechst - blue) co-staining on ascending colon sections from CONV Tnf$^{\Delta ARE/+}$ mice treated with doxycycline or vehicle. N = 4 mice per condition, age-matched at 11-12w of age at harvest. Scale bars = 200 µm.

(B) Additional replicates of H&E-stained ascending colon sections from from CONV Tnf$^{\Delta ARE/+}$ mice treated with doxycycline or vehicle. N = 4 mice per condition, age-matched at 11-12w of age at harvest. Scale bars = 200 µm.

(C) H&E-stained terminal ileum sections from from CONV Tnf$^{\Delta ARE/+}$ mice treated with doxycycline or vehicle. N = 4 mice per condition, age-matched at 11-12w of age at harvest. Scale bars = 100 µm.

(D) Colitis scores from histopathological scoring of colons from aged CONV Tnf$^{\Delta ARE/+}$ mice (13-16w at start, 18-23w at harvest) treated with doxycycline or vehicle. N = 4 mice per condition. Mean plus SEM are shown, and statistical significance was determined using an unpaired t test.

(E) Colitis subscores that contribute to overall colitis score in D. Mean plus SEM are shown, and statistical significance was determined using multiple unpaired t tests with FDR of 1%. LP = lamina propria. PMNs = polymorphonuclear leukocytes.

(F) H&E-stained ascending colon sections from aged CONV Tnf$^{\Delta ARE/+}$ mice (13-16w at start, 18-23w at harvest) treated with doxycycline or vehicle. N = 4 mice per condition. Scale bars = 200 µm.

p-value * < 0.05, ** < 0.01, *** < 0.001. **** < 0.0001.

**Related to Figure 3.**

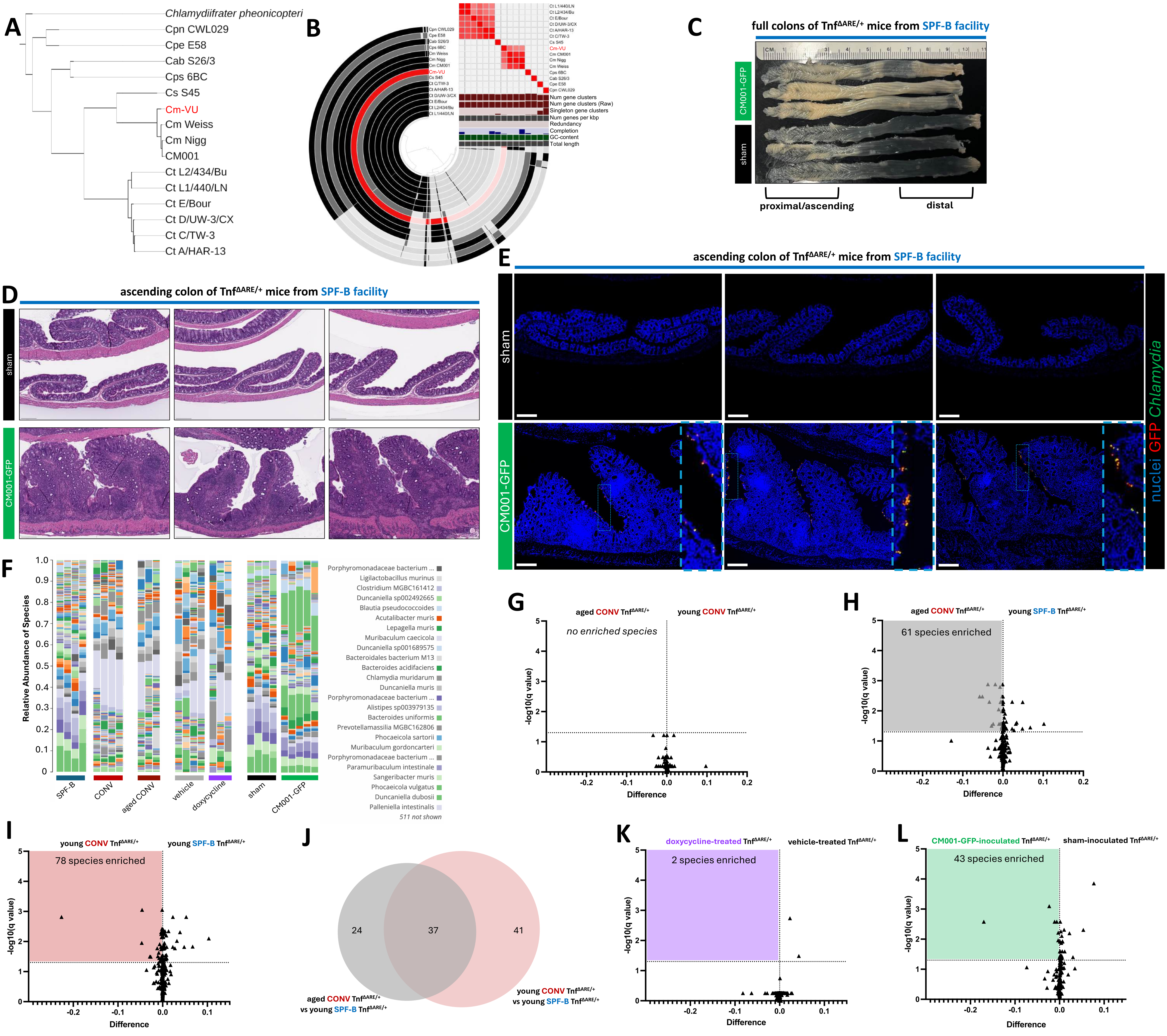


**Figure S4. *Chlamydia muridarum* strain CM001-GFP is sufficient to drive colonic inflammation in Tnf$^{\Delta ARE/+}$ mice.**

(A) Pairwise analysis of genome similarity identifies Cm-VU as *C. muridarum*. ANI percentage identity was calculated using pyANI within anvi'o and the resulting dendrogram was visualized to group *Chlamydia* genomes. *Chlamydiifrater phoenicopterid* was used as an outgroup. Notably, Cm-VU (red) is grouped with the *C. muridarum* genomes.

(B) Pangenomic analysis of *Chlamydia* genomes. The phylogram is clustered based on full ANI percentage identity, alternating species are colored in black and grey, and Cm-VU is highlighted in red. The heatmap (top-right) displays ANI similarity between genomes. Shades of red denote pairwise ANI of 95% or greater.

(C) Macroscopic images of whole colons from Tnf$^{\Delta ARE/+}$ mice that are sham or CM001-GFP-inoculated. 3 representative colons from N = 10 mice per condition, age-matched at 16-20w of age at harvest.

(D) Additional replicates of H&E-stained ascending colon sections from from Tnf$^{\Delta ARE/+}$ mice that are sham or CM001-GFP-inoculated. N = 10 mice per condition, age-matched at 16-20w of age at harvest.

(E) Additional replicates of IF images of *Chlamydia* major outer membrane protein (MOMP - green), endogenous GFP from CM001-GFP (GFP – red), and nuclei (Hoechst - blue) co-staining on ascending colon sections from Tnf$^{\Delta ARE/+}$ mice that are sham or CM001-GFP-inoculated. N = 5 mice per condition, age-matched at 16-20w of age at harvest. Inset to show colocalization of GFP and MOMP signal in CM001-GFP-inoculated conditions.

(F) Shotgun metagenomic data, filtered for only the eubacteria kingdom and represented as relative abundance at the species level, of proximal (ascending) colon luminal contents from individual Tnf$^{\Delta ARE/+}$ mice in the experimental paradigms in Figure 2A (young CONV facility), Supp 2C (aged CONV facility), Figure 3G (vehicle vs doxycycline-treated), and Figure 3P (sham vs CM001-GFP-inoculated). Highest abundance species names are shown.

(G) Statistically enriched species in aged CONV (20-24w of age) vs young CONV (6-7 w of age) Tnf$^{\Delta ARE/+}$ mice derived from shotgun metagenomic data. Statistical enrichment was determined using multiple unpaired t tests with FDR of 5%, which resulted in 0 hits.

(H) Statistically enriched species in aged CONV (20-24w of age) vs young SPF-B (6-7 w of age) Tnf$^{\Delta ARE/+}$ mice derived from shotgun metagenomic data. Statistical enrichment was determined using multiple unpaired t tests with FDR of 5%, which resulted in 61 hits.

(I) Statistically enriched species in young CONV (6-7w of age) vs young SPF-B (6-7 w of age) Tnf$^{\Delta ARE/+}$ mice derived from shotgun metagenomic data. Statistical enrichment was determined using multiple unpaired t tests with FDR of 5%, which resulted in 78 hits.

(J) Venn diagram depicting enriched species that are shared between young CONV and aged CONV Tnf$^{\Delta ARE/+}$ mice when compared to young SPF-B mice.

(K) Statistically enriched species in doxycycline-treated vs vehicle-treated Tnf$^{\Delta ARE/+}$ mice from the CONV facility of mixed ages derived from shotgun metagenomic data. Statistical enrichment was determined using multiple unpaired t tests with FDR of 5%, which resulted in 2 hits.

(L) Statistically enriched species in CM001-GFP-inoculated vs sham-inoculated Tnf$^{\Delta ARE/+}$ mice from the SPF-B facility of mixed ages derived from shotgun metagenomic data. Statistical enrichment was determined using multiple unpaired t tests with FDR of 5%, which resulted in 43 hits.

All scale bars = 200 μm. p-value * < 0.05, ** < 0.01, *** < 0.001. **** < 0.0001.

**Related to Figure 3, Table S1, Table S2, and Table S3.**

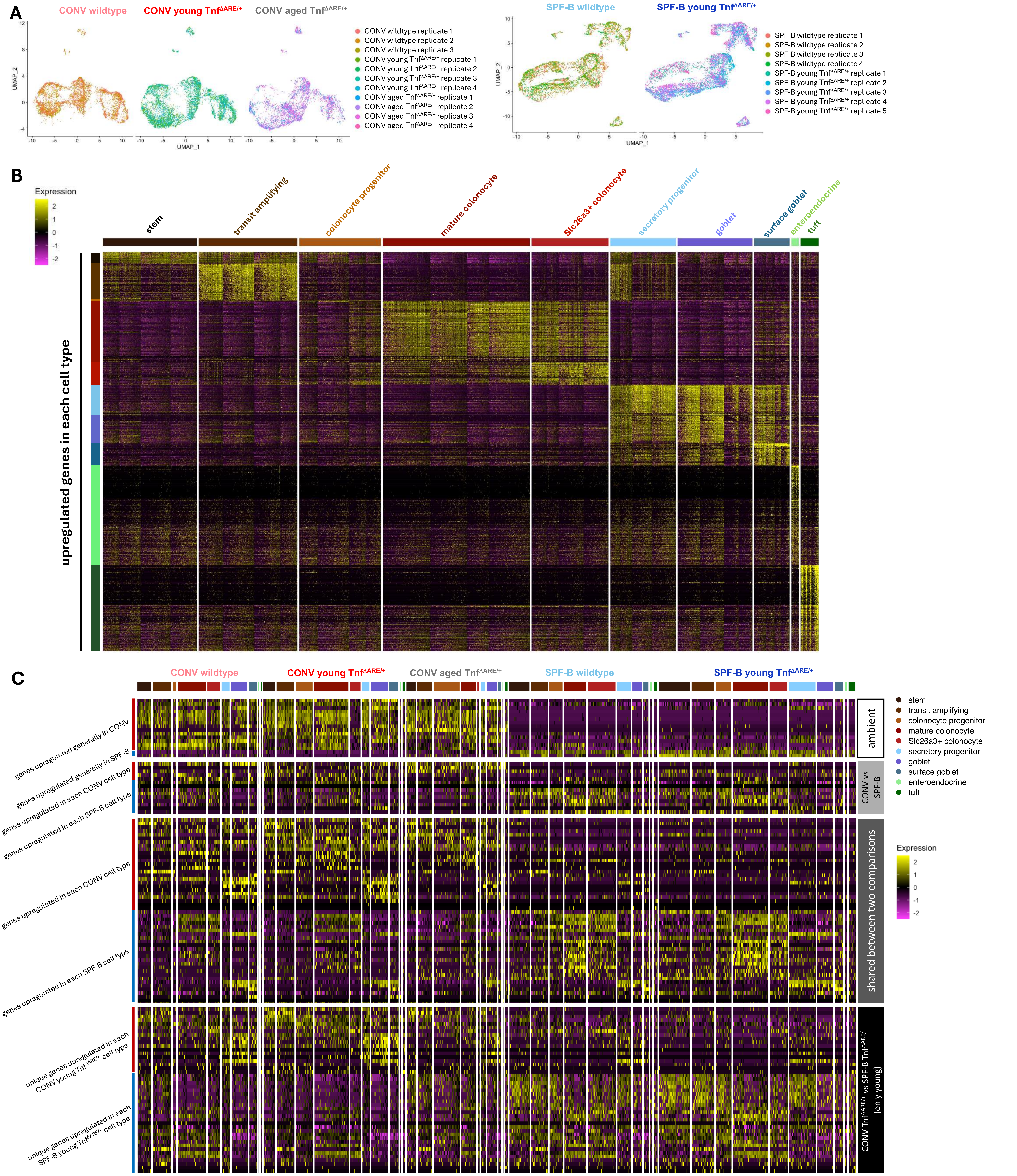


**Figure S5.** ***Chlamydia*** **induces host defense-related transcriptional changes in AC epithelial cell types.**

(A) UMAP co-embedding of scRNA-seq with sample replicate overlay indicated by color.

(B) Heatmap of cell type specific differential gene expression from scRNA-seq data, pooling wildtype and Tnf$^{\Delta ARE/+}$ samples, organized by cell type on x-axis. Genes on y-axis are organized by upregulated genes for each cell type, identified by differential expression analysis of each cell type compared to the rest of the cells. Differentially expressed genes were defined as those with log fold change > 1.

(C) Heatmap of differentially expressed genes from scRNA-seq data, split by cell type and sample type. Genes on the y-axis are organized by generally upregulated (ambient genes) in each facility and those that are differentially expressed in the comparison of CONV vs SPF-B samples from scRNA-seq data (pooled wildtype and Tnf$^{\Delta ARE/+}$ samples), differentially expressed in the comparison of CONV young Tnf$^{\Delta ARE/+}$ vs SPF-B young Tnf$^{\Delta ARE/+}$, and those that are shared between the comparisons. Differentially expressed genes were defined as those with log fold change > 1.5. Generally upregulated genes for each facility were those that were differentially expressed in more than 4 cell types.

**Related to Figure 4, Table S4, Table S5, Table S6, and Table S7.**

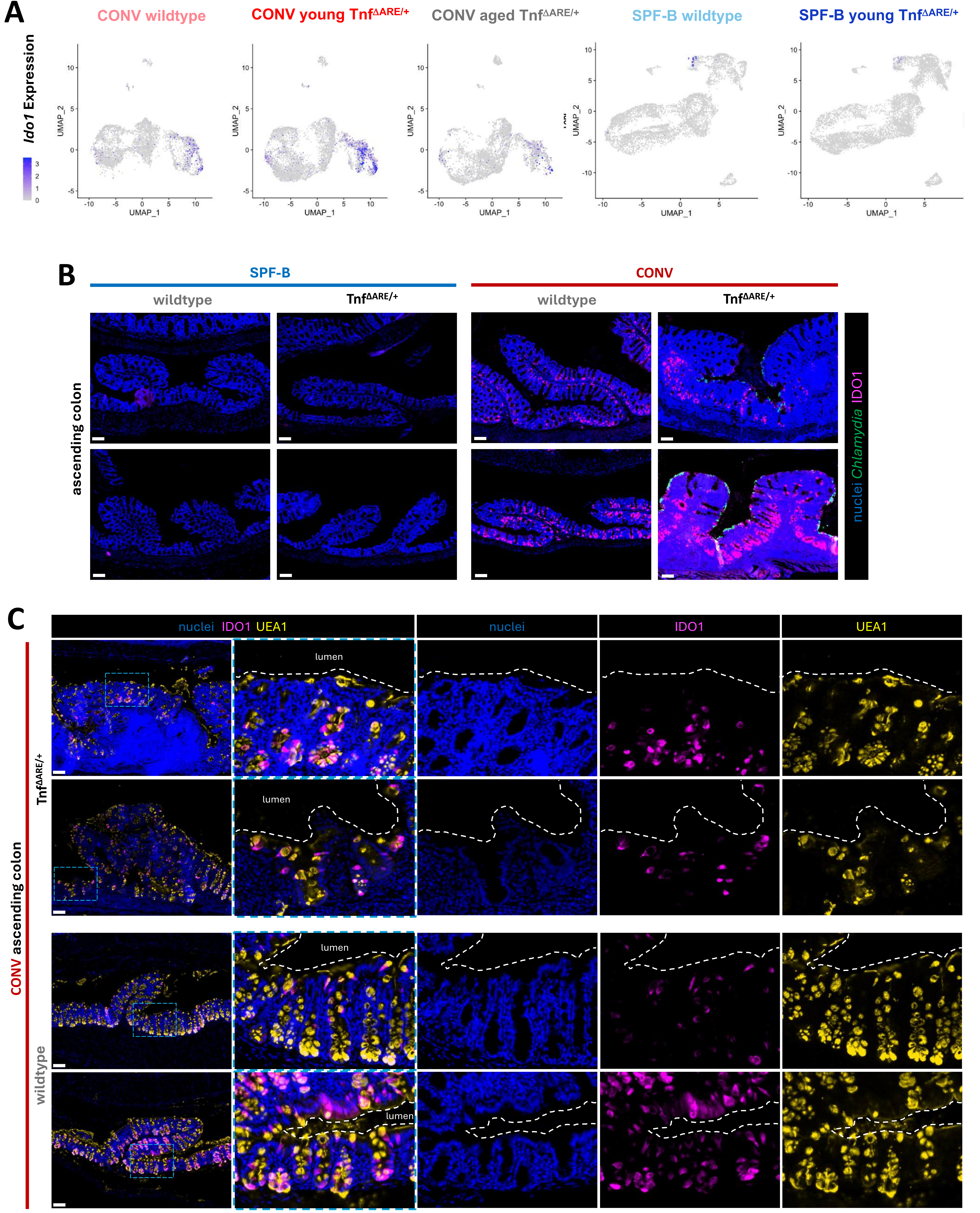


**Figure S6. IDO1 is highly expressed in goblet cells of the ascending colon in *Chlamydia*-colonized wildtype and Tnf$^{ΔARE/+}$ mice.**

(A) UMAP co-embedding of scRNA-seq, separated into individual maps for each sample type, with overlay of *Ido1* gene expression indicated by the color gradient.

(B) Additional replicates of IF images of IDO1 (magenta), *Chlamydia* major outer membrane protein (green), and nuclei (Hoechst - blue) co-staining on ascending colon sections from wildtype and Tnf$^{ΔARE/+}$ mice from the SPF-B and CONV facilities. N = 3 mice, age-matched at 34-42w of age at harvest. Scale bars = 100 µm.

(C) Additional replicates of IF images of IDO1 (magenta), UEA1 lectin (yellow), and nuclei (Hoechst - blue) co-staining on ascending colon sections from wildtype and Tnf$^{ΔARE/+}$ mice from the CONV facilty. Inset image to show colocalization of UEA1 lectin, a goblet and secretory granule marker, with IDO1. N = 3 mice, age-matched at 16-17w of age at harvest. Scale bars = 100 µm.

**Related to Figure 4, Table S5, and Table S6.**

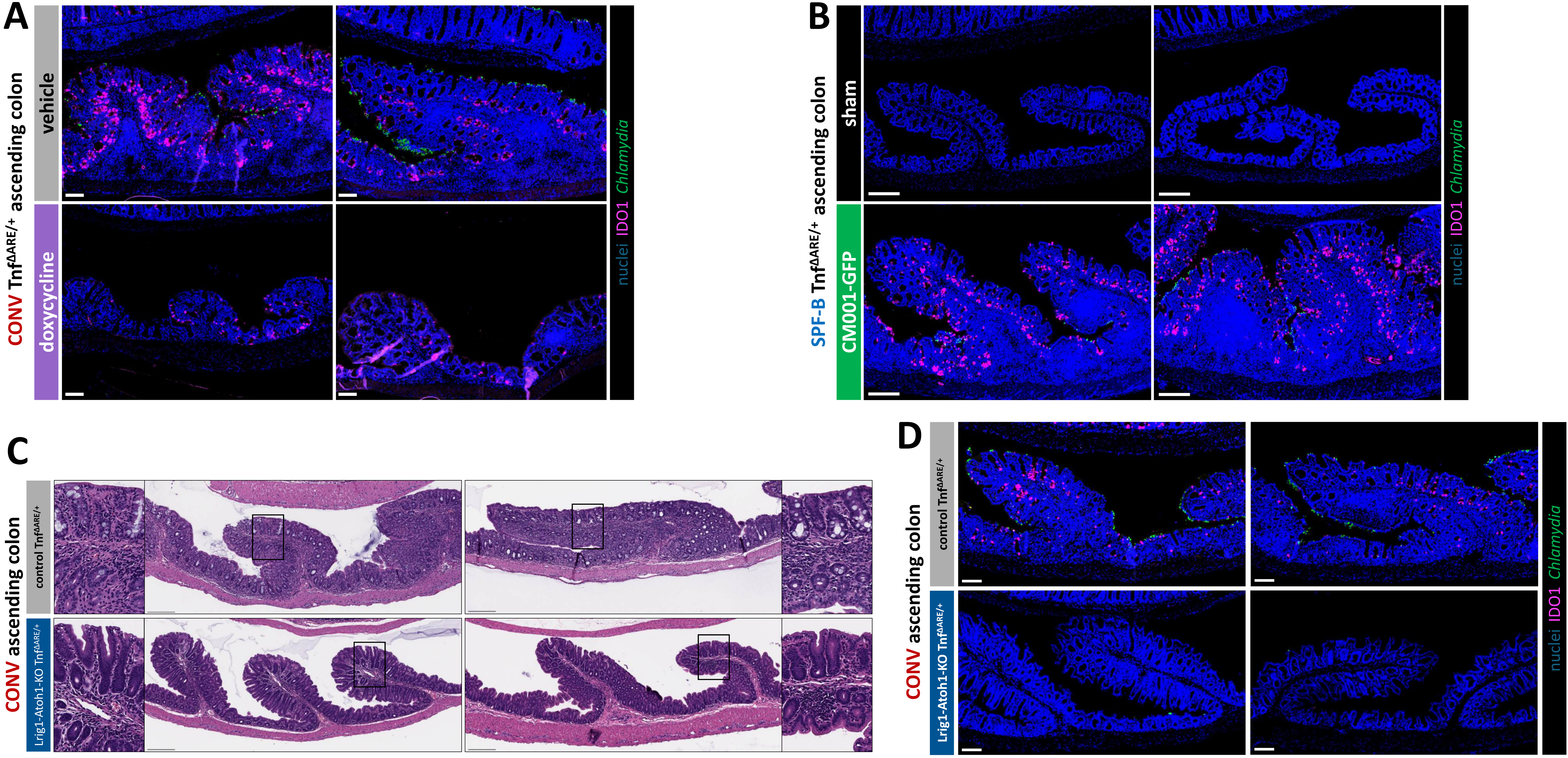


**Figure S7. Secretory cell-derived IDO1 expression is associated with ascending colonic inflammation in the Tnf$^{\Delta ARE/+}$ model.**

(A) Additional replicates of IF images of IDO1 (magenta), *Chlamydia* major outer membrane protein (MOMP - green), and nuclei (Hoechst - blue) co-staining on ascending colon sections from CONV Tnf$^{\Delta ARE/+}$ mice treated with doxycycline or vehicle. N = 4 mice per condition, age-matched at 11-12w of age at harvest. Scale bars = 100 μm.

(B) Additional replicates of IF images of IDO1 (magenta), *Chlamydia* major outer membrane protein (MOMP - green), and nuclei (Hoechst - blue) co-staining on ascending colon sections from SPF-B Tnf$^{\Delta ARE/+}$ mice that are sham or CM001-GFP-inoculated. N = 5 mice per condition, age-matched at 16-20w of age at harvest. Scale bars = 200 μm.

(C) Additional replicates of H&E-stained ascending colon sections from Tnf$^{\Delta ARE/+}$ mice with or without secretory cell ablation. N = 5 control Tnf$^{\Delta ARE/+}$ mice, N = 3 Lrig1-Atoh1-KO Tnf$^{\Delta ARE/+}$ mice, age-matched at 20-27w of age at harvest. Scale bars = 200 μm. Insets to show lack of goblet cell granules in the Lrig1-Atoh1-KO condition.

(D) Additional replicates of IF images of IDO1 (magenta), *Chlamydia* major outer membrane protein (MOMP - green), and nuclei (Hoechst - blue) co-staining on ascending colon sections from Tnf$^{\Delta ARE/+}$ mice with or without secretory cell ablation. N = 5 control Tnf$^{\Delta ARE/+}$ mice, N = 3 Lrig1-Atoh1-KO Tnf$^{\Delta ARE/+}$ mice, age-matched at 20-27w of age at harvest. Scale bars = 100 μm.

**Related to Figure 5.**

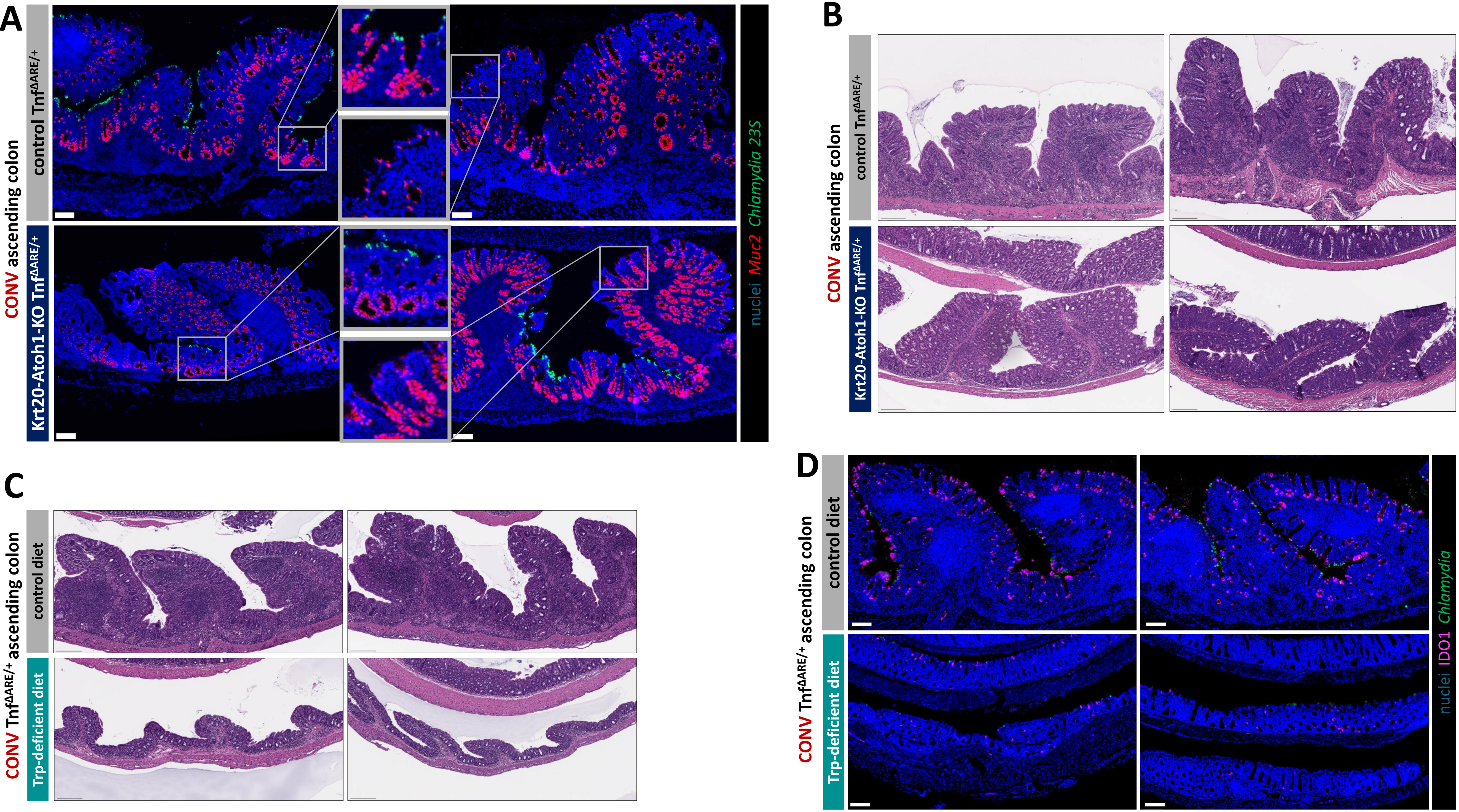


**Figure S8. Goblet cell-derived IDO1 expression is associated with ascending colonic inflammation in the Tnf$^{\Delta ARE/+}$ model.**

(A) Images of fluorescence *in situ* hybridization of *C. muridarum* 23S RNA (green), *Muc2* RNA (red), and nuclei (Hoechst – blue) co-staining in ascending colons of CONV Tnf$^{\Delta ARE/+}$ mice with or without differentiated secretory cell ablation. N = 2 mice per condition, age-matched (13w). Insets to show loss of *Muc2* expression at crypt top in Krt20-Atoh1-KO conditions. Scale bars = 100 µm.

(B) H&E-stained ascending colon sections from CONV Tnf$^{\Delta ARE/+}$ mice with or without differentiated secretory cell ablation. N = 2 mice per condition, age-matched (13w). Scale bars = 200 µm.

(C) Additional replicates of H&E-stained ascending colon sections from CONV Tnf$^{\Delta ARE/+}$ mice fed with control or tryptophan-deficient diet. N = 7 Tnf$^{\Delta ARE/+}$ mice on control diet, N = 5 Tnf$^{\Delta ARE/+}$ mice on tryptophan-deficient diet, age-matched at 11-12w of age at harvest. Scale bars = 200 µm.

(D) Additional replicates IF images of IDO1 (magenta), *Chlamydia* major outer membrane protein (MOMP - green), and nuclei (Hoechst - blue) co-staining on ascending colon sections from CONV Tnf$^{\Delta ARE/+}$ mice fed with control or tryptophan-deficient diet. N = 7 Tnf$^{\Delta ARE/+}$ mice on control diet, N = 5 Tnf$^{\Delta ARE/+}$ mice on tryptophan-deficient diet, age-matched at 11-12w of age at harvest. Scale bars = 100 µm.

**Related to Figure 5.**

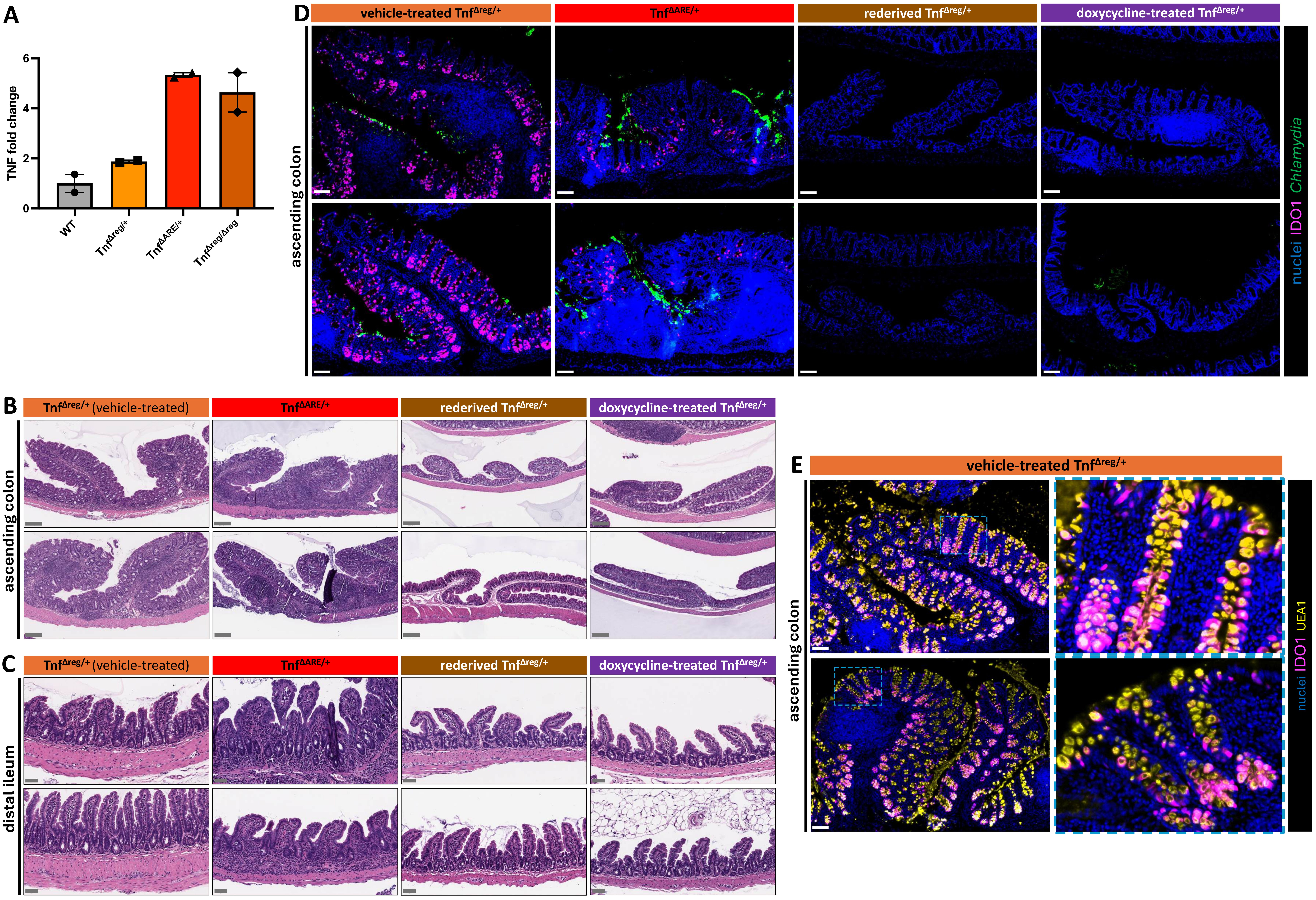


**Figure S9. *Chlamydia* and TNF-driven ascending colon inflammation is not dependent on upstream ileal inflammation.**

(A) TNF protein levels, as fold change relative to wildtype, measured by ELISA from *in vitro* stimulated bone marrow derived macrophages derived from wildtype, Tnf$^{\Delta reg/+}$, Tnf$^{\Delta ARE/+}$, and Tnf$^{\Delta reg/\Delta reg}$ mice. N = 2 mice per condition aged 13-18w at harvest. All mice are from the CONV facility.

(B) Additional replicates for H&E-stained ascending colon sections from CONV Tnf$^{\Delta reg/+}$ (N = 5), CONV Tnf$^{\Delta ARE/+}$ (N = 6), rederived Chlamydia-negative CONV Tnf$^{\Delta reg/+}$ (N = 5), and doxycycline-treated CONV Tnf$^{\Delta ARE/+}$ (N = 4) mice. Mice are age-matched at 16-17w of age at harvest. Scale bars = 200 μm.

(C) Additional replicates for H&E-stained terminal ileum sections from CONV Tnf$^{\Delta reg/+}$ (N = 5), CONV Tnf$^{\Delta ARE/+}$ (N = 6), rederived Chlamydia-negative CONV Tnf$^{\Delta reg/+}$ (N = 5), and doxycycline-treated CONV Tnf$^{\Delta ARE/+}$ (N = 4) mice. Mice are age-matched at 16-17w of age at harvest. Scale bars = 50 μm.

(D) Additional replicates for IF images of IDO1 (magenta), *Chlamydia* major outer membrane protein (MOMP - green), and nuclei (Hoechst - blue) co-staining of ascending colon sections from CONV Tnf$^{\Delta reg/+}$, CONV Tnf$^{\Delta ARE/+}$, rederived Chlamydia-negative CONV Tnf$^{\Delta reg/+}$, and doxycycline-treated CONV Tnf$^{\Delta ARE/+}$ mice. N = 3 mice per condition, age-matched at 16-17w of age at harvest. Scale bars = 100 μm.

(E) Additional replicates for IF images of IDO1 (magenta), UEA1 lectin (yellow), and nuclei (Hoechst - blue) co-staining of ascending colon sections from CONV Tnf$^{\Delta reg/+}$ mice. Inset image to show colocalization of UEA1 lectin, a goblet and secretory granule marker, with IDO1. N = 3 mice, age-matched at 16-17w of age at harvest. Scale bars = 100 μm.

**Related to Figure 6.**

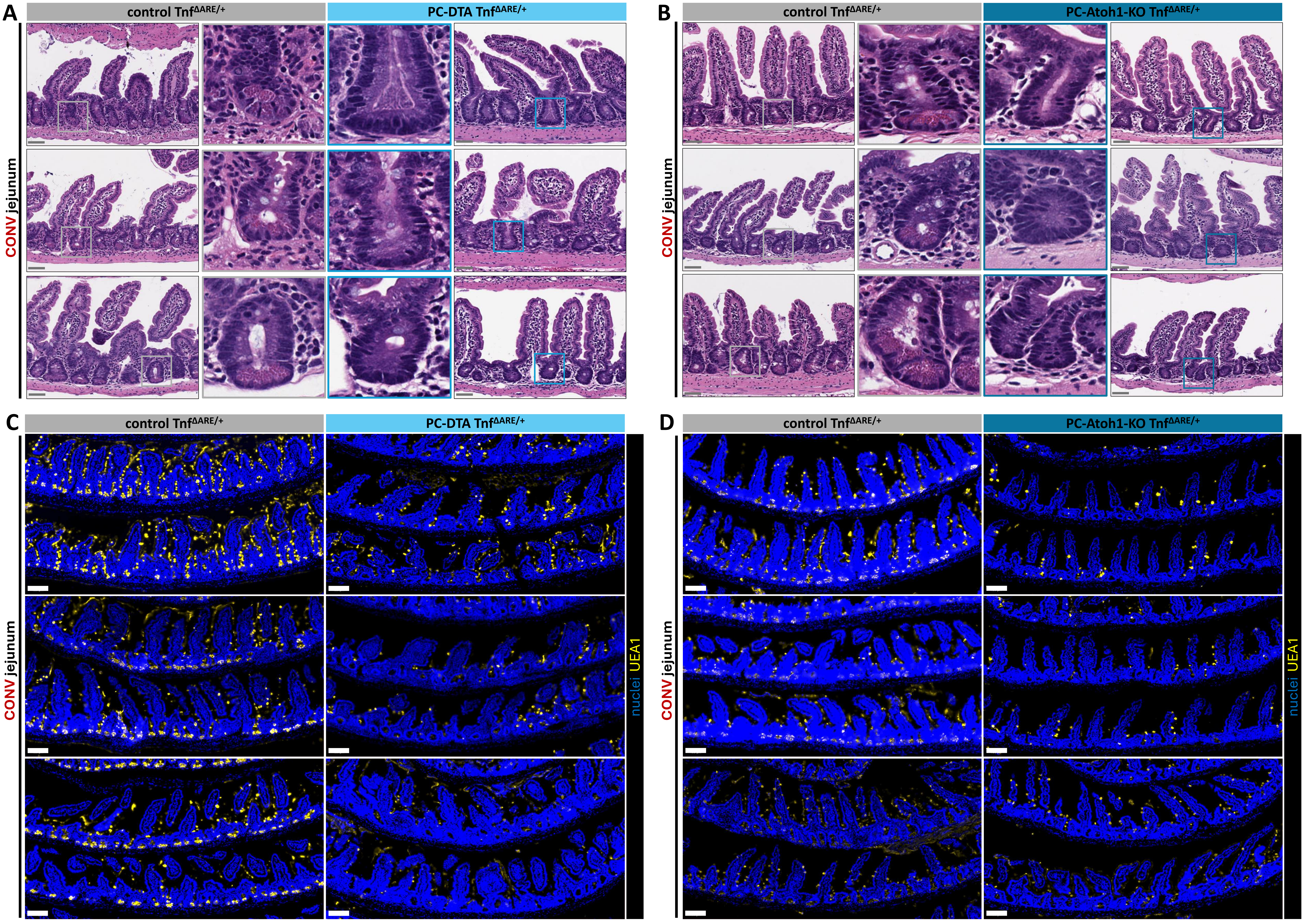


**Figure S10. Paneth cells are depleted from the small intestine of PC-DTA Tnf$^{\Delta ARE/+}$ or PC-Atoh1-KO Tnf$^{\Delta ARE/+}$ mice.**

(A) Representative H&E-stained jejunum sections from control Tnf$^{\Delta ARE/+}$ (N = 4) and PC-DTA Tnf$^{\Delta ARE/+}$ (N = 4) mice. Insets to show lack of Paneth cell granules in crypts in the PC-DTA Tnf$^{\Delta ARE/+}$ condition. Mice are from the CONV facility and are age-matched at 8w of age at harvest. Scale bars = 50 μm.

(B) Representative H&E-stained jejunum sections from control Tnf$^{\Delta ARE/+}$ (N = 3) and PC-Atoh1-KO Tnf$^{\Delta ARE/+}$ (N = 4) mice. Insets to show lack of Paneth cell granules in crypts in the PC-Atoh1-KO Tnf$^{\Delta ARE/+}$ condition. Mice are from the CONV facility and are age-matched at 6-10w of age at harvest. Scale bars = 50 μm.

(C) Representative IF images of UEA1 lectin (yellow) and nuclei (Hoechst - blue) co-staining on jejunum sections from control Tnf$^{\Delta ARE/+}$ (N = 4) and PC-DTA Tnf$^{\Delta ARE/+}$ (N = 4) mice. UEA1 lectin, a Paneth and secretory granule marker, is reduced in the crypts of PC-DTA Tnf$^{\Delta ARE/+}$ jejuna. Mice are from the CONV facility and are age-matched at 8w of age at harvest. Scale bars = 100 μm.

(D) Representative IF images of UEA1 lectin (yellow) and nuclei (Hoechst - blue) co-staining on jejunum sections from control Tnf$^{\Delta ARE/+}$ (N = 4) and PC-Atoh1-KO Tnf$^{\Delta ARE/+}$ (N = 4) mice. UEA1 lectin, a Paneth and secretory granule marker, is reduced in the crypts of PC-Atoh1-KO Tnf$^{\Delta ARE/+}$ jejuna. Mice are from the CONV facility and are age-matched at 6-10w of age at harvest. Scale bars = 100 μm.

**Related to Figure 6.**

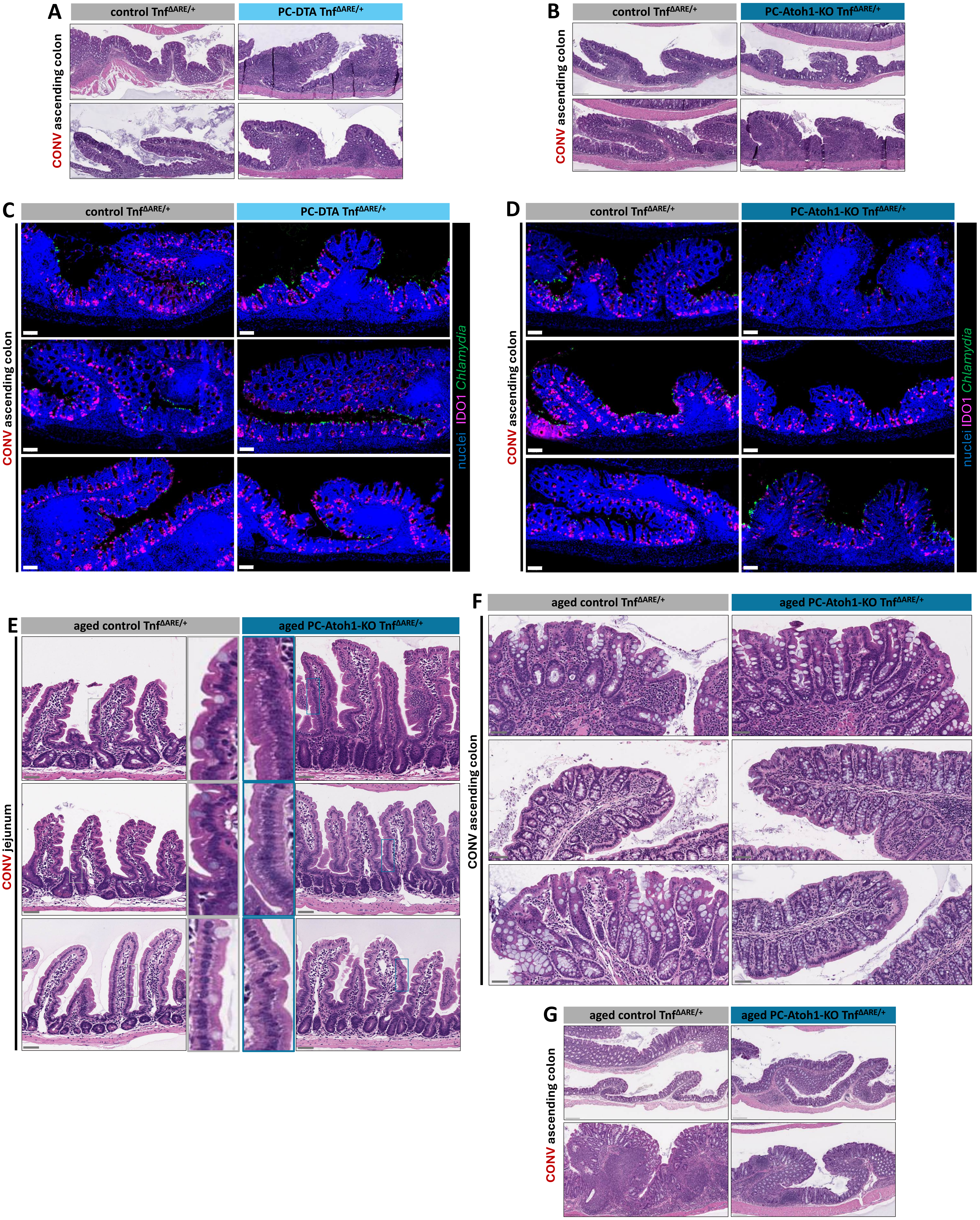


**Figure S11. Paneth cell depletion neither promotes nor protects against ascending colon inflammation in the Tnf$^{\Delta ARE/+}$ model.**

(A) Additional replicates for H&E-stained ascending colon sections from control Tnf$^{\Delta ARE/+}$ (N = 4) and PC-DTA Tnf$^{\Delta ARE/+}$ (N = 4) mice. Mice are from the CONV facility and are age-matched at 8w of age at harvest. Scale bars = 200 μm.

(B) Additional replicates for H&E-stained ascending colon sections from control Tnf$^{\Delta ARE/+}$ (N = 3) and PC-Atoh1-KO Tnf$^{\Delta ARE/+}$ (N = 4) mice. Mice are from the CONV facility and are age-matched at 6-10w of age at harvest. Scale bars = 200 μm.

(C) IF images of IDO1 (magenta), *Chlamydia* major outer membrane protein (MOMP - green), and nuclei (Hoechst - blue) co-staining on ascending colon sections from control Tnf$^{\Delta ARE/+}$ (N = 4) and PC-DTA Tnf$^{\Delta ARE/+}$ (N = 4) mice. Mice are from the CONV facility and are age-matched at 8w of age at harvest. Scale bars = 100 μm.

(D) IF images of IDO1 (magenta), *Chlamydia* major outer membrane protein (MOMP - green), and nuclei (Hoechst - blue) co-staining on ascending colon sections from control Tnf$^{\Delta ARE/+}$ (N = 3) and PC-Atoh1-KO Tnf$^{\Delta ARE/+}$ (N = 4) mice. Mice are from the CONV facility and are age-matched at 6-10w of age at harvest. Scale bars = 100 μm.

(E) Representative H&E-stained jejunum sections from aged control Tnf$^{\Delta ARE/+}$ (N = 4) and aged PC-Atoh1-KO Tnf$^{\Delta ARE/+}$ (N = 4) mice. Mice are from the CONV facility and are age-matched at 23-64w of age at harvest. Scale bars = 50 μm.

(F) Representative H&E-stained ascending colon sections from aged control Tnf$^{\Delta ARE/+}$ (N = 4) and aged PC-Atoh1-KO Tnf$^{\Delta ARE/+}$ (N = 4) mice. Mice are from the CONV facility and are age-matched at 23-64w of age at harvest. Scale bars = 50 μm.

(G) Additional replicates for H&E-stained ascending colon sections from aged control Tnf$^{\Delta ARE/+}$ (N = 4) and aged PC-Atoh1-KO Tnf$^{\Delta ARE/+}$ (N = 4) mice. Mice are from the CONV facility and are age-matched at 23-64w of age at harvest. Scale bars = 200 μm.

**Related to Figure 6.**

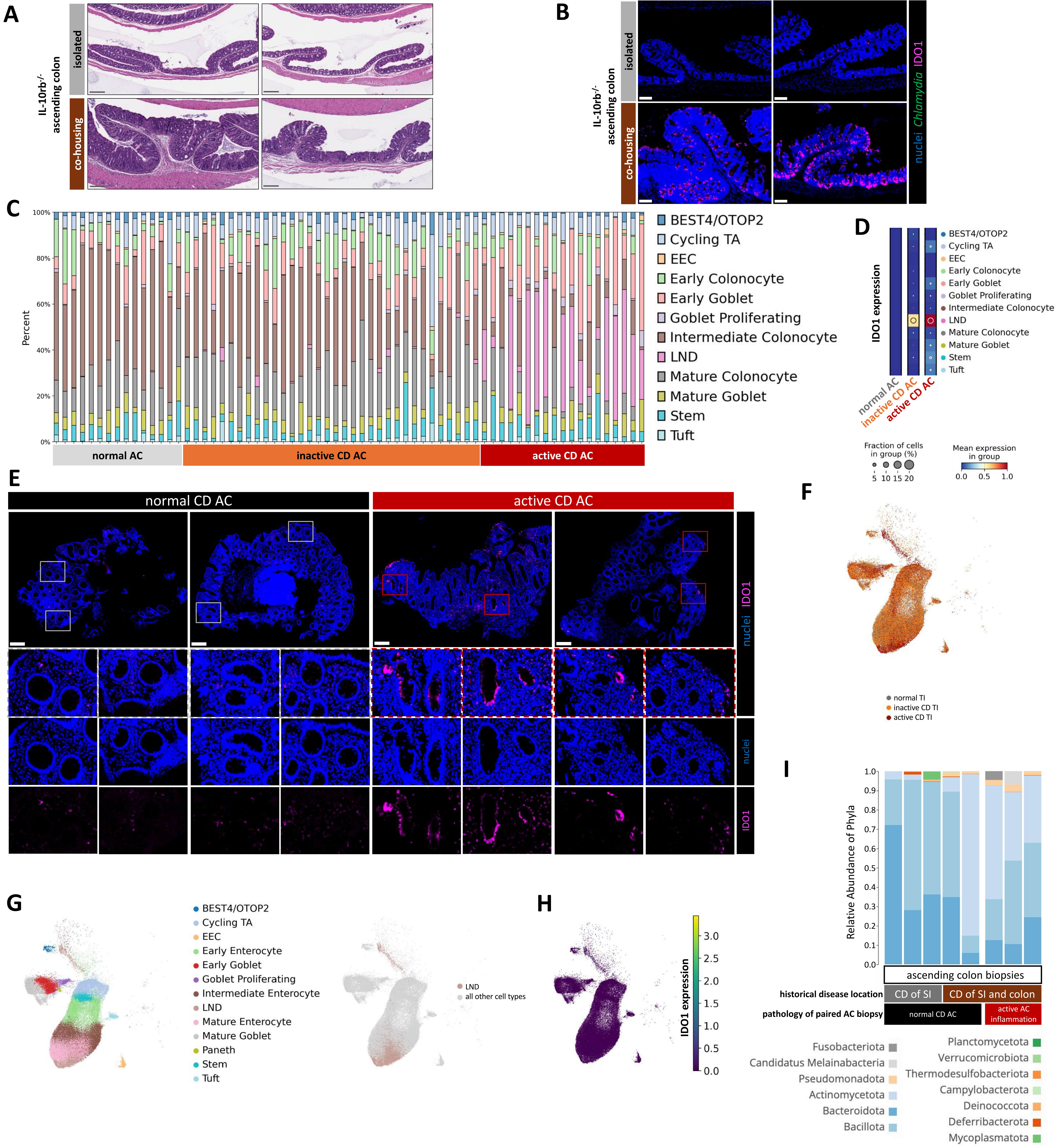


**Figure S12. CD specimens with active AC inflammation are associated with upregulation of IDO1 in epithelial cells and intracellular pathobionts.**

(A) H&E-stained ascending colon sections from IL-10rb$^{-/-}$ mice in isolated caging or *Chlamydia*-positive co-housing conditions. N = 2 mice, age-matched at 38-47w of age at harvest. Co-housing conditions were for 4-8 weeks. Scale bars = 200 µm.

(B) IF images of IDO1 (magenta), *Chlamydia* major outer membrane protein (green), and nuclei (Hoechst - blue) co-staining on ascending colon sections from IL-10rb$^{-/-}$ mice in isolated caging or *Chlamydia*-positive co-housing conditions. N = 2 mice, age-matched at 38-47w of age at harvest. Scale bars = 100 µm.

(C) Barplot with epithelial cell type proportion for each specimen, grouped by sample type, of scRNA-seq data of AC samples.

(D) Dot plot of *IDO1* expression for each cell type, separated by sample type. Percent of cells expressing *IDO1* is indicated by circle size, while expression level is indicated by color gradient.

(E) Additional replicates of IF images of IDO1 (magenta) and nuclei (Hoechst - blue) co-staining on ascending colon biopsies from CD specimens with normal (N = 3) or active (N = 3) histopathological scoring of the AC. Inset and individual channels to show IDO1 expression in epithelial cells. Scale bars = 200 µm.

(F) UMAP co-embedding of scRNA-seq samples of TI epithelial cells from normal (N = 15), inactive CD (N = 30), and active CD (N = 17) specimens. Sample type overlay is indicated by color. Normal TI is composed of healthy control specimens, inactive CD TI is composed of CD specimens histopathologically scored as normal or quiescent, and active CD TI is composed of CD specimens histopathologically scored as mild, moderate, or severe.

(G) UMAP of TI scRNA-seq data with cell type overlay indicated by color (left) and LND cell overlay indicated by color (right).

(H) UMAP of TI scRNA-seq data with overlay of *IDO1* gene expression indicated by the color gradient.

(I) Shotgun metagenomic data, filtered for only the eubacteria kingdom, of AC biopsies from CD specimens with normal (N = 5) or active (N = 3) histopathological scoring. Data is represented as relative abundance of mapped phyla for individual specimens. “CD of SI” indicates AC specimens from CD patients with history of small intestinal inflammation and no history of AC inflammation, while “CD of SI and colon” indicates AC specimens from CD patients with a history of ileocolonic inflammation.

**Related to Fig 7 and Table S8.**
